# Cyclic Electromagnetic DNA Simulation (CEDS) can regulate the Functions of Different miRNAs and DNA Motifs: Protein Signaling Pathways in RAW 264.7 Cells detected by IP-HPLC Analysis

**DOI:** 10.1101/2024.08.23.609321

**Authors:** Yeon Sook Kim, Suk Keun Lee

## Abstract

Based on the hydrogen bonding magnetic resonance (HBMR) in double-stranded DNAs (dsDNAs) exhibiting unique base pair polarities, Cyclic Electromagnetic DNA Simulation (CEDS)^1^ was designed to target nine miRNAs and four protein binding site sequences, which are known to have an anti-oncogenic or oncogenic role in cells. RAW 264.7 cells were cultured and treated with CEDS at 20-25 Gauss in a 5% CO_2_ and 37℃ incubator for 20 min. Besides the cytological observation, the expressions of murine miRNAs and proteins were assessed by quantitative polymerase chain reaction (qPCR) and immunoprecipitation high-performance liquid chromatography (IP-HPLC) using 350 antisera, respectively. The results demonstrated that CEDS was found to increase the number of the corresponding mature and primary miRNAs and influenced the protein signaling in cells. For example, miR-34a-5p*-CEDS was able to downregulate 59 out of 43 target proteins (72.9%), and attenuated RAS-NFkB signaling axis, which subsequently led to the inactivation of proliferation-related signaling axis, protection-ER stress-survival signaling axis, telomere-glycolysis-fibrosis signaling axis, and FAS- and PARP-mediated apoptosis signaling axis. Conversely, miR-34a-5p*-CEDS activated the p53/Rb/E2F1-SHH/PTCH/GLI-Notch/Jagged proliferation signaling axis, epigenetic modification-RAS-p53 associated apoptosis signaling axis, and oncogenesis-senescence-innate immunity-chronic inflammation signaling axis. Among nine CEDS targeting mature miRNAs, miR-150-5p*-CEDSs exhibited a potent anti-oncogenic effect on RAW 264.7 cells, while miR-34a-5p*-, miR-181a-5p*-, miR-216a-5p*-, miR-365a-3p*-, and miR-655-3p*-CEDSs exhibited a dual effect, inducing both anti-oncogenic and oncogenic effects. Conversely, miR-155-5p*-CEDS appears to induce marked oncogenic effect, rather than anti-oncogenic effect. Among four CEDS targeting protein binding site sequences, p53 binding site sequence AGACATGCCT*-CEDS induced an oncogenic effect by upregulating proliferation, telomere, and oncogenesis signaling, while concomitantly inducing an anti-oncogenic effect by downregulating RAS and NFkB signaling and upregulating p53- and PARP-mediated apoptosis. Canonical E2Fs binding site sequence TTTC(C/G)CGC*-CEDS facilitated an oncogenic effect by enhancing RAS and NFkB signaling, resulting in the upregulation of proliferation, telomere, and senescence signaling. Consequently, it was found that CEDS using a miRNA or protein binding site sequence impacted the protein signaling in RAW 264.7 cells in a sequence-specific manner. It is therefore proposed that CEDS can target miRNAs and DNA motifs, influencing their hybridization and conformation, and subsequently regulate the protein signaling in the cells.

## Introduction

In the previous study, it was demonstrated that CEDS has a DNA hybridization potential by increasing the hydrogen bonding strength between base pairs of short dsDNA depending on CEDS time in a sequence-specific manner^1^. CEDS was found to increase the hybridization of oligo-dsDNAs and induce unique conformations of oligo-dsDNAs based on their sequence-specific base pair polarities. Therefore, it is hypothesized that CEDS can affect the hybridization potential of miRNAs, leading to the activation of miRNA biogenesis and function. However, it remains to be seen whether CEDS has a positive effect on miRNA functions in mammalian cells.

MiRNAs are non-coding small RNAs that can interact with other RNAs and DNAs through hybridization. They play a role in post-transcriptional inhibition of protein translation on mRNAs, as well as transcriptional regulation on genomic DNAs^2^. The primary miRNAs undergo partially mismatched hybridization, resulting in the formation of hairpin-like stem loop structures in the miRNA duplex. The mismatched RNA duplexes may exhibit relatively low hybridization strength between the strands of the miRNA duplex, yet they possess high potential for interaction with other RNAs and DNAs. This environment may present a compelling opportunity for the intervention of CEDS.

CEDS was designed to provide electromagnetic stimulation to base pairs in dsDNA in a cyclic manner. The apparatus sequentially produces a magnetic field targeting each base pair of dsDNA in decagonal or dodecagonal fashion at 20-25 Gauss. To achieve optimal electromagnetic induction, each A-T and G-C pair is stimulated with a different exposure time of 280 and 480 msec, respectively. All the methods are digitally programmed in an electric control apparatus. To increase the efficiency of DNA/RNA hybridization, this study employed both strands of target miRNAs and DNA motifs sequences in CEDS, which were designated as “target sequence*-CEDS.” For example, CEDS that utilizes both strands of ds3T3A will be referred to as “3T3A*-CEDS”, while CEDS that uses only one strand of ds3T3A is referred to as “3T3A-CEDS”.

IP-HPLC has been developed to analyze the protein expression levels in comparison with the controls. It utilizes a simple immunoprecipitation method and is followed by the quantitative analysis of HPLC. The IP-HPLC has been validated through a series of experiments involving cultured cells, animal serum, human saliva, and inflammatory exudate^3–8^. It is believed that the IP-HPLC is adequate to detect multiple proteins simultaneously due to its simple, rapid, inexpensive, accurate, and automatic procedures, and furthermore, its results can be analyzed statistically.

The human genome encodes approximately 2,600 mature miRNAs (miRBase V.22), and it is known that a single miRNA can modulate thousands of genes by recognizing complementary sequences not only at the 3’ UTR region but also the 5’ UTR and open reading frame regions. Furthermore, miRNA interaction with promoter and enhancer regions also affects the transcription of associated genes^9,10^. It is estimated that over 60% of all mammalian protein-coding genes may be regulated by miRNAs^11^. The present study demonstrated that when the target genes are post-transcriptionally regulated by CEDS using a target miRNA sequence, the reduced expression of target genes can impact the upstream or downstream proteins in the associated signaling. Therefore, it is postulated that miRNAs can regulate the protein signaling more widely than previously assumed^2^.

In order to ascertain the fidelity of CEDS, the present study conducted multiple CEDSs targeting oncogenesis-related miRNAs, including miR-34a-5p, miR-150-5p, miR-155-5p, 181a-5p, miR-216a-5p, miR-365a-3p, and miR-655-3p. Furthermore, additional CEDSs were performed to target the oncogenesis-related protein binding site sequences, including a p53 cluster 1 response element (RE) binding site sequence (AGACATGCCT) and a canonical E2Fs binding site sequence (TTTC(C/G)CGC) in RAW 264.7 cells. These CEDS targets have been the subject of extensive investigation by numerous authors^12–15^.

The RAW 264.7 cells, derived from monocytes/macrophage-like cells of Balb/c mice, were utilized in this study by limiting their culture passage number to less than 20, as previously described by Taciak^16^. The cells were cultured in Dulbecco’s modified Eagle’s medium (WelGene Inc., Korea) supplemented with 10% fetal bovine serum (WelGene Inc., Korea), 100 units/mL penicillin, 100 μg/mL streptomycin, and 250 ng/mL amphotericin B (WelGene Inc., Korea), in a 5% CO_2_ incubator at 37.5°C. The cell culture product containing 10^8^-10^9^ cells was placed at the center of a horizontal type CEDS in the culture incubator and treated with dodecagonal CEDS using a target miRNA or protein binding site sequence for 20 min. Subsequently, the cells were maintained in the culture incubator without CEDS for 30 min of resting time, and then harvested by centrifugation at 150g for 5 min for the following assays. .

### 1. Quantitative polymerase chain reaction (qPCR) after CEDS targeting different mature and primary miRNAs in RAW 264.7 cells

For quantitative polymerase chain reaction (qPCR) to determine the expression levels of nine miRNAs examined in this study, total miRNAs were extracted from CEDS-treated cells using the miRNeasy kit (Qiagen, USA), and reverse strand cDNAs were generated using the Moloney murine leukemia virus reverse transcriptase (cDNA synthesis kit, Cosmo Genetech, Korea) separately. The cDNAs were then used for qPCR (ChaiBio, USA) with a primer pair targeting an objective miRNA. The unique primer sequences for the mature and primary miRNAs were obtained from the miRBase website by analyzing them with the DNA base pair polarity program (Table 1), and their synthetic primers were purchased from CosmoGenetech Inc. (Seoul, Korea). The qPCR was carried out with varying annealing temperatures (43–53°C) and different PCR cycles (30–45 cycles) to obtain accurate Cq values. This was followed by a DNA melting process to assess the qPCR fidelity. The exponential growth curve and Cq number of the experimental group were compared to those of the untreated control group.

**Table 1.**
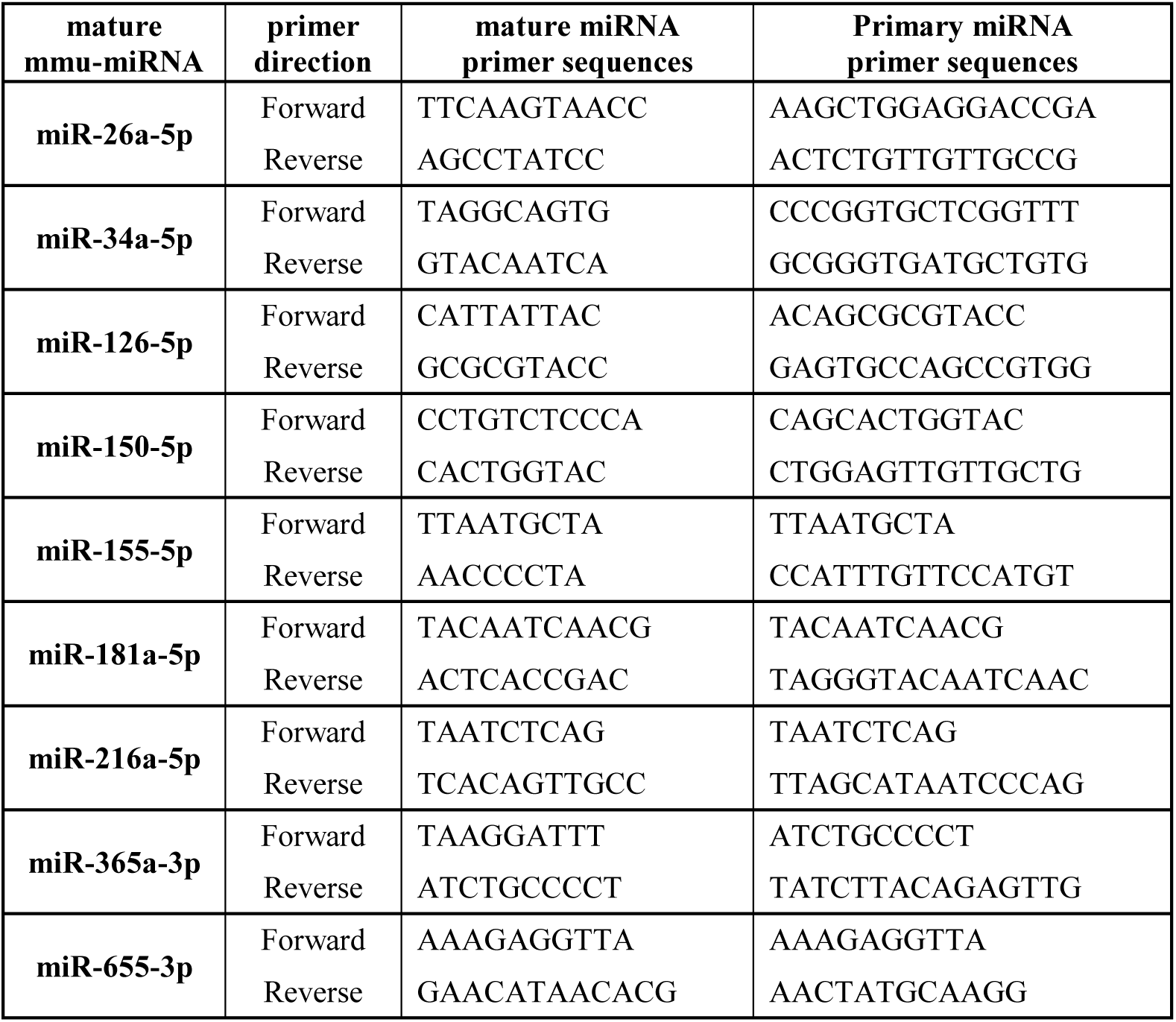
The primer pairs of miRNAs used in quantitative real time PCR (qPCR)

A series of qPCR experiments were conducted using the primer sets and cDNA templates obtained from the cell culture to determine the expression levels of primary and mature miRNAs after CEDS using a target miRNA sequence. The primary aim of this experiment was to assess the targeting efficiency of CEDS for the specific miRNA of interest among different miRNAs present in cells. The qPCRs were simultaneously conducted for 15 miRNAs selected in this study using the template cDNAs obtained from the RAW 264.7 cells treated with CEDS targeting one miRNA among the 15 miRNAs. The second objective of this experiment was to determine the incremental ratio of mature and primary miRNAs in cells following CEDS treatment compared to the untreated controls. The incremental ratio of mature and primary miRNAs was calculated based on the Cq number.

#### qPCR for the expression of mature and primary miRNAs after miR-34a-5p*-CEDS

In the multiple qPCRs for 15 mature miRNAs, mmu-miR-34a-5p sequence TGGCAGTGTCTTAGCTGGTTGT*-CEDS demonstrated a 610% increase in miR-34a-5p expression (Cq=17.34) compared to the untreated control (Cq=14.29) in RAW 264.7 cells. Concurrently, miR-34a-5p*-CEDS was found to significantly increase the expression of miR-126-5p and miR-655-3p compared to the untreated control, while decreasing the expression of miR-21-5p (Fig. 2A and B).

**Fig. 2.**
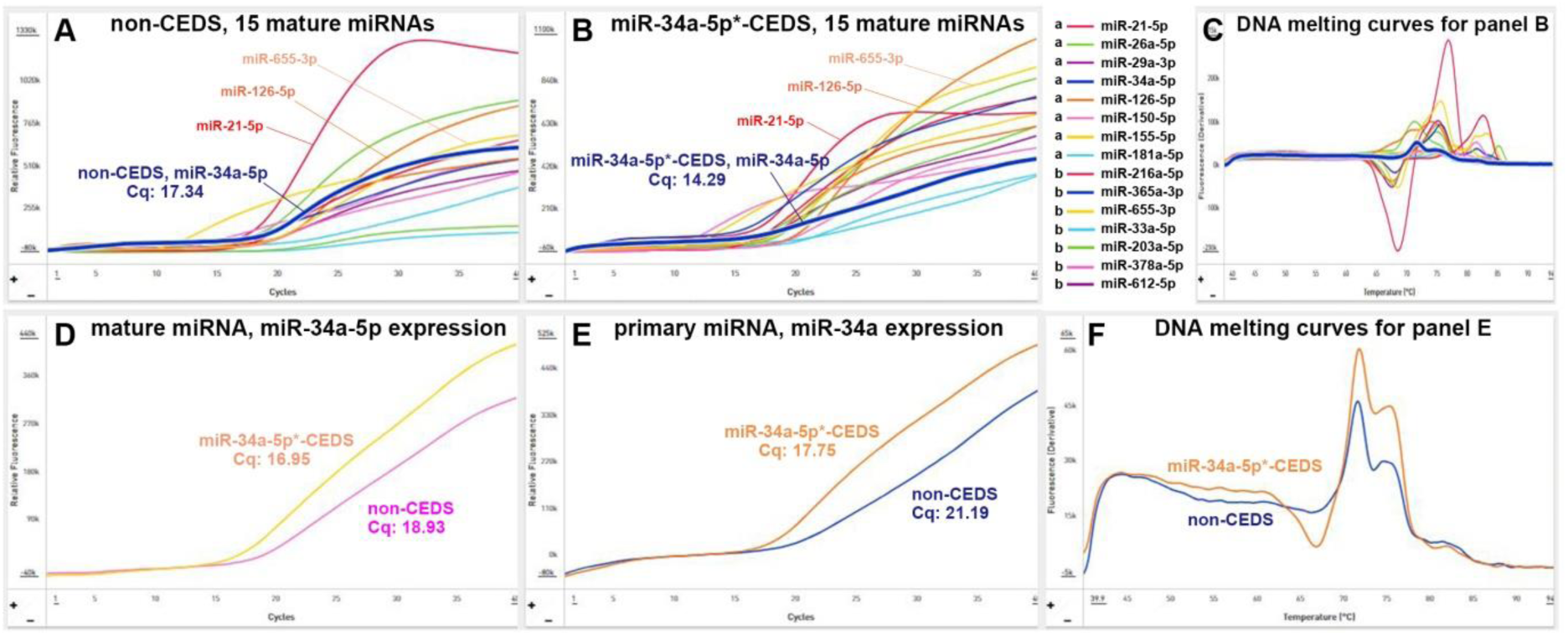
Quantitative PCR for CEDS effect on the expression of mature miR-34a-5p (D) and primary miR-34a (E) in RAW 264.7 cells. MiR-34a-5p*-CEDS increased the expression of miR-34a-5p along with an influence on the expression of other 14 miRNAs (A and B). Noted the variable melting curves of 15 mature miRNAs and similar melting curves of primary miR-34a (C and F). The representative results were presented based on the findings of triple repeated experiments.

In the qPCR assays conducted to determine CEDS-induced incremental ratio of mature and primary miRNAs, miR-34a-5p*-CEDS demonstrated a significant increase in the expression of mature miR-34a-5p and primary miR-34a, with fold changes of 396% and 688%, respectively, in comparison to the untreated controls (Fig. 2 D and E).

#### qPCR for the expression of mature and primary miRNAs after miR-150-5p*-CEDS

In the multiple qPCRs for 15 mature miRNAs, mmu-miR-150-5p sequence TCTCCCAACCCTTGTACCAGTG*-CEDS demonstrated a 1080% increase in miR-150-5p expression (Cq=14.94) expression compared to the untreated control (Cq=9.54) in RAW 264.7 cells. Concurrently, miR-150-5p*-CEDS was found to significantly increase the expression of miR-126-5p, miR-365a-3p, and miR-655-3p compared to the untreated control (Fig. 4 A and B).

**Fig. 4.**
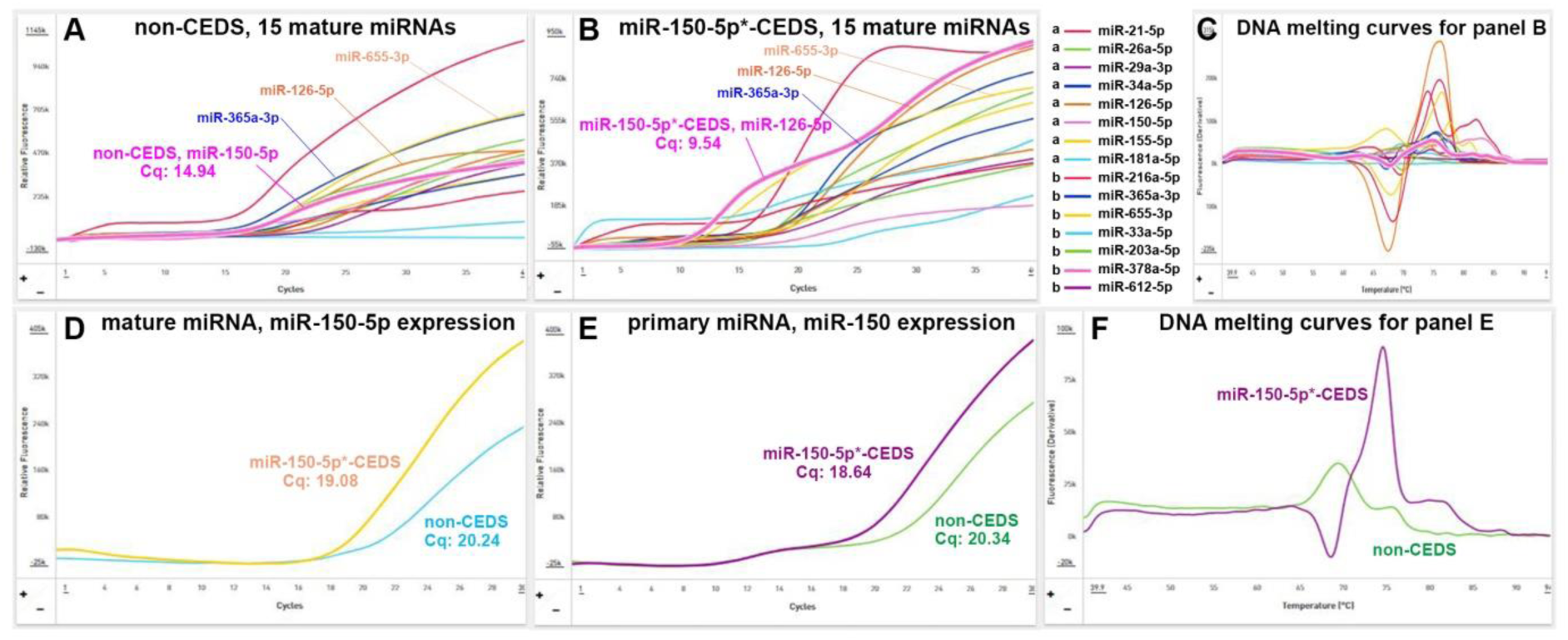
Quantitative PCR for CEDS effect on the expression of mature miR-150-5p (D) and primary miR-150 (E) in RAW 264.7 cells. MiR-150-5p*-CEDS increased the expression of miR-150-5p along with an influence on the expression of other 14 miRNAs (A and B). Noted the variable melting curves of 15 mature miRNAs and similar melting curves of primary miR-150 (C and F). The representative results were presented based on the findings of triple repeated experiments.

In the qPCR assays conducted to determine CEDS-induced incremental ratio of mature and primary miRNAs, miR-150-5p*-CEDS demonstrated a significant increase in the expression of mature miR150-5p and primary miR-150, with fold changes of 232% and 340%, respectively, in comparison to the untreated controls (Fig. 4 D and E).

#### qPCR for the expression of mature and primary miRNAs after miR-155-5p*-CEDS

In the multiple qPCRs for 15 mature miRNAs, mmu-miR-155-5p sequence TTAATGCTAATCGTGATAGGGGTT*-CEDS demonstrated a 924% increase in miR-155-5p expression (Cq=15.83) expression compared to the untreated control (Cq=11.21) in RAW 264.7 cells. Concurrently, miR-155-5p*-CEDS was found to significantly increase the expression of miR-181a-5p, miR-216a-5p, and miR-655-3p compared to the untreated control (Fig. 5 A and B).

**Fig. 5.**
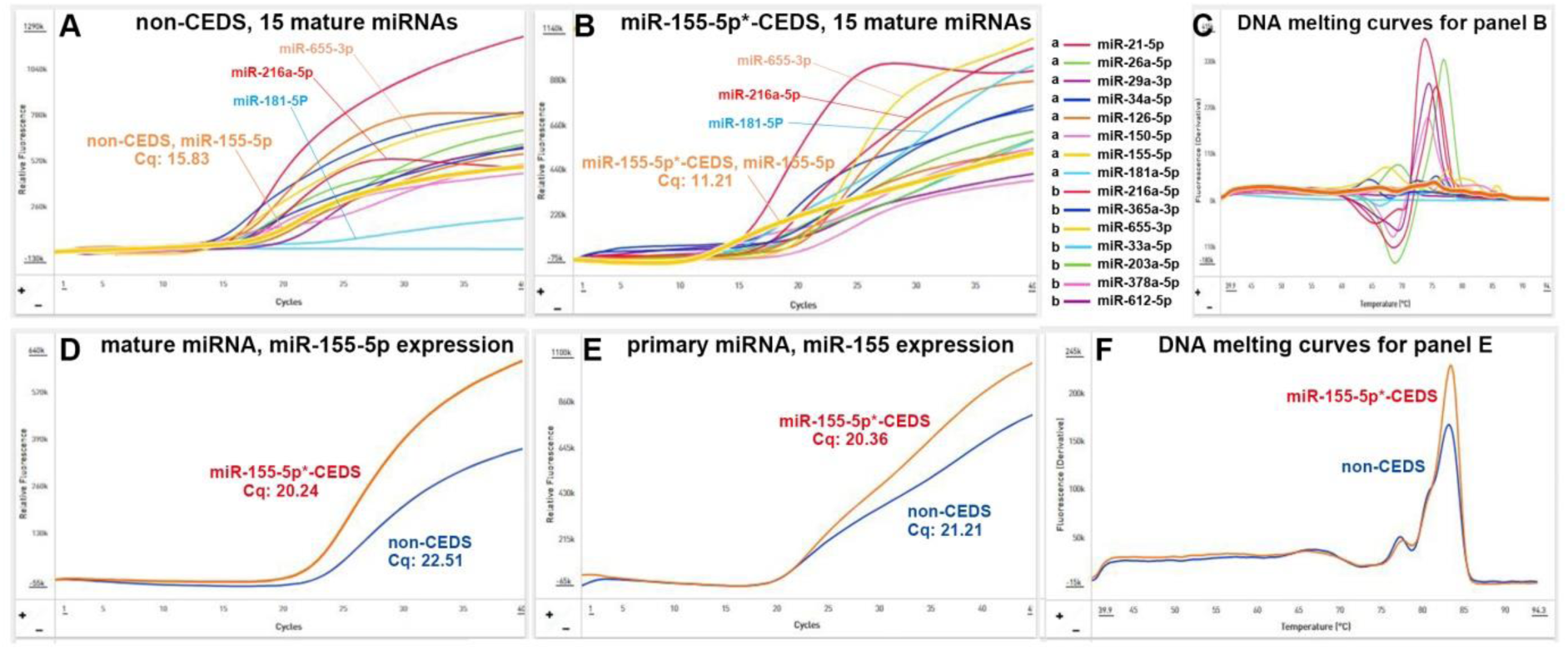
Quantitative PCR for CEDS effect on the expression of mature miR-155-5p (D) and primary miR-155 (E) in RAW 264.7 cells. MiR-155-5p*-CEDS increased the expression of miR-155-5p along with an influence on the expression of other 14 miRNAs (A and B). Noted the variable melting curves of 15 mature miRNAs and similar melting curves of primary miR-155 (C and F). The representative results were presented based on the findings of triple repeated experiments.

In the qPCR assays conducted to determine CEDS-induced incremental ratio of mature and primary miRNAs, miR-155-5p*-CEDS demonstrated a significant increase in the expression of mature miR-155-5p and primary miR-155, with fold changes of 454% and 170%, respectively, in comparison to the untreated controls (Fig. 5 D and E).

#### qPCR for the expression of mature and primary miRNAs after miR-181a-5p*-CEDS

In the multiple qPCRs for 15 mature miRNAs, mmu-miR-181a-5p*-CEDS AACATTCAACGCTGTCGGTGAGT*-CEDS demonstrated a 456% increase in miR-181a-5p expression (Cq=16.1) compared to the untreated control (Cq=14.12) in RAW 264.7 cells. Concurrently, miR-181a-5p*-CEDS was found to significantly increase the expression of miR-126-5p, miR-216a-5p, and miR-655-3p compared to the untreated control (Fig. 6 A and B).

**Fig. 6.**
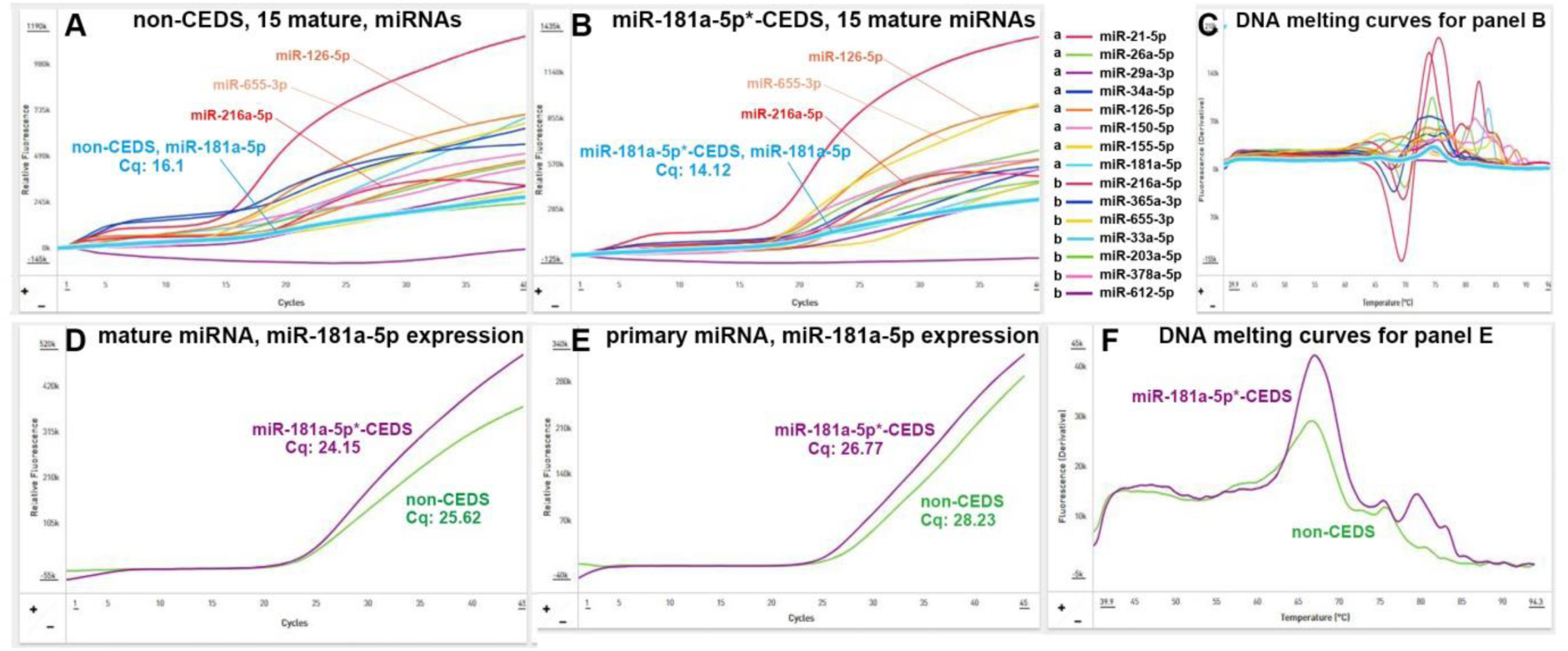
Quantitative PCR for CEDS effect on the expression of mature miR-181a-5p (D) and primary miR-181a (E) in RAW 264.7 cells. MiR-181a-5p*-CEDS increased the expression of miR-181a-5p along with an influence on the expression of other 14 miRNAs (A and B). Noted the variable melting curves of 15 mature miRNAs and similar melting curves of primary miR-181a (C and F). The representative results were presented based on the findings of triple repeated experiments.

In the qPCR assays conducted to determine CEDS-induced incremental ratio of mature and primary miRNAs, miR-181a-5p*-CEDS demonstrated a significant increase in the expression of mature miR-181a-5p and primary miR-181a, with fold changes of 294% and 292%, respectively, in comparison to the untreated controls (Fig. 6 D and E).

#### qPCR for the expression of mature and primary miRNAs after miR-216a-5p*-CEDS

In the multiple qPCRs for 15 mature miRNAs, mmu-miR-216a-5p*-CEDS TAATCTCAGCTGGCAACTGTGA*-CEDS demonstrated a 466% increase in miR-216a-5p expression (Cq=16.62) compared to the untreated control (Cq=14.29) in RAW 264.7 cells. Concurrently, miR-216a-5p*-CEDS was found to significantly increase the numbers of miR-26a-5p and miR-126-5p compared to the untreated control (Fig. 7 A and B).

**Fig. 7.**
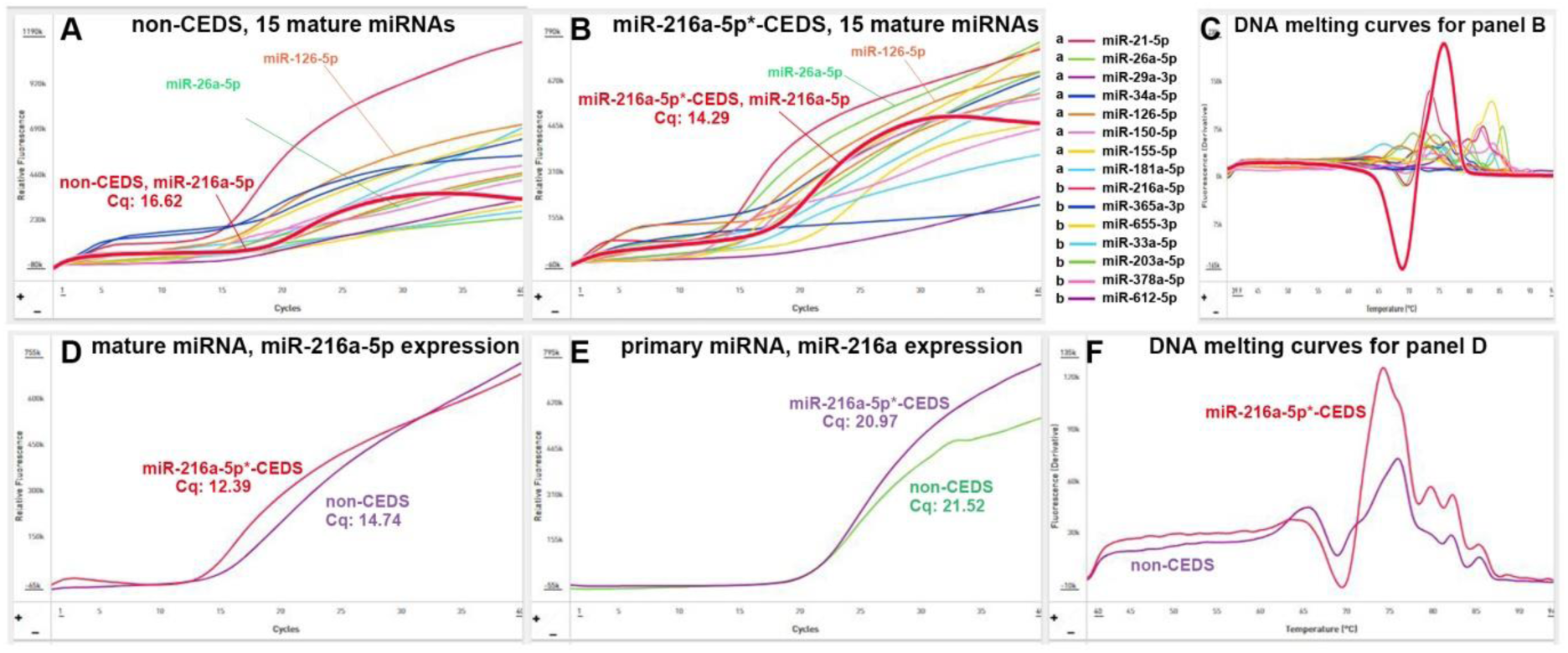
Quantitative PCR for CEDS effect on the expression of mature miR-216a-5p (D) and primary miR-216a (E) in RAW 264.7 cells. MiR-216a-5p*-CEDS increased the expression of miR-216a-5p along with an influence on the expression of other 14 miRNAs (A and B). Noted the variable melting curves of 15 mature miRNAs and similar melting curves of mature miR-216a-5p (C and F). The representative results were presented based on the findings of triple repeated experiments.

In the qPCR assays conducted to determine CEDS-induced incremental ratio of mature and primary miRNAs, miR-216a-5p*-CEDS demonstrated a significant increase in the expression of mature miR-216a-5p and primary miR-216a, with fold changes of 470% and 112%, respectively, in comparison to the untreated controls (Fig. 7 D and E).

#### qPCR for the expression of mature and primary miRNAs after miR-365a-3p*-CEDS

In the multiple qPCRs for 15 mature miRNAs, mmu-miR-365a-3p sequence TAATGCCCCTAAAAATCCTTAT*-CEDS demonstrated a 322% increase in miR-365a-3p expression (Cq=17.12) compared to the untreated control (Cq=15.51) in RAW 264.7 cells. Concurrently, miR-365a-3p*- CEDS was found to significantly increase the expression of miR-216a-5p compared to the untreated control, while decreasing the expression of miR-26a-5p and miR-126-5p (Fig. 8 A and B).

**Fig. 8.**
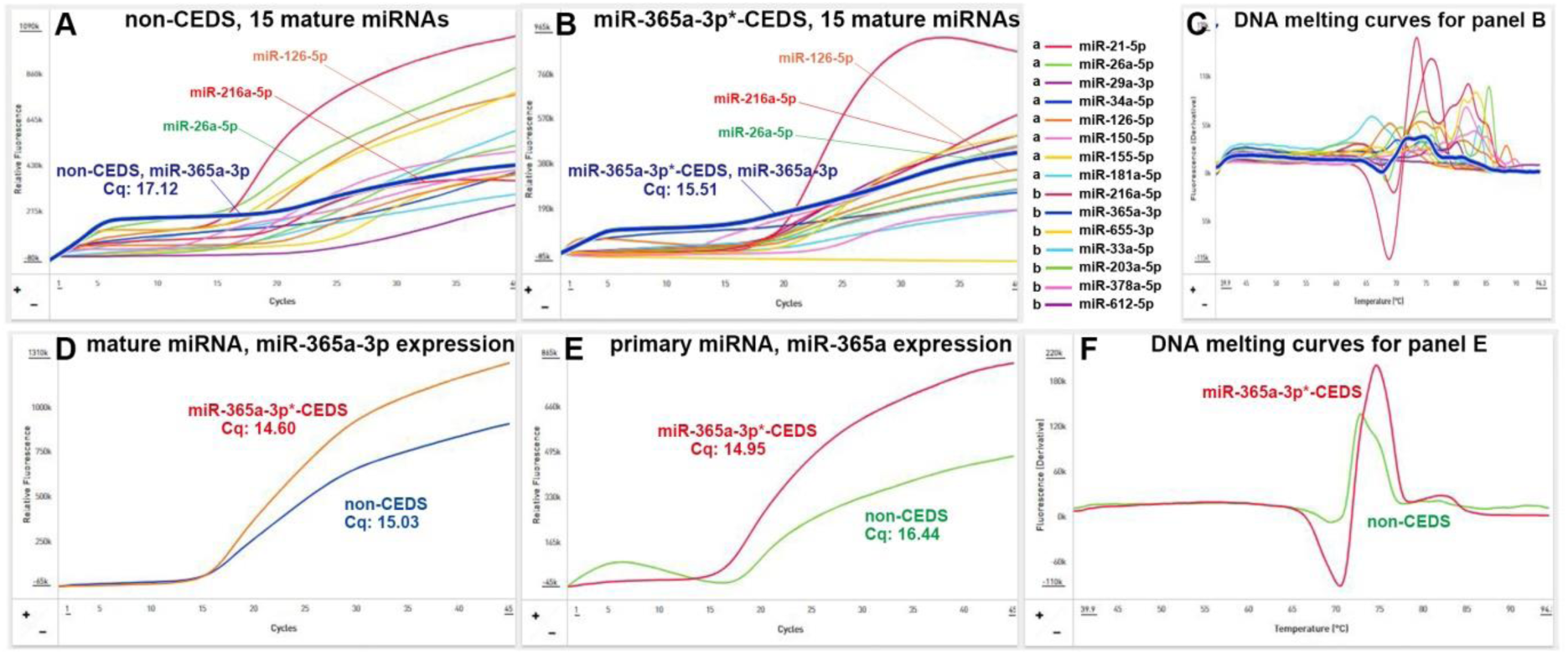
Quantitative PCR for CEDS effect on the expression of mature miR-365a-3p (D) and primary miR-365a (E) in RAW 264.7 cells. MiR-365a-3p*-CEDS increased the expression of miR-365a-3p along with an influence on the expression of other 14 miRNAs (A and B). Noted the variable melting curves of 15 mature miRNAs and similar melting curves of primary miR-365a (C and F). The representative results were presented based on the findings of triple repeated experiments.

In the qPCR assays conducted to determine CEDS-induced incremental ratio of mature and primary miRNAs, miR-365a-3p*-CEDS demonstrated a significant increase in the expression of mature miR-365a-3p and primary miR-365a, with fold changes of 86% and 298%, respectively, in comparison to the untreated controls (Fig. 8 D and E).

#### qPCR for the expression of mature and primary miRNAs after miR-655-3p*-CEDS

In the multiple qPCRs for 15 mature miRNAs, mmu-miR-655-3p sequence ATAATACATGGTTAACCTCTTT*-CEDS demonstrated a 124% increase in miR-655-3p expression (Cq=15.52) compared to the untreated control (Cq=14.9) in RAW 264.7 cells. Concurrently, miR-655-3p*- CEDS was found to significantly increase the expression of miR-365a-3p compared to the untreated control, while decreasing the expression of miR-21-5p (Fig. 9 A and B).

**Fig. 9.**
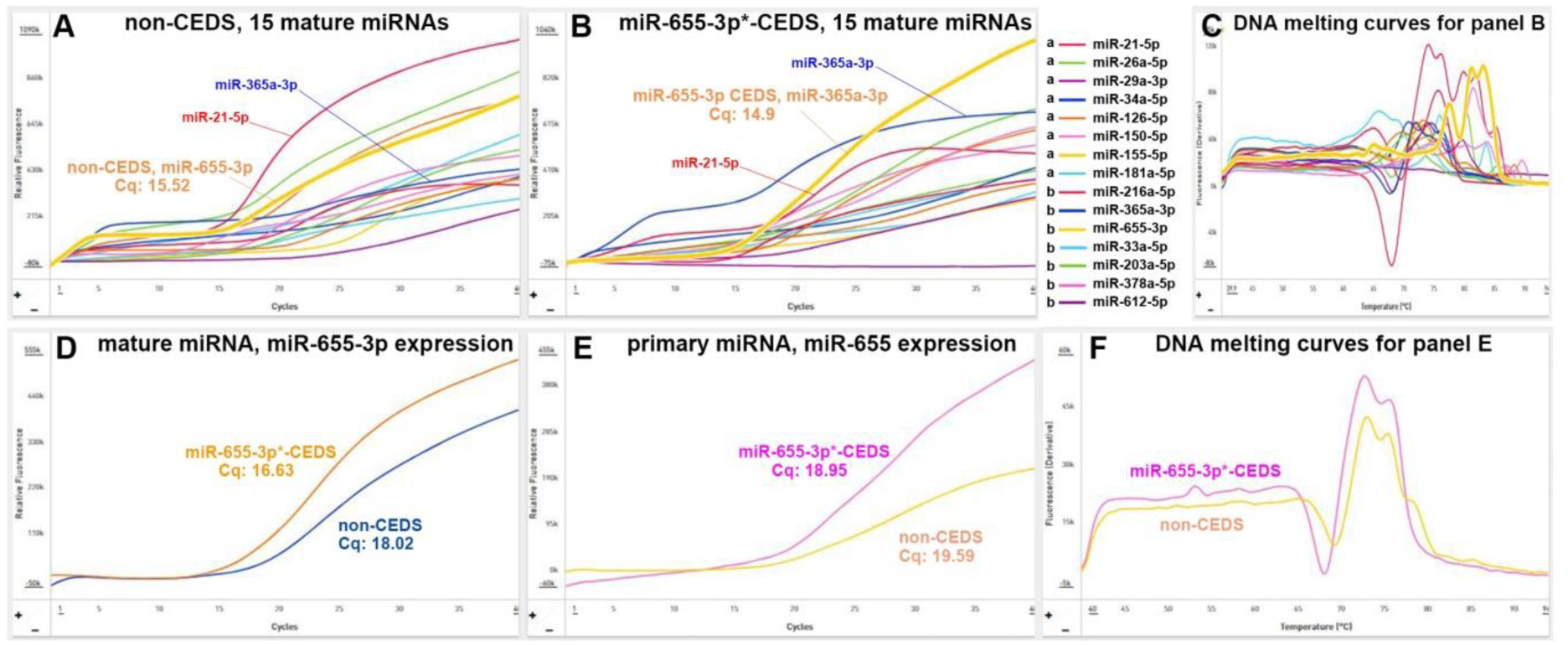
Quantitative PCR for CEDS effect on the expression of mature miR-655-3p (D) and primary miR-655 (E) in RAW 264.7 cells. MiR-655-3p*-CEDS increased the expression of miR-655-3p along with an influence on the expression of other 14 miRNAs (A and B). Noted the variable melting curves of 15 mature miRNAs and similar melting curves of primary miR-655 (C and F). The representative results were presented based on the findings of triple repeated experiments.

In the qPCR assays conducted to determine CEDS-induced incremental ratio of mature and primary miRNAs, miR-655-3p*-CEDS demonstrated a significant increase in the expression of mature miR-655-3p and primary miR-655, with fold changes of 278% and 128%, respectively, in comparison to the untreated controls (Fig. 9 D and E).

The qPCR results indicated that CEDS has the potential to increase the expression of target mature and primary miRNAs compared to untreated controls. Although the incremental ratios calculated based on the Cq number showed considerable variability in the three replicate qPCRs, there was a consistent trend showing that CEDS increased the expression of target mature and primary miRNAs in the qPCR experiments for nine miRNAs observed in this study.

The sequences of nine miRNAs observed in this study demonstrated no homology with each other in the search of the align sequence nucleotide blast program (NCBI, USA). However, it was found that CEDS targeting a mature miRNA can not only affect the expression of its own miRNA, but also the expression of other mature miRNAs. This phenomenon may be relevant to the common fact that one miRNA can regulate multiple genes, and one gene can be regulated by multiple miRNAs^17^.

To illustrate, if A-miRNA and B-miRNA are capable of targeting the same protein-α, an increase of 50% in the expression of A-miRNA may result in the targeting of the entire number of protein-α genes, thereby freeing up the B-miRNA to target other protein genes. Consequently, it is postulated that the increased number of A-miRNA will not impede the functions of B-miRNA, but rather stimulate B-miRNAs to target other protein genes secondarily.

In the present study, the qPCRs were simultaneously performed to assess 15 miRNAs at once, and its results showed a consistent trend, indicating the secondary effect of CEDS, even though the presence of slight variation in the three-times repeated qPCR results. Nevertheless, since the qPCR is the easy and rapid method to identify the secondary effect of CEDS, it would be worthy to note that the secondary effect of CEDS occurs between the relevant miRNAs in their functions as follows.

MiR-26a-5p*-CEDS can increase the expression of miR-126-5p and miR-655-3p while decreasing the expression of miR-365a-3p in RAW 264.7 cells. MiR-34a-5p*-CEDS can increase the expression of miR-126- 5p and miR-655-3p while decreasing the expression of miR-21-5p, and that miR-126-5p*-CEDS can increase the expression of miR-26a-5p and miR-655-3p. MiR-150-5p*-CEDS can increase the expression of miR-126-5p, miR-365a-3p, and miR-655-3p, and that miR-155-5p*-CEDS can increase the expression of miR-181a-5p, miR- 216a-5p, and miR-655-3p. MiR-181a-5p*-CEDS can increase the expression of miR-126-5p, miR-216a-5p, and miR-655-3p, and that miR-216a-5p*-CEDS can increase the numbers of miR-26a-5p and miR-126-5p. MiR- 365a-3p*-CEDS can increase the expression of miR-216a-5p while decreasing the expression of miR-26a-5p and miR-126-5p. And miR-655-3p*-CEDS can increase the expression of miR-365a-3p while decreasing the expression of miR-21-5p.

The secondary effect of CEDS appeared to increase or decrease the expression of other miRNAs. Therefore, it is postulated that the secondary effect of CEDS may be the result of a complex feedback signaling mechanism involving gene transcription and protein translation. However, it can be said that CEDS can not only exert the primary effect on the target miRNAs, but also the secondary effect on the relevant miRNAs in function at the same time.

### IP-HPLC analysis for the changes of the protein signaling after CEDS targeting different miRNAs in RAW 264.7 cells

The RAW 264.7 cells treated with dodecagonal CEDS using a target sequence were collected and lysed using protein lysis buffer (iNtRON Biotechnology INC, Korea), and analyzed by IP-HPLC as follows. Each protein sample was mixed with 5 mL of binding buffer (150mM NaCl, 10mM Tris pH 7.4, 1mM EDTA, 1mM EGTA, 0.2mM sodium vanadate, 0.2mM PMSF and 0.5% NP-40) and incubated with the antibody-bound protein A/G agarose bead column on a rotating stirrer at room temperature for 1 h. Following multiple washes of the columns with Tris-NaCl buffer at a pH of 7.5 and a graded NaCl concentration (0.15–0.3 M), the target proteins were eluted with 300 μL of IgG elution buffer (Pierce, USA). Column elution was conducted using a 0.15 M NaCl/20% acetonitrile solution at a flow rate of 0.5 mL/min.

The immunoprecipitated proteins were subjected to the analysis using a precision HPLC unit equipped with a reverse-phase column and a micro-analytical ultraviolet (UV) detector system (1100 series, Agilent, USA). The control and experimental samples were analyzed in sequence to permit a comparison of their HPLC peaks. The square roots of whole protein peak areas were normalized and compared to the controls to calculate the relative ratio (%) of protein expression level. The ratios of protein levels between the experimental and control groups were divided into five categories, as follows: severe underexpression (below 80%), marked underexpression (80%-below 95%), minimal expression change (95%-below 105%), marked overexpression (105%-120%), and severe overexpression (above 120%).

**Table 2.**
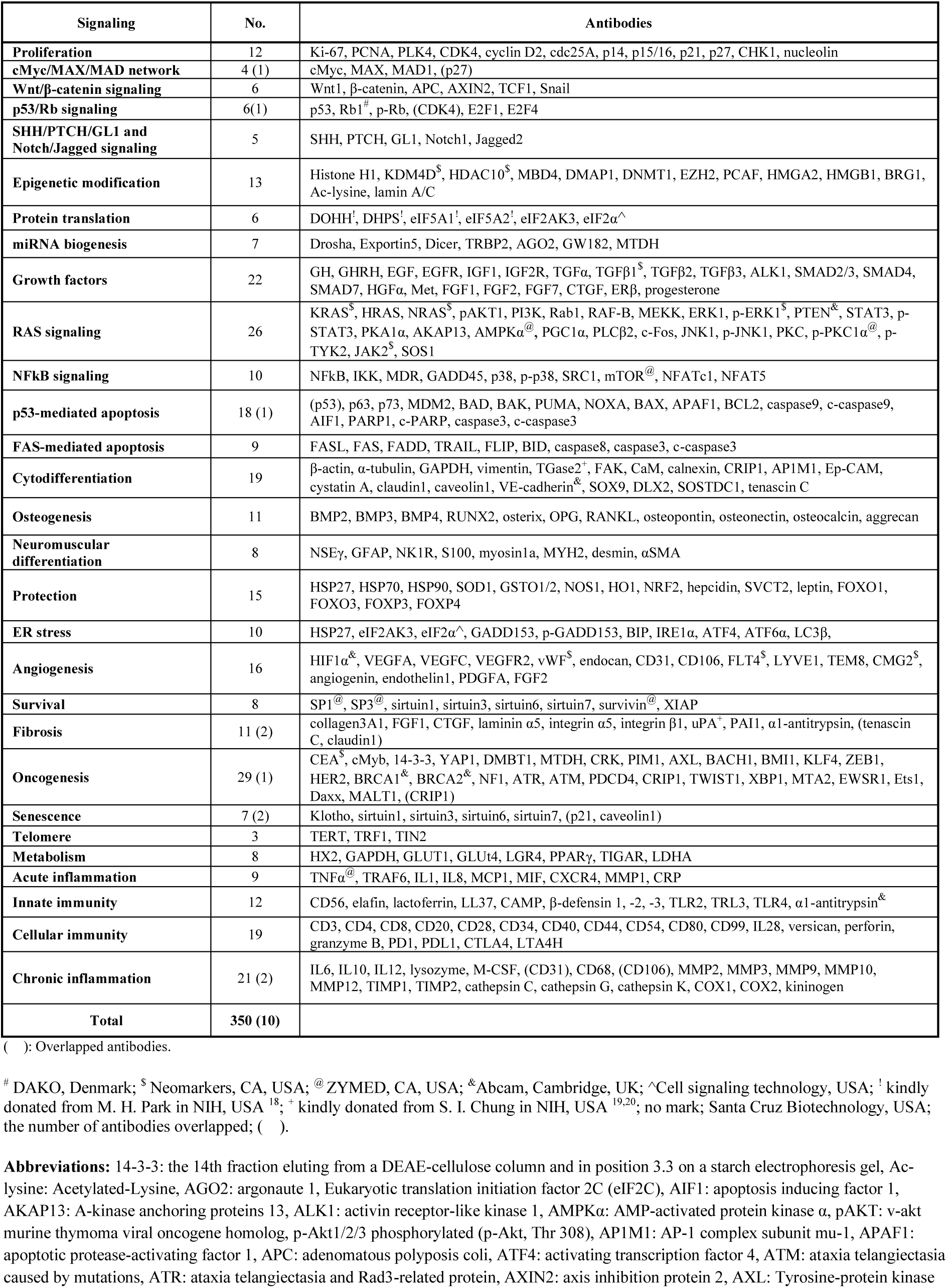
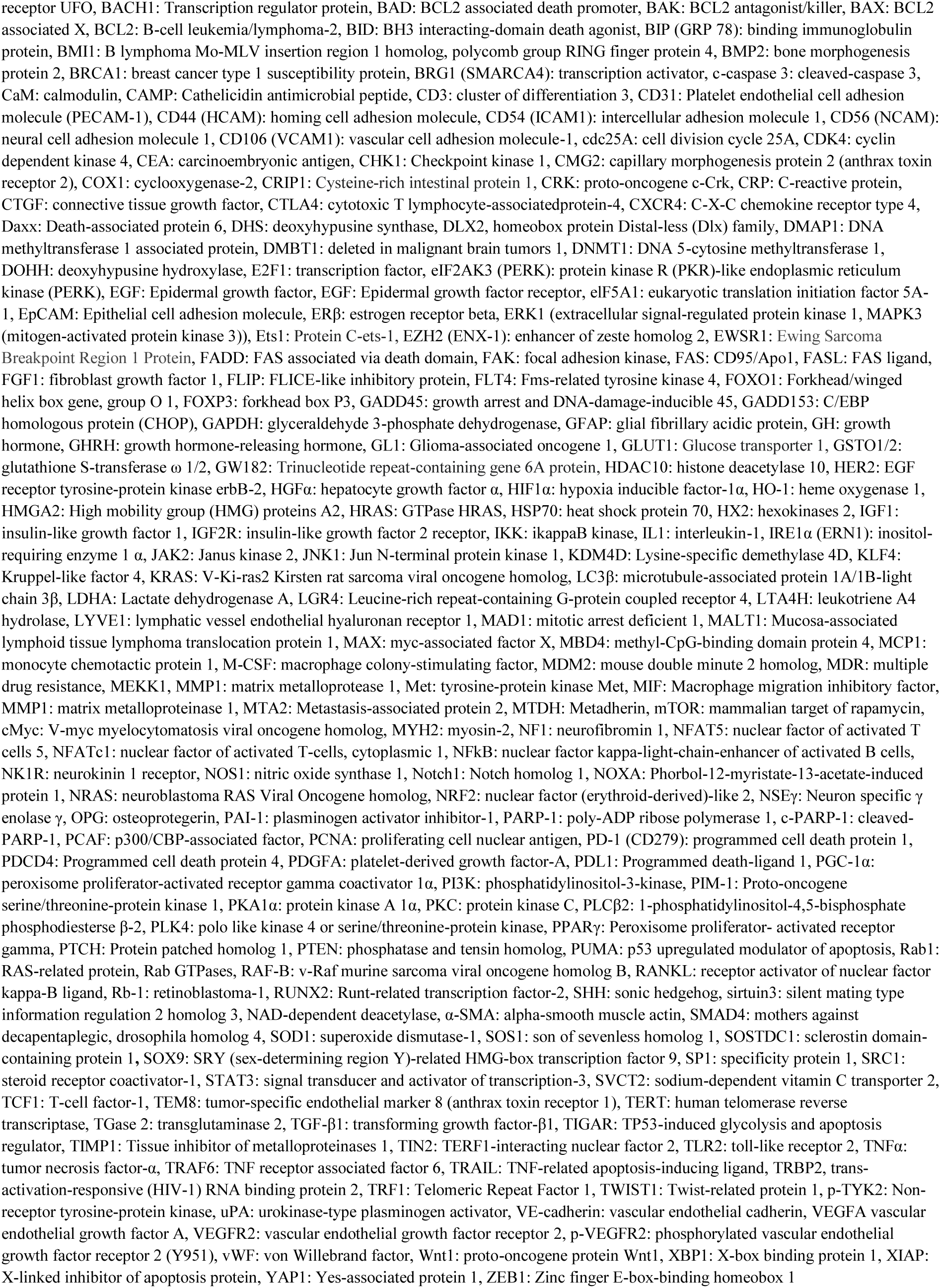
Antibodies used in the IP-HPLC analysis.

| Signaling | No. | Antibodies |
| --- | --- | --- |
| **Proliferation** | 12 | Ki-67, PCNA, PLK4, CDK4, cyclin D2, cdc25A, p14, p15/16, p21, p27, CHK1, nucleolin |
| **cMyc/MAX/MAD network** | 4 (1) | cMyc, MAX, MAD1, (p27) |
| **Wnt/β-catenin signaling** | 6 | Wnt1, β-catenin, APC, AXIN2, TCF1, Snail |
| **p53/Rb signaling** | 6(1) | p53, Rb1<sup>#</sup>, p-Rb, (CDK4), E2F1, E2F4 |
| **SHH/PTCH/GLI and Notch/Jagged signaling** | 5 | SHH, PTCH, GLI, Notch1, Jagged2 |
| **Epigenetic modification** | 13 | Histone H1, KDM4D<sup>§</sup>, HDAC10<sup>§</sup>, MBD4, DMAP1, DNMT1, EZH2, PCAF, HMG2, HMGB1, BRG1, Ac-lysine, lamin A/C |
| **Protein translation** | 6 | DOHH<sup>†</sup>, DHPS<sup>†</sup>, eIF5A1<sup>†</sup>, eIF5A2<sup>†</sup>, eIF2AK3, eIF2$\alpha^{\wedge}$ |
| **miRNA biogenesis** | 7 | Drosha, Exportin5, Dicer, TRBP2, AGO2, GW182, MTDH |
| **Growth factors** | 22 | GH, GHRH, EGF, EGFR, IGF1, IGF2R, TGFα, TGFβ1<sup>§</sup>, TGFβ2, TGFβ3, ALK1, SMAD2/3, SMAD4, SMAD7, HGFα, Met, FGF1, FGF2, FGF7, CTGF, ERβ, progesterone |
| **RAS signaling** | 26 | KRAS<sup>§</sup>, HRAS, NRAS<sup>§</sup>, pAKT1, PI3K, Rab1, RAF-B, MEKK, ERK1, p-ERK1<sup>§</sup>, PTEN<sup>§</sup>, STAT3, p-STAT3, PKA1α, AKAP13, AMPK$\alpha^{\&}$, PGC1α, PLCβ2, c-Fos, JNK1, p-JNK1, PKC, p-PKC1$\alpha^{\&}$, p-TYK2, JAK2<sup>§</sup>, SOS1 |
| **NFkB signaling** | 10 | NFkB, IKK, MDR, GADD45, p38, p-p38, SRC1, mTOR<sup>@</sup>, NFATc1, NFAT5 |
| **p53-mediated apoptosis** | 18 (1) | (p53), p63, p73, MDM2, BAD, BAK, PUMA, NOXA, BAX, APAF1, BCL2, caspase9, c-caspase9, AIF1, PARP1, c-PARP, caspase3, c-caspase3 |
| **FAS-mediated apoptosis** | 9 | FASL, FAS, FADD, TRAIL, FLIP, BID, caspase8, caspase3, c-caspase3 |
| **Cytodifferentiation** | 19 | β-actin, α-tubulin, GAPDH, vimentin, TGase2<sup>†</sup>, FAK, CaM, calnexin, CRIP1, AP1M1, Ep-CAM, cystatin A, claudin1, caveolin1, VE-cadherin<sup>&</sup>, SOX9, DLX2, SOSTDC1, tenascin C |
| **Osteogenesis** | 11 | BMP2, BMP3, BMP4, RUNX2, osterix, OPG, RANKL, osteopontin, osteonectin, osteocalcin, aggrecan |
| **Neuromuscular differentiation** | 8 | NSEγ, GFAP, NK1R, S100, myosin1a, MYH2, desmin, αSMA |
| **Protection** | 15 | HSP27, HSP70, HSP90, SOD1, GSTO1/2, NOS1, HO1, NRF2, hepcidin, SVCT2, leptin, FOXO1, FOXO3, FOSX3, FOSX4 |
| **ER stress** | 10 | HSP27, eIF2AK3, eIF2$\alpha^{\wedge}$, GADD153, p-GADD153, BIP, IRE1α, ATF4, ATF6α, LC3β, |
| **Angiogenesis** | 16 | HIF1$\alpha^{\&}$, VEGFA, VEGFC, VEGFR2, vWF<sup>§</sup>, endocan, CD31, CD106, FLT4<sup>§</sup>, LYVE1, TEM8, CMG2<sup>§</sup>, angiogenin, endothelin1, PDGFA, FGF2 |
| **Survival** | 8 | SP1<sup>@</sup>, SP3<sup>@</sup>, sirtuin1, sirtuin3, sirtuin6, sirtuin7, survivin<sup>@</sup>, XIAP |
| **Fibrosis** | 11 (2) | collagen3A1, FGF1, CTGF, laminin α5, integrin α5, integrin β1, uPA<sup>†</sup>, PAI1, α1-antitrypsin, (tenascin C, claudin1) |
| **Oncogenesis** | 29 (1) | CEA<sup>§</sup>, cMyb, 14-3-3, YAP1, DMBT1, MTDH, CRK, PIM1, AXL, BACH1, BMI1, KLF4, ZEB1, HER2, BRCA1<sup>&</sup>, BRCA2<sup>&</sup>, NF1, ATR, ATM, PDCD4, CRIP1, TWIST1, XBP1, MTA2, EWSR1, Ets1, Daxx, MALT1, (CRIP1) |
| **Senescence** | 7 (2) | Klotho, sirtuin1, sirtuin3, sirtuin6, sirtuin7, (p21, caveolin1) |
| **Telomere** | 3 | TERT, TRF1, TIN2 |
| **Metabolism** | 8 | HX2, GAPDH, GLUT1, GLUT4, LGR4, PPARγ, TIGAR, LDHA |
| **Acute inflammation** | 9 | TNF$\alpha^{\&}$, TRAF6, IL1, IL8, MCP1, MIF, CXCR4, MMP1, CRP |
| **Innate immunity** | 12 | CD56, elafin, lactoferrin, LL37, CAMP, β-defensin 1, -2, -3, TLR2, TLR3, TLR4, α1-antitrypsin<sup>&</sup> |
| **Cellular immunity** | 19 | CD3, CD4, CD8, CD20, CD28, CD34, CD40, CD44, CD54, CD80, CD99, IL28, versican, perforin, granzyme B, PD1, PDL1, CTLA4, LTA4H |
| **Chronic inflammation** | 21 (2) | IL6, IL10, IL12, lysozyme, M-CSF, (CD31), CD68, (CD106), MMP2, MMP3, MMP9, MMP10, MMP12, TIMP1, TIMP2, cathepsin C, cathepsin G, cathepsin K, COX1, COX2, kininogen |
| **Total** | **350 (10)** |  |

( ): Overlapped antibodies.
<sup>#</sup> DAKO, Denmark; <sup>§</sup> Neomarkers, CA, USA; <sup>@</sup> ZYMED, CA, USA; <sup>&</sup> Abcam, Cambridge, UK; <sup>^</sup> Cell signaling technology, USA; <sup>†</sup> kindly donated from M. H. Park in NIH, USA <sup>18</sup>; <sup>+</sup> kindly donated from S. I. Chung in NIH, USA <sup>19,20</sup>; no mark; Santa Cruz Biotechnology, USA; the number of antibodies overlapped; ( ).
**Abbreviations:** 14-3-3: the 14th fraction eluting from a DEAE-cellulose column and in position 3.3 on a starch electrophoresis gel, Ac-lysine: Acetylated-Lysine, AGO2: argonaute 1, Eukaryotic translation initiation factor 2C (eIF2C), AIF1: apoptosis inducing factor 1, AKAP13: A-kinase anchoring proteins 13, ALK1: activin receptor-like kinase 1, AMPKα : AMP-activated protein kinase $\alpha$ , pAKT: v-akt murine thymoma viral oncogene homolog, p-Akt1/2/3 phosphorylated (p-Akt, Thr 308), AP1M1: AP-1 complex subunit mu-1, APAF1: apoptotic protease-activating factor 1, APC: adenomatous polyposis coli, ATF4: activating transcription factor 4, ATM: ataxia telangiectasia caused by mutations, ATR: ataxia telangiectasia and Rad3-related protein, AXIN2: axis inhibition protein 2, AXL: Tyrosine-protein kinase

#### MiR-34a-5p*-CEDS influenced the protein signaling in RAW 264.7 cells

RAW 264.7 cells treated with mmu-miR-34a-5p sequence TGGCAGTGTCTTAGCTGGTTGT*-CEDS showed the characteristic protein expressions, resulting in the suppression of proliferation, cMyc/MAX/MAD network, Wnt/β-catenin signaling, protein translation, miRNA biogenesis, growth factors, cytodifferentiation, neuromuscular differentiation, RAS signaling, NFkB signaling, protection, ER stress, angiogenesis, survival, acute inflammation, cellular immunity, chronic inflammation, FAS-mediated apoptosis, PARP-mediated apoptosis, fibrosis, glycolysis, and telomere signaling, as well as the activation of p53/Rb/E2F1 signaling, SHH/PTCH/GLI signaling, Notch/Jagged signaling, epigenetic modification, osteogenesis, innate immunity, chronic inflammation, p53-mediated apoptosis, oncogenesis, and senescence signaling compared to the untreated controls (Fig. 11A).

**Fig. 11.**
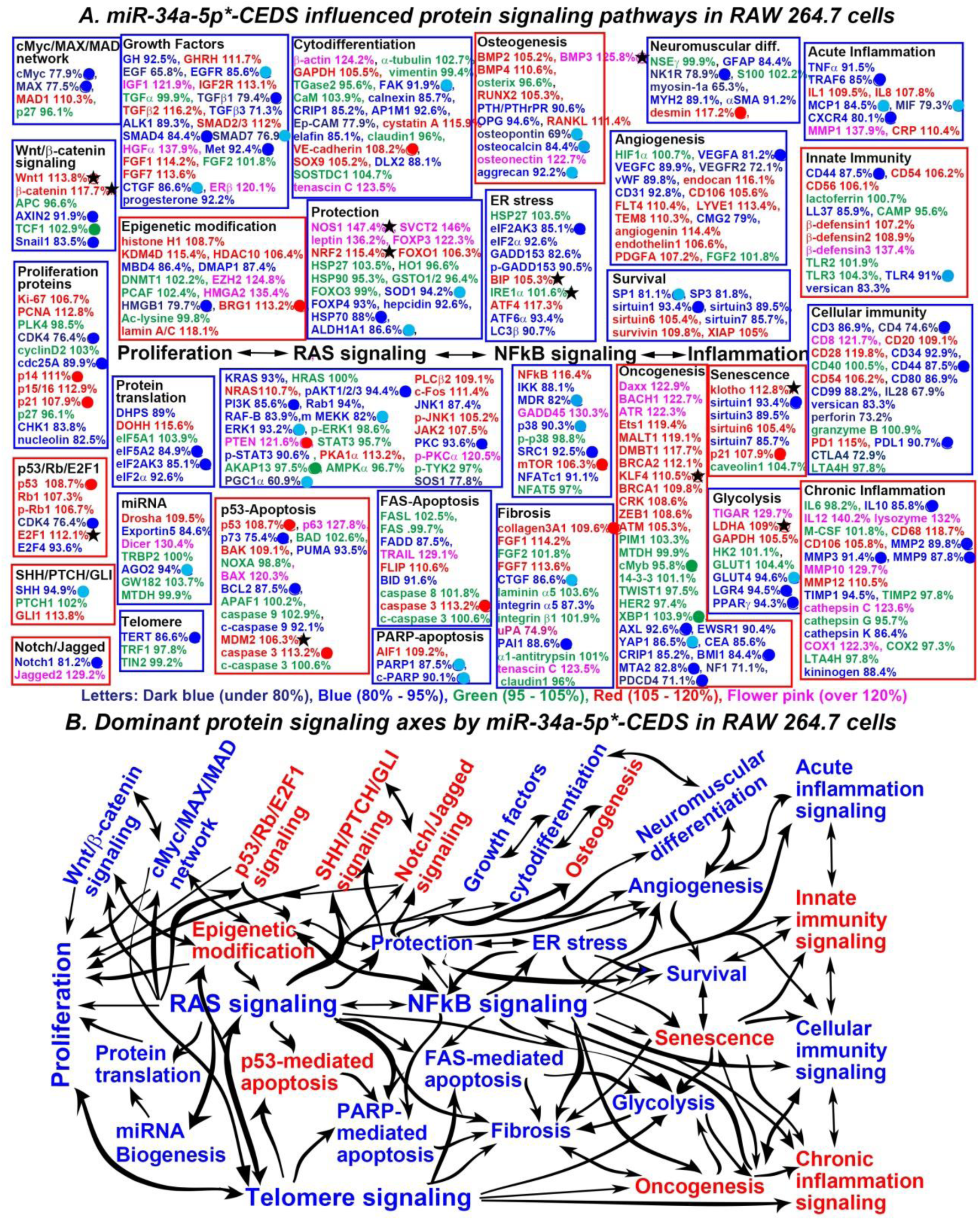
MiR-34a-5p*-CEDS influenced the protein signaling (A) and axes (B) in RAW 264.7 cells. The IP- HPLC revealed proteins downregulated 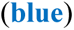, upregulated 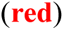, and minimally changed 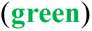 compared to the untreated controls. Dominantly suppressed 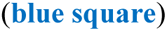 and activated 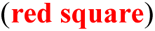 signaling. Downregulated proteins by direct targeting 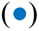 and indirect response 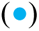, upregulated proteins by indirect response 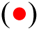, and minimal changed (±5%) proteins 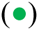. Some target proteins 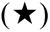 were unexpectedly upregulated contrast to the current manuscripts and data from miRDB and TargetScan websites.

**The proliferation signaling** was suppressed by downregulating miR-34a-5p target proteins CDK4 (76.4%) and cdc25A (89.9%), and p27 (92. 3%), as well as coincidentally downregulating CHK1 (83.8%) and nucleolin (82.5%), despite the upregulation of miR-34a-5p-responsible proteins p14 (111%) and p21 (107.9%) and coincidental upregulation of Ki-67 (106.7%), PCNA (112.8%), and p15/16 (112.9%).

**The cMyc/MAX/MAD network** was suppressed by downregulating miR-34a-5p target proteins cMyc (77.9%) and MAX (77.5%), and coincidental upregulation of MAD1 (110.3%).

**The Wnt/β-catenin signaling** was suppressed by downregulating miR-34a-5p target proteins AXIN2 (91.9%) and Snail1 (83.5%), despite the upregulation of Wnt1 (113.8%) and β-catenin (117.7%). Wnt1 and β- catenin are miR-34a-5p target proteins, but they were unexpectedly upregulated compared to the reference data.

**The protein translation signaling** was suppressed by downregulating miR-34a-5p target proteins eIF5A2 (84.9%) and eIF2AK3 (85.1%), and coincidentally downregulating DHPS (89%) and eIF2α (92.6%), despite the upregulation of DOHH (115.6%).

**The miRNA biogenesis signaling** appears to be attenuated by downregulating miR-34a-5p-responsible protein AGO2 (94%) and coincidentally downregulating Exportin5 (84.6%), despite the upregulation of Drosha (109.5%) and Dicer (130.4%).

**The growth factor signaling** appears to be activated by downregulating miR-34a-5p target proteins TGFβ1 (79.4%), SMAD4 (84.4%), and Met (92.4%) and miR-34a-5p-responsible proteins EGFR (85.6%), SMAD7 (76.9%), and CTGF (86.6%), as well as coincidental downregulation of GH (92.5%), EGF (65.8%), TGFβ3 (71.3%), ALK1 (89.3%), and progesterone (92.2%), despite the upregulation of GHRH (111.7%), IGF1 (121.9%), IGF2R (113.1%), TGFβ2 (116.2%), SMAD2/3 (112%), HGFα 137.9%), FGF1 (114.2%), FGF7 (113.6%), and ERβ (120.1%).

**The cytodifferentiation signaling** appear to be suppressed by downregulating miR-34a-5p-responsible protein FAK (91.9%) and coincidentally downregulating calnexin (85.7%), CRIP1 (85.2%), AP1M1 (92.6%), Ep-CAM (77.9%), elafin (85.1%), and DLX2 (88.1%), despite the upregulation of miR-34a-5p-responsible protein VE-cadherin (108.2%) and coincidental upregulation of β-actin (124.2%), GAPDH (105.5%), cystatin A (115.9%), SOX9 (105.2%), and tenascin C (123.5%).

**The neuromuscular differentiation signaling were** suppressed by downregulating miR-34a-5p target protein NK1R (78.9%) and coincidentally downregulating GFAP (84.4%), myosin-1a (65.3%), MYH2 (89.1%), and αSMA (91.2%), despite the upregulation of miR-34a-5p-responsible protein desmin (117.2%).

**The RAS signaling** was suppressed by downregulating miR-34a-5p target proteins pAKT1/2/3 (94.4%), PI3K (85.6%), and PKC (93.6%) and downregulating miR-34a-5p-responsible proteins MEKK (82%), ERK1 (93.2%), PGC1α (60.9%), as well as coincidental downregulation of KRAS (93%), Rab1 (94%), RAF-B (83.9%), p-STAT3 (90.6%), JNK1 (87.4%), and SOS1 (77.8%), despite the upregulation of miR-34a-5p- responsible protein PTEN (121.8%) and coincidental upregulation of NRAS (110.7%), PKA1α (113.2%), PLCβ2 (109.1%), c-Fos (111.4%), p-JNK1 (105.2%), JAK2 (107.5%), and p-PKCα (120.5%).

**The NFkB signaling** was suppressed by downregulating miR-34a-5p target proteins SRC1 (92.5%) and downregulating miR-34a-5p-responsible proteins MDR (82%) and p38 (90.3%), as well as coincidental downregulation of IKK (88.1%) and NFATc1 (91.1%), despite the upregulation of miR-34a-5p-responsible protein mTOR (106.3%) and coincidental upregulation of NFkB (116.4%) and GADD45 (130.3%).

**The protection signaling** was suppressed by downregulating miR-34a-5p target protein HSP70 (88%) and downregulating miR-34a-5p-responsible proteins SOD1 (94.2%) and ALDH1A1 (86.6%), as well as GADD153 (105.6%), ATF4 (106%), and ATF6α (111.4%), as well as coincidental downregulation of FOXP4 (93%) and hepcidin (92.6%), despite the upregulation of NOS1 (147.4%), SVCT2 (146%), leptin (136.2%), FOXP (122.3%), NRF2 (115.4%), and FOXO1 (106.3%). NOS1 and NRF2 are miR-34a-5p target proteins, but they were unexpectedly upregulated compared to the reference data.

**The ER stress signaling** was suppressed by downregulating miR-34a-5p target protein eIF2AK3 (85.1%) and coincidentally downregulating eIF2α (92.6%), GADD153 (82.6%), p-GADD153 (90.5%), ATF6α (93.4%), and LC3β (90.7%), despite the upregulation of BIP (105.3%) and ATF4 (117.3%). BIP is a miR-34a-5p target protein, but it was unexpectedly upregulated compared to the reference data.

**The angiogenesis signaling** was suppressed by downregulating miR-34a-5p target protein VEGFA (81.2%) and coincidentally downregulating VEGFC (89.9%), VEGFR2 (72.1%), vWF (89.8%), CD31 (92.8%), and CMG2 (79%), despite the upregulation of endocan (116.1%), CD106 (105.6%), FLT4 (110.4%), LYVE1 (113.4%), TEM8 (110.3%), angiogenin (114.4%), endothelin1 (106.6%), and PDGFA (107.2%).

**The survival signaling** was suppressed by downregulating miR-34a-5p target protein sirtuin1 (93.4%) and downregulating the miR34a-5p-responsible protein SP1 (81.1%), as well as coincidental downregulation of SP3 (81.8%), sirtuin3 (89.5%), and sirtuin7 (85.7%), despite the upregulation of sirtuin6 (105.4%), survivin (109.8%), and XIAP (105%).

**The acute inflammation signaling** was suppressed by downregulating miR-34a-5p target proteins TRAF6 (85%) and CXCR4 (80.1%), and downregulating miR-34a-5p-responsible proteins MCP1 (84.5%) and MIF (79.3%), and coincidental downregulation of TNFα (91.5%), despite the slight upregulation of IL1 (109.5%), IL8 (107.8%), CRP (110.4%), and MMP1 (137.9%).

**The cellular immunity signaling** appears to be suppressed by downregulating miR-34a-5p target proteins CD4 (74.6%), CD44 (87.5%) and coincidentally downregulating CD3 (86.9%), CD34 (92.9%), CD80 (86.9%), CD99 (88.2%), IL28 (67.9%), versican (83.3%), perforin (73.2%), and CTLA4 (72.9%) and upregulating PD1 (115%), despite the upregulation of CD8 (121.7%), CD20 (109.1%), CD28 (119.8%), and CD54 (106.2%) and downregulation of miR-34a-5p target protein PDL1 (90.7%).

**The FAS-mediated apoptosis signaling** was suppressed by downregulating FADD (87.5%) and BID (91.6%), and coincidentally upregulating FLIP (110.6%), despite the upregulation of TRAIL (129.1%) and caspase3 (113.2%). Caspase3 was alternatively upregulated as a miR-34a-5p-responsible protein.

**The PARP-mediated apoptosis signaling** was suppressed by downregulating miR-34a-5p-responsible proteins PARP1 (87.5%) and c-PARP (90.1%), despite the upregulation of AIF1 (109.2%).

**The fibrosis signaling** was suppressed by downregulating miR-34a-5p target protein PAI1 (88.6%) and miR-34a-5p-responsible protein CTGF (86.6%), as well as coincidentally downregulating integrin α1 (87.3%) and upregulating FGF7 (113.6%), despite the upregulation of collagen3A1 (109.6%), FGF1 (114.2%), uPA (74.9%), and tenascin C (123.5%). Collagen 3A1 is upregulated as a miR-34a-5p-responsible protein.

**The glycolysis signaling** was suppressed by downregulating miR-34a-5p-responsible proteins LGR4 (94.5%), PPARγ (94.3%), and miR-34a-5p-responsible protein GLUT4 (94.6%), despite the upregulation of TIGAR (129.7%), LDHA (109%), and GAPDH (105.5%). LDHA is a miR-34a-5p target protein, but it was unexpectedly upregulated compared to the reference data.

**The telomere signaling** was suppressed by downregulating miR-34a-5p target protein TERT (87%).

Conversely, **the p53/Rb/E2F1 signaling** was activated by upregulating miR-34a-5p target protein E2F1 (112.1%) and upregulating miR-34a-5p-responsible protein p53 (108.7%), as well as coincidental upregulation of Rb1 (107.3%) and E2F1 (112.1%), and downregulating E2F4 (93.6%), despite the downregulation of CDK4 (76.4%). E2F1 is a miR-34a-5p target protein, but it was unexpectedly upregulated compared to the reference data.

**The SHH/PTCH/GLI signaling** was activated by upregulating GLI1 (113.8%), despite the downregulation of miR-34a-5p-responsible protein SHH (94.9%).

**The Notch/Jagged signaling** was activated by upregulating Jagged2 (129.2%), despite the downregulation of miR-34a-5p target protein Notch1 (81.2%).

**The osteogenesis signaling** appears to be activated by upregulating BMP2 (105.2%), BMP3 (125.8%), BMP4 (110.6%), RUNX2 (105.3%), RANKL (111.4%), and osteonectin (122.7%), despite the downregulation of miR-34a-5p-responsible proteins osteopontin (69%), osteocalcin (84.4%), aggrecan (92.2%), and coincidental downregulation of PTH/PTHrPR (90.6%), and OPG (94.6%).

**The innate immunity signaling** appear to be activated by upregulating CD54 (106.2%), CD56 (106.1%), β-defensin1 (107.2%), β-defensin2 (108.9%), and β-defensin3 (137.4%), despite the downregulation of miR- 34a-5p target protein CD44 (87.5%) and miR-34a-5p-responsible protein TLR4 (91%), as well as the coincidental downregulation of LL37 (85.9%) and versican (83.3%).

**The chronic inflammation signaling** appears to be activated by upregulating IL12 (140.2%), lysozyme (132%), CD68 (118.7%), CD106 (105.8%), MMP10 (129.7%), MMP12 (110.5%), cathepsin C (123.6%), and COX1 (122.3%) and coincidentally downregulating miR-34a-5p target protein IL10 (85.8%), despite the downregulation of miR-34a-5p target proteins MMP2 (89.8%), MMP3 (91.4%), and MMP9 (87.8%) and coincidental downregulation of TIMP1 (94.5%), cathepsin K (86.4%), and kininogen (88.4%).

**The p53-mediated apoptosis signaling** was activated by downregulating miR-34a-5p target protein BCL2 (87.5%) and upregulating miR-34a-5p-responsible proteins p53 (108.7%) and caspase3 (113.2%), as well as coincidentally upregulating p63 (127.8%), BAK (109.1%), BAX (120.3%), and MDM2 (106.3%), despite the downregulation of miR-34a-5p target protein p73 (75.4%) and coincidental downregulation of PUMA (93.5%) and c-caspase9 (92.1%).

**The senescence signaling** was activated by upregulating miR-34a-5p-responsible protein p21 (107.9%) and coincidentally upregulating klotho (112.8%) and sirtuin6 (105.4%), despite the downregulation of miR-34a- 5p target protein sirtuin1 (93.4%) and coincidental downregulation of sirtuin3 (89.5%) and sirtuin7 (85.7%). Klotho is a miR-34a-5p target protein, but it was unexpectedly upregulated compared to the reference data.

In addition, miR-34a-5p*-CEDS variably influenced **the epigenetic modification signaling** by downregulating miR-34a-5p target proteins HMGB1 (79.7%) and coincidentally downregulating MBD4 (86.4%) and DMAP1 (87.4%), while it upregulated histone H1 (108.7%), KDM4D (115.4%), HDAC10 (106.4%), EZH2 (124.8%), BRG1 (113.2%), and lamin A/C (118.1%). BRG1 is a miR-34a-5p-responsible protein to be positively regulated. Consequently, miR-34a-5p*-CEDS induced a trend towards an increase in the methylation of histones and DNAs, and promotion of transcriptional repression.

MiR-34a-5p*-CEDS intensively suppressed **the oncogenesis signaling** by downregulating miR-34a-5p target proteins AXL (92.6%), BMI1 (84.4%), MTA2 (82.8%), and PDCD4 (71.1%) and miR-34a-5p- responsible protein YAP1 (86.5%), while it widely enhanced **the oncogenesis signaling** by upregulating Daxx (122.9%), BACH1 (122.7%), ATR (122.3%), Ets1 (119.4%), MALT1 (119.1%), DMBT1 (117.7%), BRCA2 (112.1%), KLF4 (110.5%), BRCA1 (109.8%), CRK (108.6%), ZEB1 (108.6%), and ATM (105.3%). KLF4 is a miR-34a-5p target protein, but it was unexpectedly upregulated compared to the reference data. Among the 28 oncoproteins, 12 were found to be overexpressed, nine were underexpressed, and seven exhibited minimal change in expression compared to the untreated controls.

Consequently, miR-34a-5p*-CEDS markedly affect the whole protein signaling in RAW 264.7 cells. MiR- 34a-5p*-CEDS resulted in the attenuation of the RAS-NFkB signaling axis, which subsequently led to the inactivation of proliferation-related signaling axis, protection-ER stress-survival signaling axis, telomere- glycolysis-fibrosis signaling axis, and FAS- and PARP-mediated apoptosis signaling axis. Conversely, miR- 34a-5p*-CEDS activated the p53/Rb/E2F1-SHH/PTCH/GLI-Notch/Jagged proliferation signaling axis, epigenetic modification-RAS-p53 associated apoptosis signaling axis, and oncogenesis-senescence-innate immunity-chronic inflammation signaling axis (Fig. 11B, Table 4).

**Table 4.** Protein expressions in the RAW 264.7 cells treated with miR-34a-5p*-CEDS.

| Signaling | Downregulated Proteins | Upregulated Proteins |
| --- | --- | --- |
| Proliferation | CDK4, cdc25A, CHK1, nucleolin | Ki-67, PCNA, p14, p15/16, p21 |
| cMyc/MAX/MAD network | cMyc, MAX | MAD1 |
| Wnt/β-catenin signaling | AXIN2, Snail | Wnt1★, β-catenin★ |
| p53/Rb/E2F1 signaling | CDK4 | p53, RB1, p-Rb1, E2F1★, E2F4 |
| SHH/PTCH/GLI signaling | SHH | GLI1 |
| Notch/Jagged signaling | Notch | Jagged2 |
| Epigenetic modification | MBD4, DMAP, HMGB1 | Histone H1, KDM4D, HDAC10, EZH2, HMGA2, BRG1, lamin A/C |
| Protein translation | DHPS, eIF5A2, eIF2AK3, eIF2α | DOHH |
| miRNA biogenesis | Exportin5, AGO2 | Drosha, Dicer |
| Growth factors | GH, EGF, EGFR, TGFβ1, TGFβ3, ALK1, SMAD4, SMAD7, Met, CTGF, progesterone | GHRH, IGF1, IGF2, TGFβ2, SMAD2/3, HGFα, FGF1, FGF7, ERβ |
| Cytodifferentiation | FAK, Clanexin, CRIP1, AP1M1, Ep-CAM, elafin, DLX2 | β-actin, GAPDH, cystatin A, VE-cadherin, SOX9, tenascin C |
| Osteogenesis | PTH/PTHrPR, OPG, osteopontin, osteocalcin, aggrecan | BMP2, BMP3★, BMP4, RUNX2, RANKL, osteonectin |
| Neuromuscular differentiation | GFAP, NK1/R, myosin-1a, MYH2, αSMA | Desmin |
| RAS signaling | KRAS, pAKT1/2/3, PI3K, Rab1, RAF-B, MEKK, ERK1, p-STAT3, PGC1α, JNK1, PKC | NRAS, PTEN, PKA1α, PLCβ2, c-Fos, p-JNK1, JAK2, p-PKCα |
| NFκB signaling | IKK, MDR, p38, SRC1, NFATc1 | NFκB, GADD45, mTOR |
| p53-mediated apoptosis | p73, PUMA, BCL2, c-caspase9 | p53, p63, BAK, BAX, MDM2★, caspase3 |
| FAS-mediated apoptosis | FADD, BID | TRAIL, FLIP, caspase3 |
| PARP-mediated apoptosis | PARP1, c-PARP | AIF1 |
| Protection | SOD1, FOXp4, hepcidin, HSP70, ALDH1A1 | NOS1★, SVCT2, leptin, FOXp3, NRF2★, FOXO1 |
| ER stress | eIF2AK3, eIF2α, GADD153, p-GADD153, ATF6α, LC3β | BIP★, ATF4 |
| Angiogenesis | VEGFA, VEGFC, VEGFR2, vWF, CD31, CMG2 | endocan, CD106, FLT4, LYVE1, TEM8, angiogenin, endothelin1, PDGFA |
| Survival | SP1, SP3, sirtuin1, sirtuin3, sirtuin7 | sirtuin6, survivin, XIAP |
| Acute inflammation | TNFα, TRAF6, MCP1, MIF, CXCR4 | IL1, IL8, MMP1, CRP |
| Innate immunity | CD44, LL37, TLR4, versican | CD54, CD56, β-defensin1, β-defensin2, β-defensin3, |
| Cellular immunity | CD3, CD4, CD34, CD44, CD80, CD99, IL28, perforin, PDL1, CTLA4 | CD8, CD20, CD28, PD1 |
| Chronic inflammation | IL10, MMP2, MMP3, MMP9, TIMP1, cathepsin K, kininogen | IL12, lysozyme, CD68, CD106, MMP10, MMP12, cathepsin C, COX1 |
| Glycolysis | GLUT4, LGR4, PPARγ | TIGAR, LDHA★, GAPDH |
| Fibrosis | CTGF, integrin α5, PAI1 | collagen3A1, FGF1, FGF7, uPA, Tenascin C |
| Oncogenesis | AXL, EWSR1, YAP1, CEA, CRIP1, BMI1, MTA2, NF1, PDCD4 | Daxx, BACH1, ATR, Ets1, MALT1, DMBT1, BRCA2, KLF4★, Daxx, BACH1, ATR, Ets1, MALT1, DMBT1, BRCA2, KLF4, BRCA1, CRK, ZEB1, ATM |
| Senescence | sirtuin1, sirtuin3, sirtuin7 | klotho★, sirtuin6, p21 |
| Telomere signaling | TERT |  |

MiR-34a-5p\*-CEDS induced the suppression (blue) or activation (red) of signaling through protein downregulation by direct miRNA targeting (blue) or indirect response (sky blue) and protein upregulation by indirect response (red). Some miRNA target proteins were upregulated (★). The data were compared to the reports of recent manuscripts and miRDB, miRmap, miRPathDB, miRTarBase, and TargetScan websites.

The results indicate that miR-34a-5p*-CEDS induced an anti-oncogenic effect on the cells by suppressing the RAS-NFkB signaling axis, and subsequently attenuating the proliferation, angiogenesis, survival, glycolysis, and FAS- and PARP-mediated apoptosis. Conversely, it also induced an oncogenic effect by activating alternative proliferation, oncogenesis, senescence, chronic inflammation, and p53-mediated apoptosis. It can be concluded that miR-34a-5p*-CEDS exerts a divergent anti-oncogenic or oncogenic effect on RAW 264.7 cells, depending on the specific cellular context.

#### MiR-150-5p*-CEDS influenced the protein signaling in RAW 264.7 cells

RAW 264.7 cells treated with mmu-miR-150-5p sequence TCTCCCAACCCTTGTACCAGTG*-CEDS showed the characteristic protein expressions, resulting in the suppression of proliferation, cMyc/MAX/MAD network, Wnt/β-catenin signaling, p53/Rb/E2F1 signaling, SHH/PTCH/GLI signaling, epigenetic modification, protein translation, growth factors, cytodifferentiation, osteogenesis, neuromuscular differentiation, RAS signaling, NFkB signaling, angiogenesis, survival, acute inflammation, innate immunity, cellular immunity, chronic inflammation, p53-mediated apoptosis, FAS-mediated apoptosis, oncogenesis, senescence, glycolysis, and telomere signaling, as well as the activation of Notch/Jagged signaling, miRNA biogenesis, protection, PARP-mediated apoptosis, and fibrosis signaling compared to the untreated controls (Fig. 13A).

**Fig. 13.**
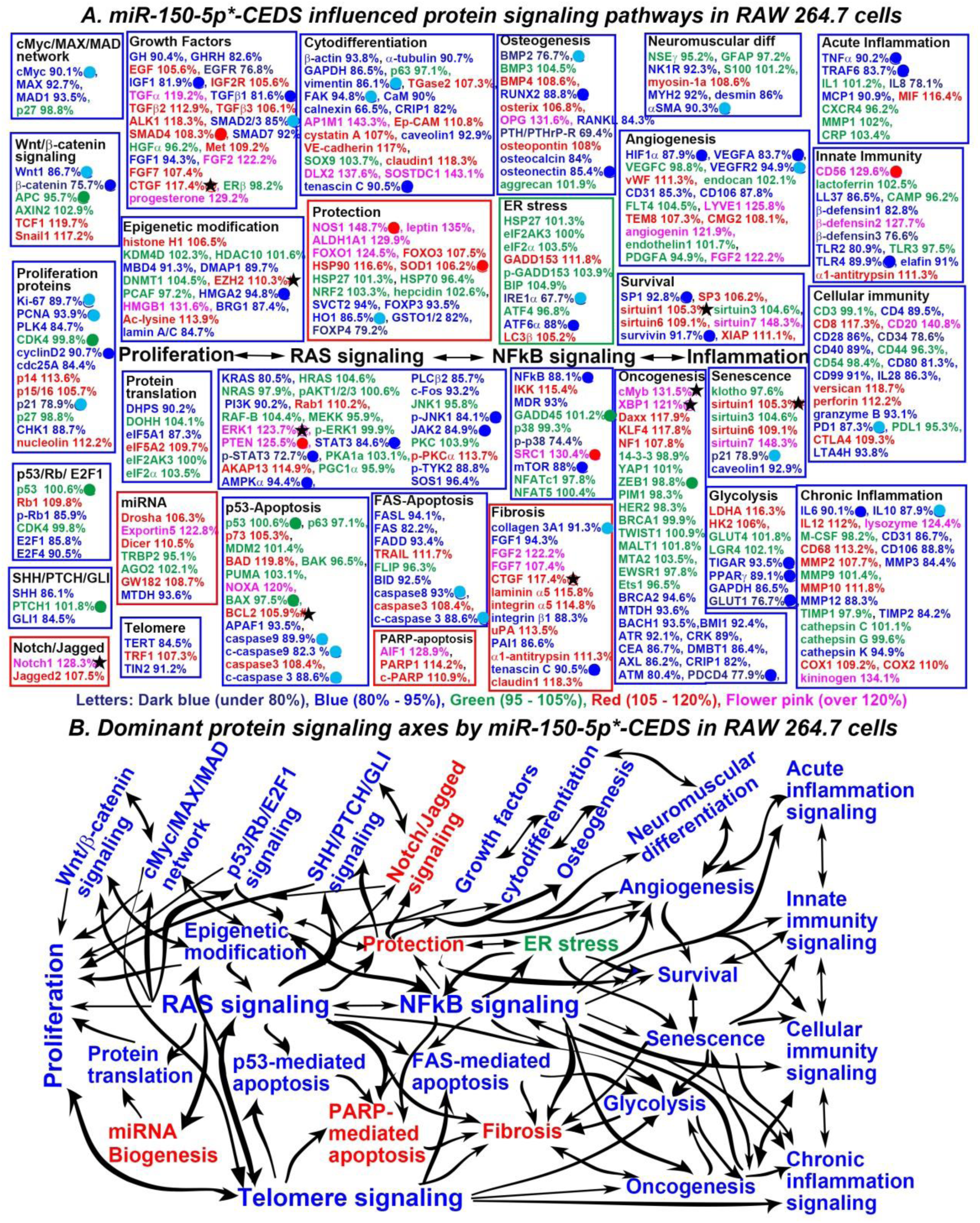
MiR-150-5p*-CEDS influenced the protein signaling (A) and axes (B) in RAW 264.7 cells. The IP- HPLC revealed proteins downregulated 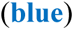, upregulated 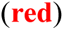, and minimally changed 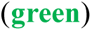 compared to the untreated controls. Dominantly suppressed 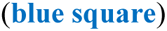 and activated 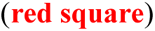 signaling. Downregulated proteins by direct targeting 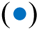 and indirect response 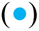, upregulated proteins by indirect response 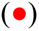, and minimal changed (±5%) proteins 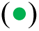. Some target proteins 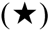 were unexpectedly upregulated contrast to the current manuscripts and data from miRDB and TargetScan websites.

**The proliferation signaling** was suppressed by downregulating miR-150-5p target protein cyclin D2 (90.7%) and miR-150-5p-responsible proteins Ki-67 (89.7%), PCNA (93.9%), and p21 (78.9%), as well as the coincidental downregulation of PLK4 (84.7%), cdc25A (84.4%), and CHK1 (88.7%), despite the upregulation of p14 (113.6%), p15/16 (105.7%), and nucleolin (112.2%).

**The cMyc/MAX/MAD network** was suppressed by downregulating miR-150-5p-responsible protein cMyc (89.7%) and coincidentally downregulating MAX (92.7%) and MAD1 (93.5%).

**The Wnt/β-catenin signaling** was suppressed by downregulating miR-150-5p target protein β-catenin (75.7%) and miR-150-5p-responsible protein Wnt1 (86.7%), despite the upregulation TCF1 (119.7%) and Snail1 (117.2%). TCF1 could be upregulated through the alternative signaling.

**The p53/Rb/E2F1 signaling** was suppressed by downregulating p-Rb1 (85.0%), E2F1 (85.8%), and E2F4 (90.5%) and coincidentally upregulating Rb1 (109.8%).

**The SHH/PTCH/GLI signaling** was suppressed by downregulating SHH (86.1%) and GLI1 (84.5%).

**The protein translation signaling** was suppressed by downregulating DHPS (90.2%), eIF5A1 (87.3%), despite the upregulation of eIF5A2 (109.7%).

**The growth factor signaling** were suppressed by downregulating miR-150-5p target proteins IGF1 (81.9%) and TGFβ1 (81.6%) and miR-150-5p-responsible protein SMAD2/3 (85%), as well as coincidentally downregulating GH (90.4%), GHRH (82.6%), EGFR (76.8%), SMAD7 (92%), and FGF1 (94.3%), despite the upregulation of miR-150-5p-responsible protein SMAD4 (108.3%) and coincidental upregulation of EGF (105.6%), IGF2R (105.6%), TGFα (119.2%), TGFβ2 (112.9%), TGFβ3 (106.1%), ALK1 (118.3%), Met (109.2%), FGF2 (122.2%), FGF7 (107.4%), CTGF (117.4%), and progesterone (129.2%). CTGF is a miR-150-5p target protein, but it was unexpectedly upregulated compared to the reference data.

**The cytodifferentiation signaling** partially were suppressed by downregulating miR-150-5p target protein tenascin C (90.5%) and miR-150-5p-responsible proteins vimentin (86.1%) and FAK (94.8%) and coincidentally downregulating β-actin (93.8%), α-tubulin (90.7%), GAPDH (86.5%), CaM (90%), calnexin (66.5%), CRIP1 (82%), and caveolin1 (92.9%), despite the upregulation of TGase2 (107.3%), AP1M1 (143.3%), Ep-CAM (110.8%), cystatin A (107%), VE-cadherin (117%), claudin1 (118.3%), DLX2 (137.6%), and SOSTDC1 (143.1%).

**The osteogenesis signaling** was inactivated by downregulating miR-150-5p target proteins RUNX2 (88.8%), osteonectin (85.4%), and miR-150-5p-responsible protein BMP2 (76.7%), as well as coincidentally downregulating RANKL (84.3%), PTH/PTHrP-R (69.4%), and osteocalcin (84%), despite the upregulation of BMP4 (108.6%), osterix (106.8%), OPG (131.6%), and osteopontin (108%).

**The neuromuscular differentiation signaling** were inactivated by downregulating miR-150-5p- rsponsible protein αSMA (90.3%) and coincidentally downregulating NK1R (92.3%), MYH2 (92%), and desmin (86%), despite the upregulation of myosin-1a (108.6%).

**The RAS signaling** was suppressed by downregulating miR-150-5p target proteins STAT3 (84.6%), p- STAT3 (72.7%), AMPKα (94.4%), p-JNK1 (84.1%), and JAK2 (84.9%), as well as coincidentally downregulating KRAS (80.5%), PI3K (90.2%), PLCβ2 (85.7%), c-Fos (93.2%), p-TYK2 (88.8%), and SOS1 (96.4%), despite the upregulation of Rab1 (110.2%), ERK1 (123.7%), AKAP13 (114.9%), and p-PKC1α (113.7%). ERK1 is a miR-150-5p target protein, but it was unexpectedly upregulated compared to the reference data.

**The NFkB signaling** was suppressed by downregulating miR-150-5p target proteins NFkB (88.1%) and mTOR (88%), as well as coincidentally downregulating MDR (93%) and p-p38 (74.4%), and upregulating IKK (115.4%), despite the upregulation of miR-150-5p-responsible protein SRC1 (130.4%).

**The angiogenesis signaling** was suppressed by downregulating miR-150-5p target proteins HIF1α (87.9%), VEGFA (83.7%), and miR-150-5p-responsible protein VEGFR2 (94.9%), as well as coincidentally downregulating CD31 (85.3%) and CD106 (87.8%), despite the upregulation of vWF (111.3%), LYVE1 (125.8%), TEM8 (107.3%), CMG2 (108.1%), angiogenin (121.9%), and FGF2 (122.2%).

**The survival signaling** was suppressed by downregulating miR-150-5p target proteins SP1 (92.8%) and survivin (91.7%), despite the upregulation of SP3 (106.2%), sirtuin1 (105.3%), sirtuin6 (109.1%), sirtuin7 (148.3%), and XIAP (111.1%). Sirtuin1 is a miR-150-5p target protein, but it was unexpectedly upregulated compared to the reference data.

**The acute inflammation signaling** was suppressed by downregulating miR-150-5p target proteins TNFα (90.2%) and TRAF6 (83.7%), as well as coincidentally downregulating IL1 (78.1%) and MCP1 (90.9%), despite the upregulation of MIF (116.4%).

**The innate immunity signaling** appears to be inactivated by downregulating miR-150-5p target protein TLR4 (89.9%) and coincidentally downregulating LL37 (86.5%), β-defensin1 (82.8%), β-defensin3 (76.6%), TLR2 (80.9%), and elafin (91%), despite the upregulation of miR-150-5p-responsible protein CD56 (129.6%) and coincidental upregulation of β-defensin2 (127.7%) and α1-antitrypsin (111.3%).

**The cellular immunity signaling** was suppressed by downregulating miR-150-5p-responsible protein PD1 (87.3%), as well as coincidentally downregulating CD4 (89.5%), CD28 (86%), CD34 (78.6%), CD40 (89%), CD80 (81.3%), CD99 (91%), IL28 (86.3%), granzyme B (93.1%), and LTA4H (93.8%), despite the upregulation of CD8 (117.3%), CD20 (140.8%), versican (118.7%), perforin (112.2%), and CTLA4 (109.3%).

**The chronic inflammation signaling** was suppressed by downregulating miR-150-5p target proteins IL6 (90.1%) and miR-150-5p-responsible protein IL10 (87.9%), as well as coincidentally downregulating CD31 (86.7%), CD106 (88.8%), MMP3 (84.4%), MMP12 (88.3%), TIMP2 (84.2%), and cathepsin K (94.9%), despite the upregulation of IL12 (112%), lysozyme (124.4%), CD68 (113.2%), MMP2 (107.7%), COX1 (109.2%), COX2 (110%), and kininogen (134.1%).

**The p53-mediated apoptosis signaling** was suppressed by downregulating miR-150-5p-responsible proteins caspase9 (89.9%), c-caspase9 (82.3%), and c-caspase3 (88.6%), as well as coincidentally downregulating APAF1(93.5%), despite the upregulation of p73 (105.3%), BAD (119.8%), NOXA (120%), BCL2 (105.9%), and caspase3 (108.4%). BCL2 is a miR-150-5p target protein, but it was unexpectedly upregulated compared to the reference data.

**The FAS-mediated apoptosis signaling** was inhibited by downregulating miR-150-5p-responsible proteins caspase8 (93%) and c-caspase3 (88.6%), as well as coincidentally downregulating FASL (94.1%), FAS (82.2%), FADD (93.4%), and BID (92.5%), despite the upregulation of TRAIL (111.7%) and caspase3 (108.4%).

**The senescence signaling** was suppressed by downregulating miR-150-5p-responsible protein p21 (78.9%) and coincidentally downregulating caveolin1 (92.9%), despite the upregulation of sirtuin1 (105.3%), sirtuin6 (109.1%), and sirtuin7 (148.3%). Sirtuin1 is a miR-150-5p target protein, but it was unexpectedly upregulated compared to the reference data.

**The glycolysis signaling** was suppressed by downregulating miR-150-5p target protein TIGAR (93.5%), PPARγ (89.1%), and GLUT1 (76.7%), as well as coincidentally downregulating GAPDH (86.5%), despite the upregulation of LDHA (116.3%) and HK2 (106%).

**The telomere signaling** was suppressed by downregulating TERT (84.5%) and TIN2 (91.2%), despite the upregulation of TRF1 (107.3%).

Conversely, **the Notch/Jagged signaling** was activated by upregulating Notch1 (128.3%) and Jagged2 (107.5%). Notch1 is a miR-150-5p target protein, but it was unexpectedly upregulated compared to the reference data.

**The miRNA biogenesis signaling** was activated by upregulating Drosha (106.3%), Exportin5 (122.8%), Dicer (110.5%) and GW182 (108.7%), and downregulating MTDH (93.6%) coincidentally.

**The protection signaling** was activated by upregulating miR-150-5p-responsible proteins NOS1 (148.7%) and SOD1 (106.2%), as well as coincidentally upregulating leptin (135%), ALDH1A1 (129.9%), FOXO1 (124.5%), FOXO3 (107.5%), and HSP90 (116.6%), despite the downregulation of miR-150-5p-responsible protein HO1 (86.5%) and coincidental downregulation of SVCT2 (94%), FOXP3 (93.5%), GSTO1/2 (82%), and FOXP4 (79.2%).

**The PARP-mediated apoptosis signaling** was activated by upregulating AIF1 (128.9%), PARP1 (114.2%), and c-PARP (110.9%).

**The fibrosis signaling** appears to be activated by upregulating FGF2 (122.2%), CTGF (117.4%), lamininα5 (115.8%), integrin α5 (114.9%), uPA (113.5%), α1-antitrypsin (111.3%), and claudin1 (118.3%), despite the downregulation of miR-150-5p target protein tenascin C (90.5%) and miR-150-5p-responsible protein collagen 3A1 (91.3%), as well as coincidentally downregulating FGF1 (94.3%), integrin β1 (88.3%), and PAI1 (86.6%) and upregulating FGF7 (107.4%). CTGF is a miR-150-5p target protein, but it was unexpectedly upregulated compared to the reference data.

In addition, miR-150-5p*-CEDS significantly influenced **the epigenetic modification signaling** by downregulating miR-150-5p target proteins HMGA2 (94.8%), as well as the coincidentally downregulating MBD4 (91.3%), DMAP1 (89.7%), BRG1 (87.4%), and lamin A/C (84.7%), while it upregulated histone H1 (106.5%), EZH2 (110.3%), HMGB1 (131.6%), and Ac-lysine (113.9%). EZH2 is a miR-150-5p target protein, but it was unexpectedly upregulated compared to the reference data. Consequently, miR-150-5p*-CEDS induced a trend towards a reduction in the methylation of histones and DNAs, and attenuation of transcriptional repression.

MiR-150-5p*-CEDS markedly suppressed **the oncogenesis signaling** by downregulating miR-150-5p target proteins PDCD4 (77.9%) and coincidentally downregulating BRCA2 (94.6%), MTDH (93.6%), BACH1 (93.5%), BMI1 (92.4%), ATR (92.1%), CRK (89%), CEA (86.7%), DMBT1 (86.4%), AXL (86.2%), CRIP1 (82%), and ATM (80.4%), while it partly enhanced **the oncogenesis signaling** by upregulating cMyb (131.5%), XBP1 (121%), KLF4 (117.8%), NF1 (107.8%), and Daxx (117.9%). cMyb and XBP1 are miR-150-5p target proteins, but they were unexpectedly upregulated compared to the reference data. Among the 28 oncoproteins, 12 were found to be underexpressed, five were overexpressed, and 11 exhibited minimal change in expression compared to the untreated controls.

Consequently, miR-150-5p*-CEDS had a profound impact on the entire protein signaling in RAW 264.7 cells. MiR-150-5p*-CEDS suppressed the RAS-NFkB signaling axis, subsequently inactivating the majority of proliferation-related signaling axis, growth and cytodifferentiation signaling axis, ER stress-angiogenesis-survival-senescence signaling axis, and so on. MiR-150-5p*-CEDS also impacted the inflammation-immunity signaling axis, glycolysis-oncogenesis-chronic inflammation signaling axis, telomere-epigenetic modification signaling axis, and p53- and FAS-mediated apoptosis signaling. Conversely, it activated protection- Notch/Jagged signaling axis, PARP-mediated apoptosis-fibrosis signaling axis, and miRNA biogenesis signaling (Fig 13B, Table 6).

**Table 6.** Protein expressions in the RAW 264.7 cells treated with miR-150-5p*-CEDS.

| Signaling | Downregulated Proteins | Upregulated Proteins |
| --- | --- | --- |
| Proliferation | Ki-67, PCNA, PLK4, cyclin D2, cdc25A, p21, CHK1 | P14, p15/16, nucleolin |
| cMyc/MAX/MAD network | cMyc, MAX, MAD1 |  |
| Wnt/β-catenin signaling | Wnt1, β-catenin | TCF1, Snail |
| p53/Rb/E2F1 signaling | p-Rb1, E2F1, E2F4 | Rb1 |
| SHH/PTCH/GLI signaling | SHH, GLI1 |  |
| Notch/Jagged signaling |  | Notch1 ★, Jagged2 |
| Epigenetic modification | MBD4, DMAP1, HMG A2, BRG1, lamin A/C | Histone H1, EZH2 ★, HMGB1, Ac-lysine |
| Protein translation | DHPS, eIF5A1 | eIF5A2 |
| miRNA biogenesis | MTDH | Drosha, Exportin5, Dicer, GW182 |
| Growth factors | GH, GHRH, EGFR, IGF1, TGFβ1, SMAD2/3, SMAD7, FGF1 | EGF, IGF2R, TGFα, TGFβ2, TGFβ3, ALK1, SMAD4, Met, FGF2, FGF7, CTGF ★, progesterone |
| Cytodifferentiation | β-actin, α-tubulin, GAPDH, vimentin, FAK, CaM, calnexin, CRIP1, caveolin1, tenascin C | TGase2, AP1M1, Ep-CAM, cystatin A, VE-cadherin, claudin1, DLX2, SOSTDC1 |
| Osteogenesis | BMP2, RUNX2, RANKL, PTH/PTHrPR, osteocalcin, osteonectin | BMP4, osterix, OPG, osteopontin |
| Neuromuscular differentiation | NK1R, MYH2, desmin, αSMA | myosin-1a |
| RAS signaling | KRAS, PI3K, STAT3, p-STAT3, AMPKα, PLCβ2, c-Fos, p-JNK1, JAK2, p-TYK2, SOS1 | Rab1, ERK1 ★, PTEN, AKAP13, p-PKC1α |
| NFkB signaling | NFkB, MDR, p-p38, mTOR | IKK, SRC1 |
| p53-mediated apoptosis | APAF1, caspase9, c-caspase9, c-caspase3 | p73, BAD, NOXA, BCL2 ★, caspase3 |
| FAS-mediated apoptosis | FASL, FAS, FADD, BID, caspase9, c-caspase3 | TRAIL, caspase3 |
| PARP-mediated apoptosis |  | AIF1, PARP1, c-PARP |
| Protection | SVCT2, FOXP3, HO1, GSTO1/2, FOXP4 | NOS1, leptin, ALDH1A1, FOXO3, HSP90, SOD1, |
| ER stress | IRE1α, ATF6α | GADD153, LC3β |
| Angiogenesis | HIF1α, VEGFA, VEGFR2, CD31, CD106, | vWF, LYVE1, TEM8, CMG2, angiogenin, FGF2 |
| Survival | SP1, survivin, | SP3, sirtuin1 ★, sirtuin6, sirtuin7, XIAP |
| Acute inflammation | TNFα, TRAF6, IL8, MCP1 | MIF |
| Innate immunity | LL37, β-defensin1, β-defensin3, TLR2, TLR4, elafin, | CD56, β-defensin2, α1-antitrypsin |
| Cellular immunity | CD4, CD28, CD34, CD40, CD80, CD99, IL28, granzyme B, PD1, LTA4H | CD8, CD20, versican, perforin, CTLA4 |
| Chronic inflammation | IL6, IL10, CD31, CD106, MMP3, MMP12, TIMP2, cathepsin K | IL12, lysozyme, CD68, MMP2, MMP10, COX1, COX2, kininogen |
| Glycolysis | TIGAR, PPARγ, GAPDH, GLUT1 | LDHA, HK2 |
| Fibrosis | collagen 3A1, FGF1, integrin β1, PAI1, tenascin C | FGF2, FGF7, CTGF ★, laminin α5, integrin α5, uPA, α1-antitrypsin, claudin1 |
| Oncogenesis | DMBT1, CRK, MTDH, CEA, AXL, BACH1, BMI1, BRCA2, ATR, ATM, CRIP1, PDCD4 | cMyb ★, XBP1 ★, KLF4, NF1, Daxx |
| Senescence | p21, caveolin1 | sirtuin1 ★, sirtuin6, sirtuin7 |
| Telomere signaling | TERT, TIN2 | TRF1 |

miR-150-5p\*-CEDS induced the suppression (blue) or activation (red) of signaling through protein downregulation by direct miRNA targeting (blue) or indirect response (sky blue) and protein upregulation by indirect response (red). Some miRNA target proteins were upregulated (★). The data were compared to the reports of recent manuscripts and miRDB, miRmap, miRPathDB, miRTarBase, and TargetScan websites.

The results demonstrate that miR-150-5p*-CEDS exerts a potent anti-oncogenic effect on cells by suppressing proliferation, angiogenesis, survival, senescence, glycolysis, oncogenesis, chronic inflammation, and telomere instability. Conversely, it increased the PARP-mediated apoptosis and enhanced the ROS-Notch signaling. Additionally, it is worthy to note that miR-150-5p*-CEDS attenuated innate and cellular immunity, reduced major p53- and FAS-mediated apoptosis, and resulted in fibrosis signaling.

#### MiR-155-5p*-CEDS influenced the protein signaling in RAW 264.7 cells

RAW 264.7 cells treated with mmu-miR-155-5p sequence TTAATGCTAATCGTGATAGGGGTT*- CEDS showed the characteristic protein expressions, resulting in the suppression of cMyc/MAX/MAD network, Wnt/β-catenin signaling, Notch/Jagged signaling, epigenetic modification, miRNA biogenesis, osteogenesis, NFkB signaling, ER stress, angiogenesis, acute inflammation, chronic inflammation, p53-mediated apoptosis, PARP-mediated apoptosis, oncogenesis, and glycolysis signaling, as well as the activation of proliferation, p53/Rb/E2F1 signaling, SHH/PTCH/GLI signaling, growth factors, cytodifferentiation, neuromuscular differentiation, RAS signaling, protection, survival, innate immunity, cellular immunity, FAS-mediated apoptosis, fibrosis, and senescence compared to the untreated controls (Fig. 14A).

**Fig. 14.**
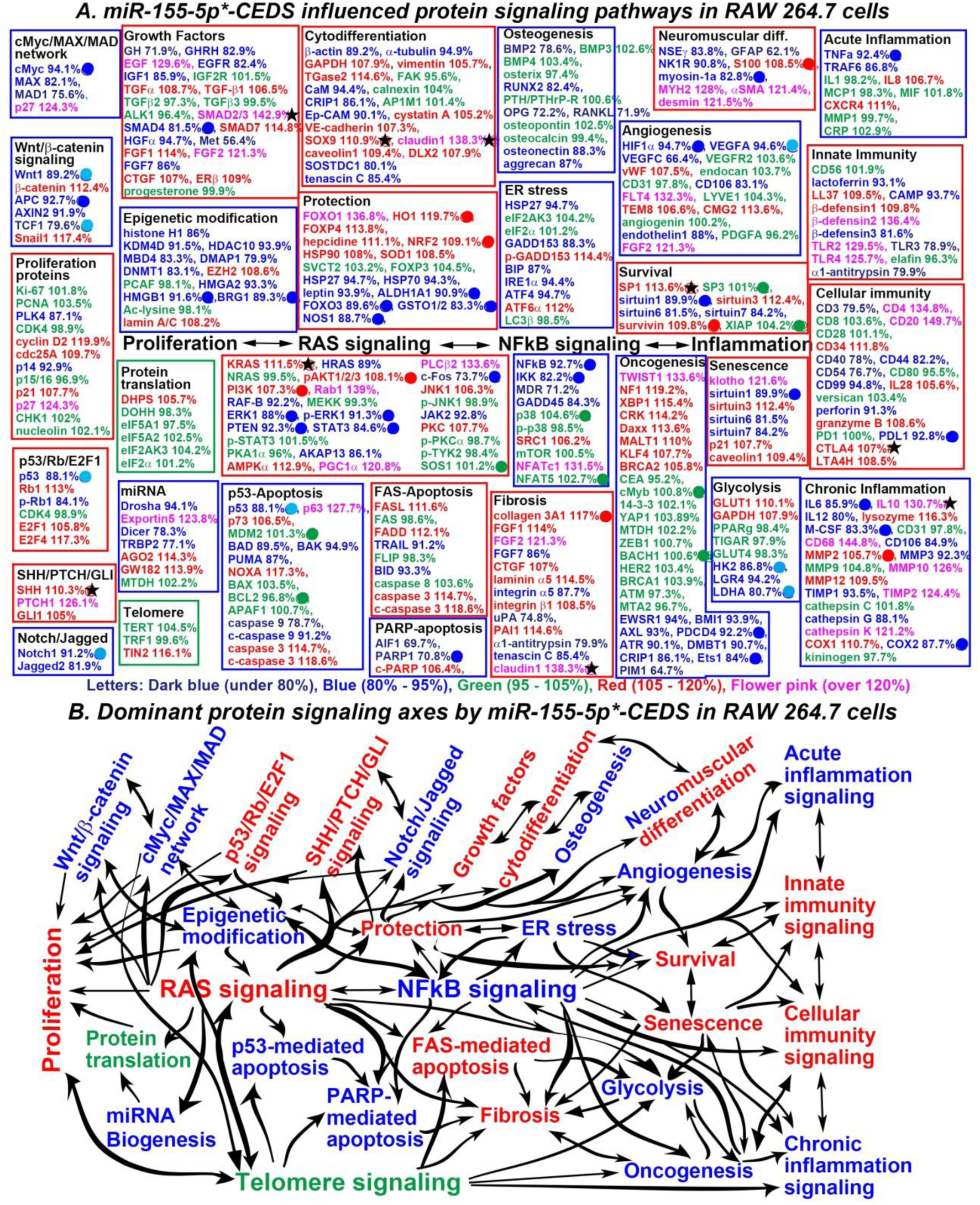
MiR-155-5p*-CEDS influenced the protein signaling (A) and axes (B) in RAW 264.7 cells. The IP- HPLC revealed proteins downregulated 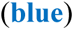, upregulated 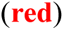, and minimally changed 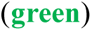 compared to the untreated controls. Dominantly suppressed 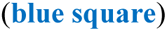 and activated 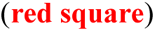 signaling. Downregulated proteins by direct targeting 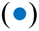 and indirect response 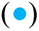, upregulated proteins by indirect response 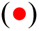, and minimal changed (±5%) proteins 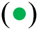. Some target proteins 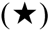 were unexpectedly upregulated contrast to the current manuscripts and data from miRDB and TargetScan websites.

**The cMyc/MAX/MAD network** was suppressed by downregulating miR-155-5p target protein cMyc (94.1%) and coincidentally downregulating MAX (82.1%) and MAD1 (75.6%).

**The Wnt/β-catenin signaling** was suppressed by downregulating miR-155-5p target protein APC (92.7%) and miR-155-5p-responsible proteins Wnt1 (89.2%) and TCF1 (79.6%), and coincidental downregulation of AXIN2 (91.9%), despite the upregulation of β-catenin (112.4%) and Snail1 (117.4%).

**The Notch/Jagged signaling** was suppressed by downregulating miR-155-5p-responsible protein Notch1 (91.2%) and coincidentally downregulating Jagged2 (81.9%).

**The miRNA biogenesis signaling** was inactivated by downregulating Dicer (78.3%) and TRBP2 (77.1%). despite the upregulation of Exportin5 (123.8%), AGO2 (114.3%), and GW182 (113.9%).

**The osteogenesis signaling** was suppressed by downregulating BMP2 (78.6%), RUNX2 (82.4%), OPG (72.2%), RANKL (71.9%), osteonectin (88.3%), and aggrecan (87%).

**The NFkB signaling** was suppressed by downregulating miR-155-5p target proteins NFkB (92.7%), IKK (82.2%), as well as coincidentally downregulating MDR (71.2%), GADD45 (84.3%), despite the upregulation of SRC1 (106.2%) and NFATc1 (131.5%).

**The ER stress signaling** was suppressed by downregulating HSP27 (94.7%), GADD153 (88.3%), BIP (87%), IRE1α (94.4%), ATF4 (94.7%), despite the upregulation of p-GADD153 (114.4%) and ATF6α (112%).

**The angiogenesis signaling** was suppressed by downregulating miR-155-5p target protein HIF1α (94.7%) and miR-155-5p-responsible protein VEGFA (94.6%), as well as coincidentally downregulating VEGFC (66.4%), CD106 (83.1%), and endothelin1 (88%), despite the upregulation of vWF (66.4%), FLT4 (132.3%), TEM8 (106.6%), CMG2 (113.6%), and FGF2 (121.3%).

**The acute inflammation signaling** was suppressed by downregulating miR-155-5p target protein TNFα (92.4%) and TRAF6 (86.8%), despite the upregulation of IL8 (106.7%) and CXCR4 (111%).

**The chronic inflammation signaling** was suppressed by downregulating miR-155-5p target proteins IL6 (85.9%), M-CSF (83.3%), and COX2 (87.7%), and coincidentally downregulating IL12 (80%), CD106 (84.9%), TIMP1 (93.5%), and cathepsin G (88.1%), despite the upregulation of IL10 (130.7%), lysozyme (116.3%), CD68 (144.8%), MMP2 (105.7%), MMP10 (126%), MMP12 (109.5%), TIMP2 (124.4%), cathepsin K (121.2%), and COX1 (110.7%). IL10 is a miR-155-5p target protein, but it was unexpectedly upregulated compared to the reference data.

**The p53-mediated apoptosis signaling** appears to be suppressed by downregulating miR-155-5p- responsible protein p53 (88.1%), as well as coincidentally downregulating BAD (89.5%), BAK (94.9%), PUMA (87%), caspase9 (78.7%), and c-caspase9 (91.2%), despite the upregulation of p63 (127.7%), p73 (106.5%), NOXA (117.3%), caspase3 (114.7%), and c-caspase3 (118.6%).

**The PARP-mediated apoptosis signaling** appears to be suppressed by downregulating miR-155-5p- responsible proteins PARP1 (70.8%) and coincidentally downregulating AIF1 (69.7%), despite the slight upregulation of c-PARP (106.4%).

**The glycolysis signaling** was suppressed by downregulating miR-155-5p-responsible protein HK2 (86.8%) and LDHA (80.7%) and coincidentally downregulating LGR4 (94.2%), despite the slight upregulation of GLUT1 (110.1%) and GAPDH (107.9%).

Conversely, **the proliferation signaling** appears to be enhanced by the upregulation of cyclin D2 (119.9%), cdc25A (109.7%), and the coincidental downregulation of p14 (92.9%), despite the downregulation of PLK4 (87.1%) and the upregulation of p21 (107.7%) and p27 (124.3%).

**The p53/Rb/E2F1 signaling** appears to be activated by downregulating miR-155-5p-responsible protein p53 (88.1%) and upregulating E2F1 (105.8%), despite the upregulation of Rb1 (113%) and E2F4 (117.3%), as well as the downregulation of p-Rb1 (84.1%).

**The SHH/PTCH/GLI signaling** was enhanced by upregulation of SHH (110.3%), PTCH1 (126.1%), and GLI1 (105%). SHH is a miR-155-5p target protein, but it was unexpectedly upregulated compared to the reference data.

**The growth factor signaling** appears to be activated by upregulating EGF (129.6%), TGFα (108.7%), TGFβ1 (106.5%), SMAD2/3 (142.9%), SMAD7 (114.8%), FGF1 (114%), FGF2 (121.3%), CTGF (107%), and ERβ (109%), and coincidentally downregulating FGF7 (86%), despite the downregulation of miR-155-5p target protein SMAD4 (81.5%) and coincidental downregulation of GH (71.9%), GHRH (82.9%), IGF1 (85.9%), HGFα (94.7%), and Met (56.4%). SMAD4 is a miR-155-5p target protein, but it was unexpectedly upregulated compared to the reference data.

**The cytodifferentiation signaling were** enhanced by upregulating GAPDH (107.9%), vimentin (105.7%), TGase2 (114.6%), cystatin A (105.2%), VE-cadherin (107.3%), SOX9 (110.9%), claudin1 (138.3%), caveolin1 (109.4%), and DLX2 (107.9%), despite the downregulation of β-actin (89.2%), α-tubulin (94.9%), CaM (94.4%), CRIP1 (86.1%), Ep-CAM (90.1%), SOSTDC1 (80.1%), and tenascin C (85.4%). SOX9 and claudin1 are miR-155-5p target proteins, but they were unexpectedly upregulated compared to the reference data.

**The neuromuscular differentiation signaling** were activated by upregulating miR-155-5p-responsible protein S100 (108.5%) and coincidentally upregulating MYH2 (128%), αSMA (121.4%), and desmin (121.5%), despite the downregulation of miR-155-5p target protein myosin-1a (82.8%) and coincidental downregulation of NSEγ (83.8%), GFAP (62.1%), and NK1R (90.8%).

**The RAS signaling** appears to be activated by upregulating miR-155-5p-responsible proteins pAKT1/2/3 (108.1%) and PI3K (107.3%) and coincidentally upregulating KRAS (111.5%), Rab1 (139%), AMPKα (112.9%), PGC1α (120.8%), PLCβ2 (133.6%), JNK1 (106.3%), and PKC (107.7%), despite the downregulation of miR-155-5p target proteins ERK1 (88%), p-ERK1 (91.3%), PTEN (92.3%), STAT3 (84.6%), and c-Fos (73.7%), as well as the coincidental downregulation of HRAS (89%), AKAP13 (86.1%), and JAK2 (92.8%). KRAS is a miR-155-5p target protein, but it was unexpectedly upregulated compared to the reference data.

**The protection signaling** appears to be activated by upregulating miR-155-5p-responsible proteins HO1 (119.7%) and NRF2 (109.1%) and coincidentally upregulating FOXO1 (136.8%), FOXP4 (113.8%), hepcidine (111.1%), HSP90 (108%), SOD1 (108.5%), despite the downregulation of miR-155-5p target proteins ALDH1A1 (90.9%), FOXO3 (89.6%), GSTO1/2 (83.3%), and NOS1 (88.7%) and coincidental downregulation of HSP27 (94.7%), HSP70 (94.3%), and leptin (93.9%).

**The survival signaling** was activated by upregulating miR-155-5p-responsible protein survivin (109.8%) and coincidentally upregulating SP1 (113.6%) and sirtuin3 (112.4%), despite the downregulation of sirtuin1 (89.9%), sirtuin6 (81.5%), and sirtuin7 (84.2%). SP1 is a miR-155-5p target protein, but it was unexpectedly upregulated compared to the reference data.

**The innate immunity signaling** appears to be activated by upregulating LL37 (109.5%), β-defensin1 (109.8%), β-defensin2 (136.4%), TLR2 (129.5%), and TLR4 (125.7%), despite the downregulation of lactoferrin (93.1%), CAMP (93.7%), β-defensin3 (81.6%), TLR3 (78.9%), and α1-antitrypsin (79.9%).

**The cellular immunity signaling** was slightly activated by downregulating miR-155-5p target protein PDL1 (92.8%) and coincidentally upregulating CD4 (134.8%), CD20 (149.7%), CD34 (111.8%), IL28 (105.6%), granzyme B (108.6%), CTLA4 (107%), and LTA4H (108.5%). CTLA4 is a miR-155-5p target protein, but it was unexpectedly upregulated compared to the reference data.

**The FAS-mediated apoptosis signaling** was activated by upregulating FASL (111.6%), FADD (112.1%), caspase3 (114.7%), and c-caspase3 (118.6%), despite the downregulation of TRAIL (91.2%) and BID (93.3%).

**The fibrosis signaling** was enhanced by upregulating miR-155-5p target protein collagen 3A1 (117%), as well as coincidental upregulation of FGF1 (114%), FGF2 (121.3%), CTGF (107%), laminin α5 (114.5%), integrin β1 (108.5%), PAI1 (114.6%), and claudin1 (138.3%) and downregulating FGF7 (86%), despite the downregulation of integrin α5 (87.7%), uPA (74.8%), α1-antitrypsin (79.9%), and tenascin C (85.4%). Claudin1 is a miR-155-5p target protein, but it was unexpectedly upregulated compared to the reference data.

**The senescence signaling** was activated by upregulating klotho (121.6%), sirtuin3 (112.4%), p21 (107.7%), and caveolin1 (109.4%), despite the downregulation of miR-155-5p target protein sirtuin1 (89.9%) and the coincidental downregulation of sirtuin6 (81.5%) and sirtuin7 (84.2%).

In addition, miR-155-5p*-CEDS markedly influenced **the epigenetic modification signaling** by downregulating miR-155-5p target proteins HMGB1 (91.6%) and BRG1 (89.3%), as well as the coincidental downregulation of histone H1 (86%), KDM4D (91.5%), HDAC10 (93.9%), MBD4 (83.3%), DMAP1 (79.9%), DNMT1 (83.1%), and HMGA2 (93.3%), while it upregulated EZH2 (108.6%) and lamin A/C (108.2%). Consequently, miR-155-5p*-CEDS induced a trend towards a reduction in the methylation of histones and DNAs, and attenuation of transcriptional repression.

MiR-155-5p*-CEDS significantly suppressed **the oncogenesis signaling** by downregulating miR-155-5p target proteins PDCD4 (92.2%) and Ets (84%), as well as coincidentally downregulating EWSR (94%), BMI (93.9%), AXL (93%), ATR (90.1%), DMBT1 (90.7%), CRIP1 (86.1%), and PIM1 (64.7%), while it enhanced **the oncogenesis signaling** by upregulating TWIST1 (133.6%), NF1 (119.2%), XBP1 (115.4%), CRK (114.2%), Daxx (113.6%), MALT1 (110%), KLF4 (107.7%), and BRCA2 (105.8%). Among the 28 oncoproteins, nine were found to be underexpressed, eight were overexpressed, and 11 exhibited minimal change in expression compared to the untreated controls.

Consequently, miR-155-5p*-CEDS exerted a profound influence on the entire protein signaling in RAW 264.7 cells. MiR-155-5p*-CEDS suppressed the NFkB signaling, inactivating the epigenetic modification- cMyc/MAX/MAD network-Wnt/β-catenin signaling axis, ER stress-angiogenesis-acute inflammation, and glycolysis-oncogenesis-chronic inflammation signaling axis. Conversely, it enhanced the RAS signaling, and subsequently activated proliferation signaling axis, protection-survival-senescence signaling axis, and FAS-mediated apoptosis-fibrosis signaling axis (Fig. 14B, Table 7).

**Table 7.** Protein expressions in the RAW 264.7 cells treated with miR-155-5p*-CEDS.

| Signaling | Downregulated Proteins | Upregulated Proteins |
| --- | --- | --- |
| **Proliferation** | PLK4, p14 | cyclin D2, cdc25A, p21, p27 |
| **cMyc/MAX/MAD network** | **cMyc**, MAX, MAD1 | p27 |
| **Wnt/β-catenin signaling** | **Wnt1**, APC, AXIN2, **TCF1** | β-catenin, Snail |
| **p53/Rb/E2F1 signaling** | **p53**, p-Rb1, | Rb1, E2F1, E2F4 |
| **SHH/PTCH/GLI signaling** |  | **SHH**★, PTCH1, GLI1 |
| **Notch/Jagged signaling** | **Notch1**, Jagged2 |  |
| **Epigenetic modification** | histone H1, KDM4D, HDAC10, MBD4, CMAP1, DNMT1, HMG A2, **HMG B1**, **BRG1** | EZH2, lamin A/C |
| **Protein translation** |  | DHPS |
| **miRNA biogenesis** | Drosha, Dicer, TRBP2 | Exportin5, AGO2, GW182 |
| **Growth factors** | GH, GHRH, EGFR, IGF1, **SMAD4**, HGFα, Met, FGF7 | EGF, TGFα, TGFβ1, **SMAD2/3**★, SMA7, FGF1, FGF2, CTGF, ERβ |
| **Cytodifferentiation** | β-actin, α-tubulin, CaM, CRIP1, Ep-CAM, SOSTDC1, tenascin C | GAPDH, vimentin, TGase2, cystatin A, VE-cadherin, **SOX9**★, **claudin1**★, caveolin1, DLX2 |
| **Osteogenesis** | BMP2, RUNX2, OPG, RANKL, osteonectin, aggrecan |  |
| **Neuromuscular differentiation** | NSEγ, GFAP, NK1R, **myosin-1a** | **S100**, MYH2, αSMA, desmin |
| **RAS signaling** | HRAS, RAF-B, **ERK1**, **p-ERK1**, PTEN, **STAT3**, AKAP13, **c-Fos**, JAK2 | **KRAS**★, **pAKT1/2/3**, **PI3K**, Rab1, AMPKα, PGC1α, PLCβ2, JNK1, PKC |
| **NFκB signaling** | **NFκB**, **IKK**, MDR, GADD45 | SRC1, NFATc1 |
| **p53-mediated apoptosis** | **p53**, BAD, BAK, PUMA, caspase9<br>c-caspase9 | p63, p73, NOXA, caspase3, c-caspase3 |
| **FAS-mediated apoptosis** | TRAIL, BID, | FASL, FADD, caspase3, c-caspase3 |
| **PARP-mediated apoptosis** | AIF1, **PARP1** | c-PARP1 |
| **Protection** | HSP27, HSP70, leptin, **ALDH1A1**, **FOXO3**, **GSTO1/2**, **NOS1** | FOXO1, **HO1**, FOX P4, hepcidin, **NRF2**, HSP90, SOD1 |
| **ER stress** | HSP27, GADD153, BIP, IRE1α, ATF4 | p-GADD153, ATF6α |
| **Angiogenesis** | **HIF1α**, **VEGFA**, VEGFC, CD106, endothelin1 | vWF, FLT4, TEM8, CMG2, FGF2 |
| **Survival** | **sirtuin1**, sirtuin6, sirtuin7, | **SP1**★, sirtuin3, **survivin**, |
| **Acute inflammation** | **TNFα**, TRAF6 | IL1, CXCR4 |
| **Innate immunity** | lactoferrin, CAMP, β-defensin3, TLR3, α1-antitrypsin | LL37, β-defensin1, β-defensin2, TLR2, TLR4 |
| **Cellular immunity** | CD3, CD40, CD44, CD54, CD99, perforin, **PDL1** | CD4, CD20, CD34, IL28, granzyme B, **CTLA4**★, LTA4H |
| **Chronic inflammation** | **IL6**, IL12, **M-CSF**, CD106, MMP3, TIMP1, cathepsin G, **COX2** | **IL10**★, lysozyme, CD68, **MMP2**, MMP10, MMP12, TIMP2, cathepsin K, COX1 |
| **Glycolysis** | **HK2**, LGR4, **LDHA** | GLUT1, GAPDH |
| **Fibrosis** | FGF7, integrin α5, uPA, α1-antitrypsin, tenascin C | **collagen 3A1**, FGF1, FGF2, CTGF, laminin α5, integrin β1, PAI1, **claudin1**★ |
| **Oncogenesis** | EWSR1, BMI1, AXL, **PDCD4**, ATR, DMBT1, CRIP1, **Ets1**, PIM1 | TWIST1, CRK, NF1, XBP1, Daxx, MALT1, KLF4, BRCA2 |
| **Senescence** | **sirtuin1**, sirtuin6, sirtuin7 | klotho, sirtuin3, p21, caveolin1 |
| **Telomere signaling** |  | TIN2 |

MiR-155-5p\*-CEDS induced the suppression (blue) or activation (red) of signaling through protein downregulation by direct miRNA targeting (blue) or indirect response (sky blue) and protein upregulation by indirect response (red). Some miRNA target proteins were upregulated (★). The data were compared to the reports of recent manuscripts and miRDB, miRmap, miRPathDB, miRTarBase, and TargetScan websites.

The results indicate that miR-155-5p*-CEDS had an anti-oncogenic effect on RAW 264.7 cells by impeding ER stress, angiogenesis, glycolysis, oncogenesis, and acute and chronic inflammation. Conversely, it also had an oncogenic effect on the cells by increasing proliferation, protection, survival, and senescence. It is notable that miR-155-5p*-CEDS can enhance the potential of innate and cellular immunity, as well as growth and cytodifferentiation, which are essential for immune surveillance and recovery.

#### MiR-181a-5p*-CEDS influenced the protein signaling in RAW 264.7 cells

RAW 264.7 cells treated with mmu-miR-181a-5p sequence AACATTCAACGCTGTCGGTGAGT*- CEDS showed the characteristic protein expressions, resulting in the suppression of proliferation, p53/Rb/E2F1 signaling, Notch/Jagged signaling, epigenetic modification, protein translation, miRNA biogenesis, growth factors, cytodifferentiation, osteogenesis, RAS signaling, NFkB signaling, ER stress, survival, innate immunity, cellular immunity, chronic inflammation, p53-mediated apoptosis, FAS-mediated apoptosis, PARP-mediated apoptosis, oncogenesis, and senescence signaling, as well as the activation of cMyc/MAX/MAD network, Wnt/β-catenin signaling, SHH/PTCH/GLI signaling, Notch/Jagged signaling, protection, angiogenesis, acute inflammation, fibrosis, glycolysis, and telomere signaling compared to the untreated controls (Fig. 15A).

**Fig. 15.**
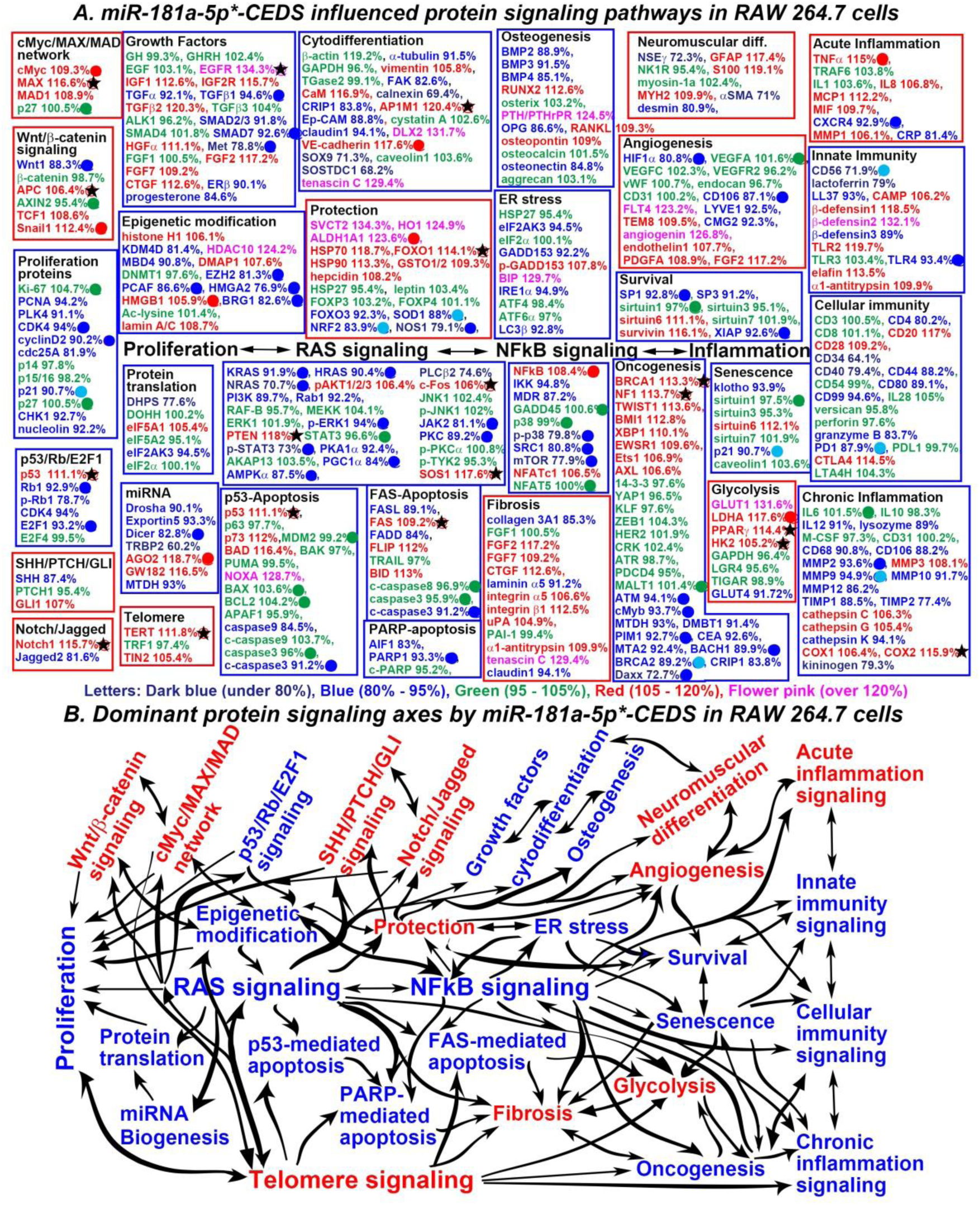
MiR-181a-5p*-CEDS influenced the protein signaling (A) and axes (B) in RAW 264.7 cells. The IP- HPLC revealed proteins downregulated 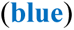, upregulated 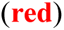, and minimally changed 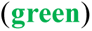 compared to the untreated controls. Dominantly suppressed 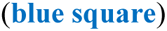 and activated 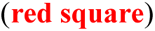 signaling. Downregulated proteins by direct targeting 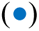 and indirect response 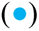, upregulated proteins by indirect response 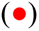, and minimal changed (±5%) proteins 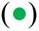. Some target proteins 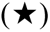 were unexpectedly upregulated contrast to the current manuscripts and data from miRDB and TargetScan websites.

**The proliferation signaling** was suppressed by downregulating miR-181a-5p target proteins CDK4 (94%) and cyclin D2 (90.2%) and miR-181a-5p-responsible protein p21 (90.7%), as well as coincidentally downregulating PCNA (94.2%), PLK4 (91.1%), cdc25A (91.9%), CHK1 (92.7%), and nucleolin (92.2%).

**The p53/Rb/E2F1 signaling** was suppressed by downregulating miR-181a-5p target proteins Rb1 (92.9%) and E2F1 (93.2%) and coincidentally downregulating p-Rb (78.7%) and CDK4 (94%), despite the upregulation of miR-181a target protein p53 (111.1%). P53 is a miR-181a-5p target protein, but it was unexpectedly upregulated compared to the reference data.

**The protein translation signaling** was appears to be impeded by downregulating DHPS (77.6%) and eIF2AK3 (94.5%), despite the slight upregulation of eIF5A1 (105.4%).

**The miRNA biogenesis signaling** appears to be suppressed by downregulating miR-181a-5p target protein Dicer (82.8%) and coincidentally downregulating Drosha (90.1%), Exportin5 (93.3%), Dicer (182.8%), and TRBP2 (60.2%), despite the upregulation of miR-181a-5p-responsible protein AGO2 (118.7%), and coincidental upregulation of GW182 (116.5%) and downregulation of MTDH (93%).

**The growth factor signaling** were partially suppressed by downregulating miR-181a-5p target proteins TGFβ1 (94.6%), SMAD7 (92.6%), and Met (78.8%), as well as coincidentally downregulating TGFα (92.1%), SMAD2/3 (91.8%), ERβ (90.1%), and progesterone (84.6%), despite the upregulation of EFGR (134.3%), IGF1 (112.6%), IGF2R (115.7%), TGFβ2 (120.3%), HGFα (111.1%), FGF2 (117.2%), FGF7 (109.2%), and CTGF (112.6%). EGFR is a miR-181a-5p target protein, but it was unexpectedly upregulated compared to the reference data.

**The cytodifferentiation signaling** were partially inactivated by downregulating α-tubulin (91.5%), FAK (82.6%), calnexin (69.4%), CRIP1 (83.8%), SOX9 (71.3%), and SOSTDC1 (68.2%), despite the upregulation of miR-181a-5p-responsible protein VE-cadherin (117.6%) and coincidental upregulation of vimentin (105.8%), CaM (116.9%), AP1M1 (120.4%), DLX2 (131.7%), and tenascin C (129.4%). AP1M1 is a miR-181a-5p target protein, but it was unexpectedly upregulated compared to the reference data.

**The osteogenesis signaling** was suppressed by downregulating BMP2 (88.9%), BMP3 (91.5%), BMP4 (85.1%), OPG (86.6%), and osteonectin (84.4%), despite the upregulation of RUNX2 (112.6%), PTH/PTHrPR (124.5%), RANKL (109.3%), and osteopontin (109%).

**The RAS signaling** was markedly suppressed by downregulating miR-181a-5p target proteins KRAS (91.9%), NRAS (90.4%), NRAS (70.7%), p-ERK1 (94%), p-STAT3 (73%), PGC1α (84%), AMPKα (87.5%), JAK2 (81.1%), and PKC (89.2%), as well as coincidentally downregulating PI3K (89.7%), Rab1 (92.2%), PKA1α (92.4%), and PLCβ2 (74.6%), despite the upregulation of pAKT1/2/3 (106%), PTEN (118%), c-FOS (106%), and SOS1 (117.6%). PTEN and c-Fos are miR-181a-5p target proteins, but they were unexpectedly upregulated compared to the reference data.

**The NFkB signaling** was suppressed by downregulating miR-181a-5p target proteins p-p38 (79.8%), SRC1 (80.8%), and mTOR (77.9%), as well as coincidentally downregulating IKK (94.8%) and MDR (87.2%), despite the slight upregulation of miR-181a-responsible protein NFkB (108.4%).

**The ER stress signaling** was suppressed by downregulating eIF2AK3 (94.5%), GADD153 (92.2%), IRE1α (94.9%), and LC3β (92.8%), despite the upregulation of p-GADD153 (107.8%) and BIP (129.7%).

**The survival signaling** was suppressed by downregulating SP1 (92.8%) and XIAP (92.6%), and coincidentally downregulating SP3 (91.2%), despite the upregulation of sirtuin6 (111.1%) and sirtuin7 (116.1%).

**The innate immunity signaling** appears to be suppressed by downregulating miR-181a-5p target proteins TLR4 (93.4%) and miR-181a-5p-responsible protein CD56 (71.9%), as well as coincidentally downregulating lactoferrin (79%), LL37 (93%), and β-defensin3 (89%), despite the upregulation of CAMP (106.2%), β- defensin1 (118.5%), β-defensin2 (132.1%), TLR2 (119.7%), elafin (119.7%), and α1-antitrypsin (109.9%).

**The cellular immunity signaling** was suppressed by downregulating miR-181a-5p-responsible proteins PD1 (87.9%), as well as coincidentally downregulating CD4 (80.2%), CD34 (64.1%), CD40 (79.4%), CD80 (89.1%), CD99 (94.6%), and granzyme B (83.7%), despite the upregulation of CD20 (117%), CD28 (109.2%), and CTLA4 (114.5%).

**The chronic inflammation signaling** was suppressed by downregulating miR-181a-5p target proteins MMP2 (93.6%) and miR-181a-5p-responsible protein MMP9 (94.9%), as well as coincidentally downregulating IL12 (91%), lysozyme (89%), CD68 (90.8%), CD106 (88.2%), MMP10 (91.7%), MMP12 (86.2%), TIMP1 (88.5%), TIMP2 (77.4%), cathepsin K (94.1%), and kininogen (79.3%), despite the upregulation of MMP3 (108.1%), cathepsin C (106,3%), cathepsin G (105.4%), COX1 (106.4%), and COX2 (115.9%). COX2 is a miR-181a-5p target protein, but it was unexpectedly upregulated compared to the reference data.

**The p53-mediated apoptosis signaling** appears to be attenuated by downregulating miR-181a-5p target proteins c-caspase3 (91.2%) and coincidentally downregulating caspase9 (84.5%), despite the upregulation of p53 (111.1%), p73 (112%), BAD (116.4%), and NOXA (128.7%). P53 is a miR-181a-5p target protein, but it was unexpectedly upregulated compared to the reference data.

**The FAS-mediated apoptosis signaling** was inhibited by downregulating miR-181a-5p target proteins c- caspase3 (91.2%) and coincidentally downregulating FASL (89.1%) and FADD (84%), and upregulating FLIP (112%), despite the upregulation of FAS (109.2%) and BID (113%). FAS is a miR-181a-5p target protein, but it was unexpectedly upregulated compared to the reference data.

**The PARP-mediated apoptosis signaling** was suppressed by downregulating miR-181a-5p target protein PARP1 (93.3%) and coincidentally downregulating AIF1 (83%).

**The senescence signaling** appears to be suppressed by downregulating miR-181a-5p-responsible protein p21 (90.7%), as well as coincidentally downregulating klotho (93.9%).

Conversely, **the cMyc/MAX/MAD network** was activated by upregulating miR-181a-5p-responsible protein cMyc (109.3%) and coincidentally upregulating MAX (116.6%) and MAD1 (108.9%). MAX is a miR- 181a-5p target protein, but it was unexpectedly upregulated compared to the reference data.

**The Wnt/β-catenin signaling** appears to be activated by upregulating miR-181a-5p-responsible protein Snail1 (112.4%) and coincidental upregulation of TCF1 (108.6%) and APC (106.4%), despite the downregulation of miR-181a target protein Wnt1 (88.3%).

**The SHH/PTCH/GLI signaling** appears to be alternatively activated by upregulating GLI1 (107%), despite the downregulation of SHH (87.4%).

**The Notch/Jagged signaling** appear to be activated by upregulating Notch1 (115.7%), despite the downregulation of Jagged2 (81.6%). Notch1 is a miR-181a-5p target protein, but it was unexpectedly upregulated compared to the reference data.

**The neuromuscular differentiation signaling** were activated by upregulating GFAP (117.4%), S100 (119.1%), and MYH2 (109.9%), despite the downregulation of NSEγ (72.3%), αSMA (71%), and desmin (80.9%).

**The protection signaling** appears to be activated by upregulating miR-181a-5p-responsible proteins ALDH1A1 (123.6%) and coincidentally upregulating SVCT2 (134.3%), HO1 (124.9%), HSP70 (116.7%), FOXO1 (114.1%), HSP90 (113.3%), GSTO1/2 (109.3%), and hepcidin (108.2%), despite the downregulation of miR-181a-5p target protein NOS1 (79.1%) and miR-181a-5p-responsible protein SOD1 (88%) and NRF2 (83.9%), and the coincidental downregulation of FOXO3 (92.3%). FOXO1 is a miR-181a-5p target protein, but it was unexpectedly upregulated compared to the reference data.

**The angiogenesis signaling** was activated by upregulating FLT4 (123.2%), TEM8 (109.5%), angiogenin (126.8%), endothelin1 (107.7%), PDGFA (108.9%), and FGF2 (117.2%), despite the downregulation of miR- 181a-5p target proteins HIF1α (80.8%) and CD106 (87.1%), as well as coincidental downregulation of LYVE1 (92.5%) and CMG2 (92.3%).

**The acute inflammation signaling** was activated by upregulating miR-181a-5p-responsible protein TNFα (115%) and coincidentally upregulating IL8 (106.8%), MCP1 (112.2%), MIF (109.7%), and MMP1 (106.1%), despite the downregulation of miR-181a-5p target protein CXCR4 (92.9%) and the coincidental downregulation of CRP (81.4%).

**The fibrosis signaling** was activated by upregulating FGF2 (177.2%), CTGF (112.6%), integrin α5 (106.6%), integrin β1 (112.5%), uPA (104.9%), α1-antitrypsin (109.9%), and tenascin C (129.4%), despite the downregulation of collagen 3A1 (85.3%), laminin α5 (91.2%), and claudin1 (94.1%) and the upregulation of FGF7 (109.2%).

**The glycolysis signaling** was activated by upregulating miR-181a-5p-responsible protein LDHA (117.6%), as well as coincidentally upregulating GLUT1 (131.6%), PPARγ (114.4%), and HK2 (105.2%), despite the downregulation of GLUT4 (91.7%). PPARγ and HK2 are miR-181a-5p target proteins, but they are unexpectedly upregulated compared to the reference data.

**The telomere signaling** was activated by upregulating TERT (111.8%) and TIN2 (105.4%).

In addition, miR-181a-5p*-CEDS markedly influenced **the epigenetic modification signaling** by downregulating miR-181a-5p target proteins EZH2 (81.3%), PCAF (86.6%), HMGA2 (76.9%), and BRG1 (80.9%), as well as the coincidental downregulation of KDM4D (81.4%) and MBD4 (90.8%), while it upregulated miR-181a-5p-responsible protein HMGB1 (105.9%) and coincidental upregulation of histone H1 (106.1%), HDAC10 (124.2%), DMAP1 (107.6%), and lamin A/C 108.7%). Consequently, miR-181a-5p*- CEDS induced a trend towards a reduction in the methylation of histones and DNAs, and attenuation of transcriptional repression.

MiR-181a-5p*-CEDS significantly suppressed **the oncogenesis signaling** by downregulating miR-181a-5p target proteins ATM (94.1%), cMyb (93.7%), PIM1 (92.7%), BACH1 (89.9%), and Daxx (72.7%) and miR- 181a-5p-responsible protein BRCA2 (89.2%), while it also activated **the oncogenesis signaling** by upregulating BRCA1 (113.3%), NF1 (113.7%), TWIST1 (113.6%), BMI1 (112.8%), XBP1 (110.1%), EWSR1 (109.6%), Ets1 (106.9%), and AXL (106.6%). BRCA1 and NF1 are miR-181a-5p target proteins, but they were unexpectedly upregulated compared to the reference data. Among the 28 oncoproteins, 11 were found to be underexpressed, eight were overexpressed, and nine exhibited minimal change in expression compared to the untreated controls.

Consequently, miR-181a-5p*-CEDS exerted a profound influence on the entire protein signaling in RAW 264.7 cells. MiR-181a-5p*-CEDS suppressed the NFkB signaling, inactivating the epigenetic modification- cMyc/MAX/MAD network-Wnt/β-catenin signaling axis, ER stress-angiogenesis-acute inflammation, and glycolysis-oncogenesis-chronic inflammation signaling axis. Conversely, it enhanced the RAS signaling, and subsequently activated proliferation signaling axis, protection-survival-senescence signaling axis, and FAS-mediated apoptosis-fibrosis signaling axis (Fig. 15B, Table 8).

**Table 8.** Protein expressions in the RAW 264.7 cells treated with miR-181a-5p*-CEDS.

| Signaling | Downregulated Proteins | Upregulated Proteins |
| --- | --- | --- |
| Proliferation | PCNA, PLK4, CDK4, **cyclin D2**, cdc25A, **p21**, CHK1, nucleolin |  |
| cMyc/MAX/MAD network |  | **cMyc**, **MAX**★, MAD1 |
| Wnt/β-catenin signaling | Wnt1 | **APC**★, **Snail** |
| p53/Rb/E2F1 signaling | **Rb1**, p-Rb1, CDK4, **E2F1** | **p53**★ |
| SHH/PTCH/GLI signaling | SHH | GLI1 |
| Notch/Jagged signaling | Jagged2 | **Notch1**★ |
| Epigenetic modification | KDM4D, MBD4, **EZH2**, **PCAF**, **HMG2**, **BRG1** | histone H1, HDAC10, DMAP1, **HMG1**, lamin A/C |
| Protein translation | DHPS, eIF2AK3 | eIF5A1 |
| miRNA biogenesis | Drosha, Exportin5, **Dicer**, TRBP2, MTDH | **AGO2**, GW182 |
| Growth factors | TGFα, **TGFβ1**, SMAD2/3, **SMAD7**, **Met**, ERβ, progesterone | **EGFR**★, IGF1, IGF2R, TGFβ2, HGFα, FGF2, FGF7, CTGF |
| Cytodifferentiation | α-tubulin, FAK, calnexin, CRIP1, , Ep-CAM, claudin1, SOX9, SOSTDC1 | vimentin, CaM, **AP1M1**★, DLX2, **VE-cadherin**, tenascin C |
| Osteogenesis | BMP2, BMP3, BMP4, OPG, osteonectin | RUNX2, PTH/PTHrPR, RANKL, osteopontin |
| Neuromuscular differentiation | NSEγ, αSMA, desmin, | GFAP, S100, MYH2 |
| RAS signaling | **KRAS**, **HRAS**, **NRAS**, PI3K, Rab1, RAF-B, **p-ERK1**, p-STAT3, PKA1α, **PGC1α**, **AMPKα**, PLCβ2, **JAK2**, **PKC**, p-TYK2 | pAKT1/2/3, **PTEN**★, c-Fos★, **SOS1**★ |
| NFκB signaling | IKK, MDR, **p-p38**, **SRC1**, **mTOR** | **NFκB**, NFATc1 |
| p53-mediated apoptosis | caspase9, c-caspase9 | **p53**★, p73, BAD, NOXA |
| FAS-mediated apoptosis | FASL, FADD, **c-caspase3** | **FAS**★, FLIP, BID |
| PARP-mediated apoptosis | AIF1, **PARP1** |  |
| Protection | FOXO3, **SOD1**, **NRF2**, **NOS1** | SVCT2, HO1, **ALDH1A1**, HSP70, **FOXO1**★, HSP90, GSTO1/2, hepcidin |
| ER stress | eIF2AK3, GADD153, IRE1α, LC3β | p-GADD153, BIP |
| Angiogenesis | **HIF1α**, **CD106**, LYVE1, CMG2 | FLT4, TEM8, angiogenin, endothelin1, PDGFA, FGF2 |
| Survival | **SP1**, SP3, **XIAP** | sirtuin6, survivin, |
| Acute inflammation | **CXCR4**, CRP | **TNFα**, IL8, MCP1, MIF, MMP1 |
| Innate immunity | **CD56**, lactoferrin, LL37, β-defensin3, **TLR4** | CAMP, β-defensin1, β-defensin2, TLR2, elafin, α1-antitrypsin |
| Cellular immunity | CD4, CD34, CD40, CD44, CD80, CD99, granzyme B, **PD1** | CD20, CD28, CTLA4 |
| Chronic inflammation | IL12, lysozyme, CD68, CD106, **MMP2**, **MMP9**, MMP10, MMP12, TIMP1, TIMP2, cathepsin K, kininogen | MMP3, cathepsin C, cathepsin G, COX1, **COX2**★ |
| Glycolysis | GLUT4 | GLUT1, **LDHA**, **PPARγ**★, **HK2**★ |
| Fibrosis | collagen 3A1, laminin α5, claudin1 | FGF2, FGF7, CTGF, integrin α5, integrin β1, uPA, α1-antitrypsin, tenascin C |
| Oncogenesis | **ATM**, cMyb, MTDH, DMBT1, **PIM1**, CEA, MTA2, **BACH1**, **BRCA2**, CRIP1, **Daxx** | **BRCA1**★, **NF1**★, TWIST1, BMI1, XBP1, EWSR1, Ets1, AXL, ATM |
| Senescence | klotho, **p21**, | sirtuin6 |
| Telomere signaling |  | **TERT**★, TIN2 |

MiR-181a-5p\*-CEDS induced the suppression (blue) or activation (red) of signaling through protein downregulation by direct miRNA targeting (blue) or indirect response (sky blue) and protein upregulation by indirect response (red). Some miRNA target proteins were upregulated (★). The data were compared to the reports of recent manuscripts and miRDB, miRmap, miRPathDB, miRTarBase, and TargetScan websites.

The results indicate that miR-181a-5p*-CEDS had an anti-oncogenic effect on RAW 264.7 cells by impeding ER stress, angiogenesis, glycolysis, oncogenesis, and acute and chronic inflammation. Conversely, it also had an oncogenic effect on the cells by increasing proliferation, protection, survival, and senescence. It is notable that miR-181a-5p*-CEDS can enhance the potential of innate and cellular immunity, as well as growth and cytodifferentiation, which are essential for immune surveillance and recovery.

#### MiR-216a-5p*-CEDS influenced the protein signaling in RAW 264.7 cells

RAW 264.7 cells treated with mmu-miR-216a-5p sequence TAATCTCAGCTGGCAACTGTGA*-CEDS showed the characteristic protein expressions, resulting in the suppression of proliferation, Wnt/β-catenin signaling, p53/Rb/E2F1 signaling, SHH/PTCH/GLI signaling, Notch/Jagged signaling, epigenetic modification, growth factors, cytodifferentiation, neuromuscular differentiation, RAS signaling, NFkB signaling, angiogenesis, acute inflammation, innate immunity, cellular immunity, chronic inflammation, FAS-mediated apoptosis, PARP-mediated apoptosis, fibrosis, oncogenesis, and senescence, as well as the activation of cMyc/MAX/MAD network, miRNA biogenesis, osteogenesis, protection, ER stress, survival, p53-mediated apoptosis, glycolysis, and telomere signaling compared to the untreated controls (Fig. 16A).

**Fig. 16.**
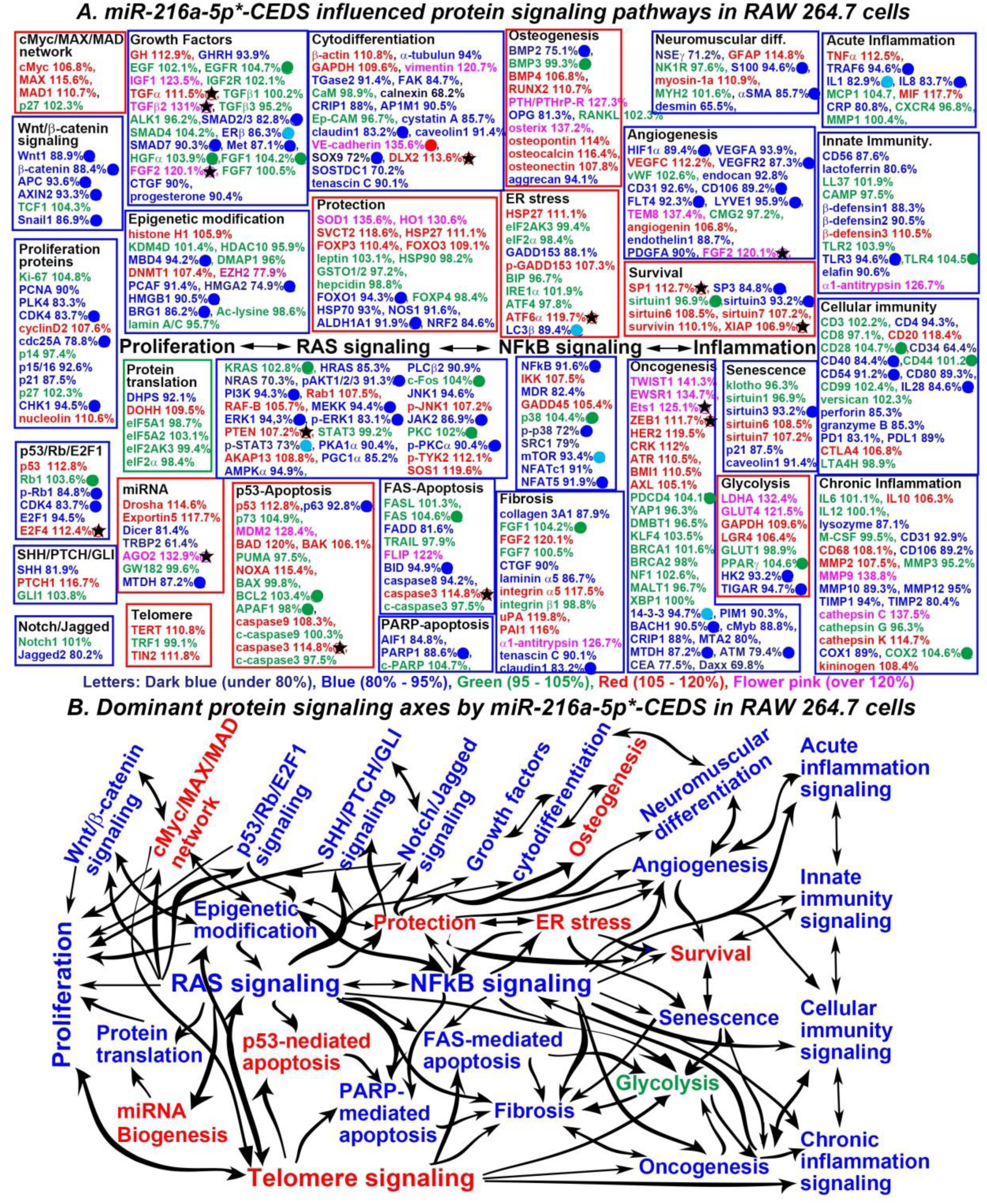
MiR-216a-5p*-CEDS influenced the protein signaling (A) and axes (B) in RAW 264.7 cells. The IP- HPLC revealed proteins downregulated 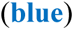, upregulated 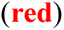, and minimally changed 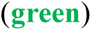 compared to the untreated controls. Dominantly suppressed 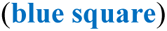 and activated 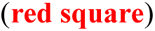 signaling. Downregulated proteins by direct targeting 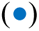 and indirect response 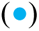, upregulated proteins by indirect response 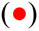, and minimal changed (±5%) proteins 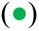. Some target proteins 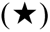 were unexpectedly upregulated contrast to the current manuscripts and data from miRDB and TargetScan websites.

**The proliferation signaling** was markedly suppressed by downregulating miR-216a-5p target proteins CDK4 (83.7%), cdc25A (78.8%), and CHK1 (94.5%), as well as coincidentally downregulating PCNA (90%), and PLK4 (83.3%), despite the upregulation of cyclin D2 (107.6%) and nucleolin (110.6%) and the downregulation of p15/16 (92.6%) and p21 (87.5%).

**The Wnt/β-catenin signaling** was markedly suppressed by downregulating miR-216a-5p target proteins Wnt1 (88.9%), β-catenin (88.4%), APC (93.6%), and Snail1 (86.9%).

**The p53/Rb/E2F1 signaling** was suppressed by downregulating miR-216a-5p target proteins p-Rb (84.8%) and CDK4 (83.7%), as well as the downregulation of E2F1 (94.5%) and upregulation of E2F4 (112.4%), despite the upregulation of p53 (112.8%).

**The SHH/PTCH/GLI signaling** appears to be inactivated by downregulating SHH (81.9%), despite the upregulation of PTCH1 (116.7%).

**The Notch/Jagged signaling** was inactivated by downregulating Jagged2 (80.2%).

**The growth factor signaling** were suppressed by downregulating miR-216a-5p target proteins SMAD2/3 (82.8%), SMAD7 (90.3%), and Met (87.1%) and miR-216a-5p-responsible protein ERβ (86.3%), as well as coincidentally downregulating GHRH (93.99%), CTGF (90%), and progesterone (90.4%), despite the upregulation of GH (112.9%), IGF1 (123.5%), TGFα (111.5%), TGFβ2 (131%), and FGF2 (120.1%). TGFα (111.5%), TGFβ2 (131%), and FGF2 are miR-216a-5p target proteins, but they were unexpectedly upregulated compared to the reference data.

**The cytodifferentiation signaling** were suppressed by downregulating miR-216a-5p target proteins claudin1 (83.2%) and SOX9 (72%) and coincidentally downregulating α-tubulin (94%), TGase2 (91.4%), FAK (84.7%), calnexin (68.2%), CRIP1 (88%), AP1M1 (90.5%), cystatin A (85.7%), claudin1 (83.2%), caveolin1 (91.4%), SOX9 (72%), SOSTDC1 (70.2%), and tenascin C (90.1%), despite the upregulation of β-actin (110.8%), GAPDH (109.6%), vimentin (120.7%), VE-cadherin (135.6%), and DLX2 (113.6%). DLX2 is a miR-216a-5p target protein, but it was unexpectedly upregulated compared to the reference data.

**The neuromuscular signaling** was suppressed by downregulating miR-216a-5p target proteins S100 (94.6%) and αSMA (85.7%) and coincidentally downregulating desmin (65.5%), despite the upregulation of GFAP (114.8%) and myosin-1a (110.9%).

**The RAS signaling** was suppressed by downregulating miR-216a-5p target proteins pAKT1/2/3 (91.3%), PI3K (94.3%), MEKK (94.4%), ERK1 (94.3%), p-ERK1 (83.1%), JAK2 (86.9%), and p-PKCα (90.4%) and miR-216a-5p-responsible protein p-STAT3 (73%), as well as coincidentally downregulating HRAS (85.3%), NRAS (70.3%), PKA1α (90.4%), PGC1α (85.2%), AMPKα (94.9%), PLCβ2 (90.9%), and JNK1 (94.6%), despite the upregulation of Rab1 (107.5%), RAF-B (105.7%), PTEN (107.2%), AKAP13 (108.8%), p-JNK1 (107.2%), p-TYK2 (112.1%), and SOS1 (119.6%). PTEN is a miR-216a-5p target protein, but it was unexpectedly upregulated compared to the reference data.

**The NFkB signaling** was suppressed by downregulating miR-216a-5p target proteins NFkB (91.6%), p- p38 (72%), and NFAT5 (91.9%) and miR-216a-5p-responsible protein mTOR (93.4%), as well as coincidentally downregulating MDR (82.4%), SRC1 (79%), and NFATc1 (91%), despite the upregulation of IKK (107.5%) and GADD45 (105.4%).

**The angiogenesis signaling** was suppressed by downregulating miR-216a-5p target proteins HIF1α (89.4%), VEGFR2 (87.3%), CD106 (89.2%), FLT4 (92.3%), and LYVE1 (95.9%), as well as coincidentally downregulating VEGFA (93.9%), endocan (92.8%), CD31 (92.6%), endothelin1 (88.7%), and PDGFA (90%), despite the upregulation of VEGFC (112.2%), TEM8 (137.4%), angiogenin (106.8%), and FGF2 (120.1%).

**The acute inflammation signaling** was suppressed by downregulating miR-216a-5p target proteins TRAF6 (94.6%), IL8 (83.7%), and miR-216a-5p-responsible protein IL1 (82.9%), despite the upregulation of TNFα (112.5%) and MIF (117.7%).

**The innate immunity signaling** was suppressed by downregulating miR-216a-5p target protein TLR3 (94.6%), as well as coincidentally downregulating CD56 (87.6%), lactoferrin (80.6%), β-defensin1 (88.3%), β- defensin2 (90.5%), and elafin (90.6%), despite the upregulation of β-defensin3 (110.5%) and α1-antitrypsin (126.7%).

**The cellular immunity signaling** was suppressed by downregulating miR-216a-5p-responsible proteins CD40 (84.4%), CD54 (91.2%), and IL2 (84.6%), as well as coincidentally downregulating CD4 (94.3%), CD34 (64.4%), CD80 (89.3%), perforin (85.3%), and granzyme B (85.3%), despite the downregulation of PD1 (83.1%) and PDL1 (89%), and coincidental upregulation of CD20 (118.4%) and CTLA4 (106.8%).

**The chronic inflammation signaling** appears to be inactivated by downregulating lysozyme (87.1%), CD31 (92.9%), CD106 (89.2%), MMP10 (89.3%), MMP12 (95%), TIMP1 (94%), TIMP2 (80.4%), and COX1 (89%), despite the upregulation of IL10 (106.3%), CD68 (108.1%), MMP2 (107.5%), MMP9 (138.8%), cathepsin C (137.5%), cathepsin K (114.7%), and kininogen (108.4%).

**The FAS-mediated apoptosis signaling** appears to be inhibited by downregulating miR-216a-5p target protein BID (94.9%) and coincidentally downregulating FADD (81.6%) and upregulating FLIP (122%), despite the upregulation of caspase3 (114.8%). Caspase3 is a miR-216a-5p target protein, but it was unexpectedly upregulated compared to the reference data.

**The PARP-mediated apoptosis signaling** was suppressed by downregulating miR-216a-5p target protein PARP1 (88.6%) and coincidentally downregulating AIF1 (84.8%).

**The fibrosis signaling** was inactivated by downregulating miR-216a-5p target protein claudin (83.2%), as well as coincidentally downregulating collagen 3A1 (87.9%), CTGF (90%), laminin α5 (86.7%), and tenascin C (90.1%), despite the upregulation of FGF2 (120.1%), integrin α5 (86.7%), uPA (119.8%), PAI1 (116%), and α1-antitrypsin (126.7%).

**The senescence signaling** was inactivated by downregulating miR-216a-5p target protein sirtuin3 (93.2%), as well as coincidentally downregulating p21 (87.5%), and caveolin1 (91.4%), despite the slight upregulation of sirtuin6 (108.5%) and sirtuin7 (107.2%).

Conversely, **the cMyc/MAX/MAD network** was activated by upregulating cMyc(106.8%), MAX (115.6%), and MAD1 (110.7%).

**The miRNA biogenesis signaling** was activated by downregulating miR-216a-5p target proteins MTDH (87.2%) and coincidental upregulating Drosha (114.6%), Exportin5 (117.7%), and AGO2 (132.9%), despite the downregulation of Dicer (81.4%) and TRBP2 (61.4%).

**The osteogenesis signaling** was activated by upregulating BMP4 (106.8%), RUNX2 (110.7%), PTH/PTHrP-R (127.3%), osterix (137.2%), osteopontin (114%), osteocalcin (116.4%), and osteonectin (107.8%), despite the downregulation of miR-216a-5p target protein BMP2 (75.1%), and coincidental downregulation of OPG (81.3%) and aggrecan (94.1%).

**The protection signaling** was activated by upregulating SOD1 (135.6%), HO1 (130.6%), SVCT2 (118.6%), HSP27 (111.1%), FOXP3 (110.4%), and FOXO3 (109.1%), despite the downregulation of miR-216a- 5p target protein FOXO1 (94.3%), and ALDH1A1 (91.9%) and coincidental downregulation of HSP70 (93%), NOS1 (91.6%), and NRF2 (84.6%).

**The ER stress signaling** appears to be activated by upregulating HSP27 (111.1%), p-GADD163 (107.3%), and ATF6α (119.7%), despite the downregulation of miR-216a-5p-responsible proteins LC3β (89.4%) and coincidental downregulation of GADD153 (88.1%). ATF6α is a miR-216a-5p target protein, but it was unexpectedly upregulated compared to the reference data.

**The survival signaling** was activated by upregulating SP1 (112.7%), sirtuin6 (108.5%), sirtuin7 (107.2%), survivin (110.1%) and XIAP (106.9%), despite the downregulation of miR-216a-5p target proteins SP3 (84.4%) and sirtuin3 (93.2%). SP1 and XIAP are miR-216a-5p target proteins, but they were unexpectedly upregulated compared to the reference data.

**The p53-mediated apoptosis signaling** appears to be activated by upregulating p53 (112.8%), MDM2 (128.4%), BAD (120%), BAK (106.1%), NOXA (115.4%), caspase9 (108.3%), and caspase3 (114.8%), despite the downregulation of miR-216a-5p target proteins p63 (92.8%). Caspase3 is a miR-216a-5p target protein, but it was unexpectedly upregulated compared to the reference data.

**The glycolysis signaling** was activated by upregulating LDHA (132.4%), GLUT4 (121.5%), GAPDH (109.6%), and LGR4 (106.4%), despite the slight downregulation of miR-216a-5p target protein HK2 (93.2%) and TIGAR (94.7%).

**The telomere signaling** was activated by upregulating TERT (110.8%) and TIN2 (111.8%).

In addition, miR-216a-5p*-CEDS markedly influenced **the epigenetic modification signaling** by downregulating miR-216a-5p target proteins MBD4 (94.2%), HMGA2 (74.9%), HMGB1 (90.5%), and BRG1 (86.2%), as well as the coincidental downregulation of PCAF (91.4%), while it upregulated histone H1 (105.9%), DNMT1 (107.4%), and EZH2 (77.9%). Consequently, miR-216a-5p*-CEDS induced a trend towards a reduction in the methylation of histones and DNAs, and attenuation of transcriptional repression.

MiR-216a-5p*-CEDS suppressed **the oncogenesis signaling** by downregulating miR-216a-5p target proteins BACH1 (90.5%), MTDH 87.2%), and ATM (79.4%) and miR-216a-5p-responsible protein 14-3-3 (94.7%), as well as coincidentally downregulating PIM1 (90.3%), cMyb (88.8%), CRIP1 (88%), MTA2 (80%), CEA (77.5%), and Daxx (69.8%), while it partly enhanced **the oncogenesis signaling** by upregulating TWIST1 (141.3%), EWSR1 (134.7%), Ets1 (125.1%), ZEB1 (111,7%), HER2 (119.5%), CRK (112%), ATR (110.5%), BMI1 (110.5%), and AXL (105.1%). Ets1 and ZEB1 are miR-216a-5p target proteins, but they were unexpectedly upregulated compared to the reference data. Among the 28 oncoproteins, 10 were found to be underexpressed, nine were overexpressed, and nine exhibited minimal change in expression compared to the untreated controls.

Consequently, miR-216a-5p*-CEDS significantly impacted the entire protein signaling in RAW 264.7 cells. MiR-216a-5p*-CEDS suppressed the RAS-NFkB signaling axis, and subsequently inactivated the proliferation-related signaling axis, angiogenesis-inflammation axis, and senescence-oncogenesis axis. Conversely, it activated protection-ER stress-survival signaling axis, protection-cMyc-telomere, and p53- mediated apoptosis signaling axis (Fig. 16B, Table 9).

**Table 9.** Protein expressions in the RAW 264.7 cells treated with miR-216a-5p*-CEDS.

| Signaling | Downregulated Proteins | Upregulated Proteins |
| --- | --- | --- |
| Proliferation | PCNA, PLK4, **CDK4**, **cdc25A**, p15/16, p21, **CHK1** | Cyclin D2, nucleolin |
| **cMyc/MAX/MAD network** |  | cMyc, MAX, MAD1 |
| Wnt/β-catenin signaling | **Wnt1**, β-catenin, APC, AXIN2, Snail | |
| p53/Rb/E2F1 signaling | **p-Rb1**, CDK4, E2F1 | p53, E2F4 |
| SHH/PTCH/GLI signaling | SHH | PTCH1 |
| Notch/Jagged signaling | Jagged2 |  |
| Epigenetic modification | **MBD4**, PCAF, **HMGA2**, **HMGB1**, **BRG1**, Ac-lysine | histone H1, DNMT1, EZH2 |
| Protein translation | DHPS | DOHH |
| **miRNA biogenesis** | Dicer, TRBP2, **MTDH** | Drosha, Exportin5, **AGO2**★ |
| Growth factors | GHRH, **SMAD2/3**, **SMAD7**, Met, CTGF, **ERβ**, progesterone | GH, EGFR, IGF1, **TGFα★**, **TGFβ2★**, HGFα, FGF1, FGF2 |
| Cytodifferentiation | α-tubulin, TGase2, FAK, calnexin, CRIP1, AP1M1, cystatin A, **claudin1**, caveolin1, **SOX9**, SOSTDC1, tenascin C | β-actin, GAPDH, vimentin, **VE-cadherin**, **DLX2**★ |
| **Osteogenesis** | **BMP2**, OPG, aggrecan | BMP4, RUNX2, osterix, PTH/PTHrPR, osteopontin, osteocalcin, osteonectin |
| Neuromuscular differentiation | NSEγ, **S100**, **αSMA**, desmin | GFAP, myosin-1a |
| RAS signaling | HRAS, NRAS, **pAKT1/2/3**, **PI3K**, **MEKK**, **ERK1**, p-ERK1, **p-STAT3**, PKA1α, PGC1α, AMPKα, PLCβ2, JNK1, **JAK2**, **p-PKC1α** | Rab1, RAF-B, **PTEN**★, AKAP13, c-Fos, p-JNK1, p-TYK2, SOS1 |
| NFκB signaling | **NFκB**, MDR, p-p38, SRC1, **mTOR**, NFATc1, **NFAT5** | IKK, GADD45 |
| **p53-mediated apoptosis** | **p63** | p53, MDM2, BAD, BAK, NOXA, BCL2, caspase9, **caspase3**★ |
| FAS-mediated apoptosis | FADD, **BID**, caspase8 | FLIP, **caspase3**★ |
| PARP-mediated apoptosis | AIF1, **PARP1** |  |
| **Protection** | **FOXO1**, HSP70, NOS1, **ALDH1A1**, NRF2 | SOD1, HO1, SVCT2, HSP27, FOXp3, FOXO3 |
| **ER stress** | GADD153, **LC3β** | HSP27, p-GADD153, **ATF6α★** |
| Angiogenesis | **HIF1α**, VEGFA, **VEGFR2**, endocan, CD31, **CD106**, **FLT4**, **LYVE1**, endothelin1, PDGFA | VEGFC, TEM8, angiogenin, **FGF2**★ |
| **Survival** | **SP3**, sirtuin3 | **SP1**★, sirtuin6, sirtuin7, survivin, XIAP |
| Acute inflammation | **ITRAF6**, **L1**, **IL8**, CRP | TNFα, MIF |
| Innate immunity | CD56, lactoferrin, β-defensin1, β-defensin1, β-defensin2, **TLR3**, elafin | β-defensin3, α1-antitrypsin |
| Cellular immunity | CD4, CD34, **CD40**, **CD54**, CD80, **IL28**, perforin, granzyme B, PD1, PDL1 | CD20, CD28, CTLA4 |
| Chronic inflammation | Lysozyme, CD31, CD106, MMP10, MMP12, TIMP1, TIMP2, COX1 | IL10, CD68, MMP2, MMP9, cathepsin C, cathepsin K, COX2, kininogen |
| **Glycolysis** | **HK2**, **TIGAR** | LDHA, GAPDH, LGR4 |
| Fibrosis | collagen 3A1, CTGF, laminin α5, tenascin C, **claudin1** | FGF2, integrin α5, uPA, PAI1, α1-antitrypsin |
| Oncogenesis | **14-3-3**, PIM1, **BACH1**, cMyb, CRIP1, MTA2, **MTDH**, **ATM**, CEA, Daxx | TWIST1, EWSR1, **Ets1**★, **ZEB1**★, HER2, CRK, ATR, BMI, PDCD4, AXL |
| Senescence | **sirtuin3**, p21, caveolin1 | sirtuin6, sirtuin7 |
| **Telomere signaling** |  | TERT, TIN2 |

MiR-216a-5p\*-CEDS induced the suppression (blue) or activation (red) of signaling through protein downregulation by direct miRNA targeting (blue) or indirect response (sky blue) and protein upregulation by indirect response (red). Some miRNA target proteins were upregulated (★). The data were compared to the reports of recent manuscripts and miRDB, miRmap, miRPathDB, miRTarBase, and TargetScan websites.

The results indicate that miR-216a-5p*-CEDS had a dual effect on RAW 264.7 cells. It had an anti-oncogenic effect, impeding proliferation, angiogenesis, senescence, inflammation, and oncogenesis. On the other hand, it also exerted an oncogenic effect, enhancing ROS protection, telomere instability, and p53- mediated apoptosis. Therefore, it is postulated that miR-216a-5p*-CEDS may play an oncogenic role through the increase of ROS protection signaling, depending on the context of the cells. However, it is notable that miR- 216a-5p*-CEDS could attenuate the capacity of innate and cellular immunity, as well as chronic inflammation.

#### MiR-365a-3p*-CEDS influenced the protein signaling in RAW 264.7 cells

RAW 264.7 cells treated with mmu-miR-365a-3p sequence TAATGCCCCTAAAAATCCTTAT*-CEDS showed the characteristic protein expressions, resulting in the suppression of proliferation, cMyc/MAX/MAD network, Wnt/β-catenin signaling, p53/Rb/E2F1 signaling, SHH/PTCH/GLI signaling, Notch/Jagged signaling, epigenetic modification, protein translation, growth factors, cytodifferentiation, RAS signaling, NFkB signaling, protection, ER stress, innate immunity, cellular immunity, chronic inflammation, FAS-mediated apoptosis, and glycolysis signaling, as well as the activation of miRNA biogenesis, osteogenesis, neuromuscular differentiation, angiogenesis, survival, acute inflammation, p53-mediated apoptosis, PARP-mediated apoptosis, fibrosis, oncogenesis, and senescence compared to the untreated controls (Fig. 17A).

**Fig. 17.**
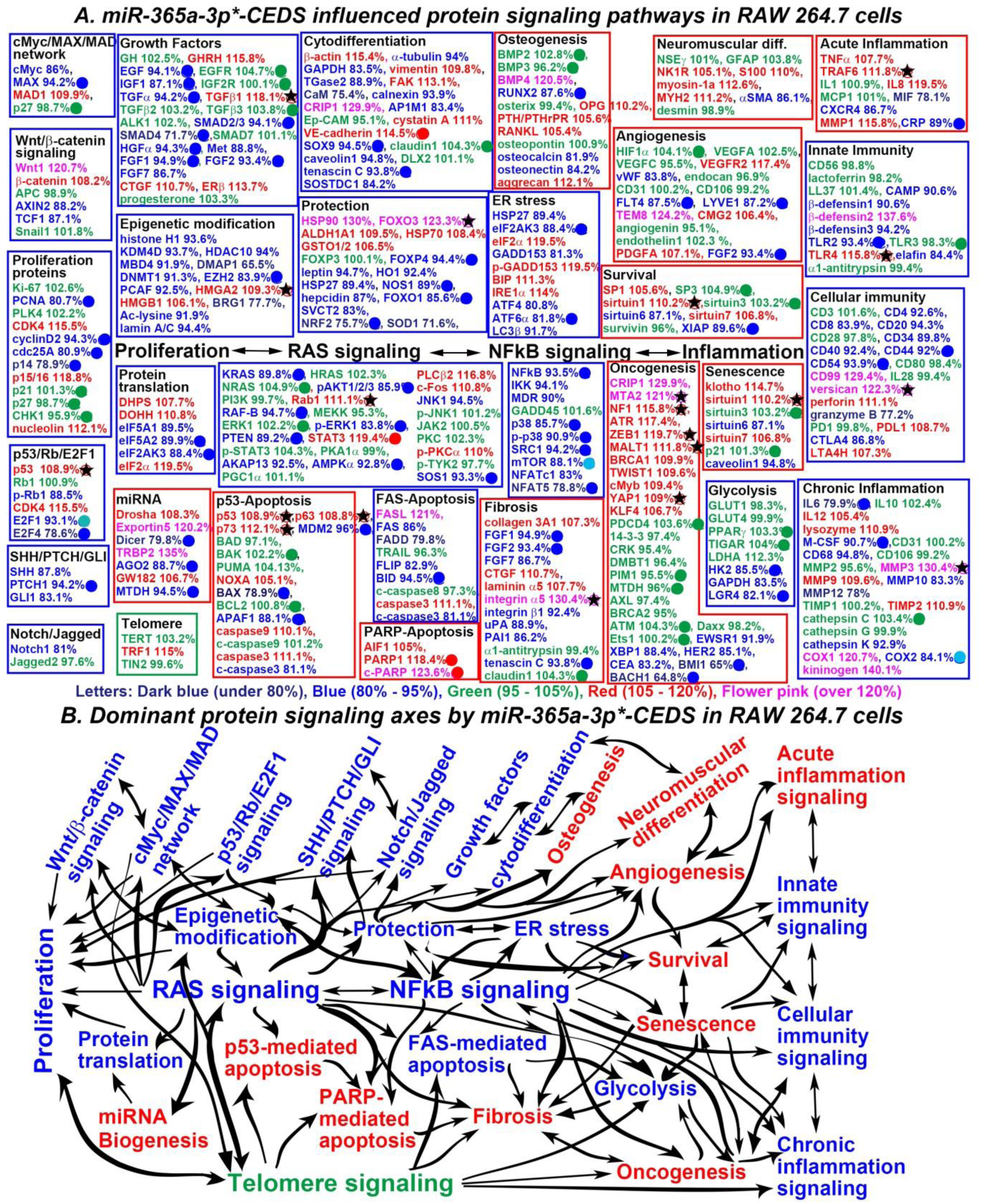
MiR-365a-3p*-CEDS influenced the protein signaling (A) and axes (B) in RAW 264.7 cells. The IP- HPLC revealed proteins downregulated 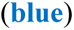, upregulated 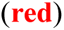, and minimally changed 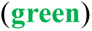 compared to the untreated controls. Dominantly suppressed 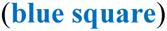 and activated 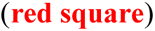 signaling. Downregulated proteins by direct targeting 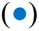 and indirect response 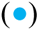, upregulated proteins by indirect response 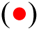, and minimal changed (±5%) proteins 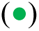. Some target proteins 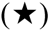 were unexpectedly upregulated contrast to the current manuscripts and data from miRDB and TargetScan websites.

**The proliferation signaling** appears to be suppressed by downregulating miR-365a-3p target proteins PCNA (80.7%), cyclin D2 (94.3%), and cdc25A (80.9%), as well as coincidentally upregulating p15/16 (118.8%), despite the upregulation of CDK4 (115.5%) and nucleolin (109.8%), and the downregulation of miR- 365a-3p target protein p14 (78.9%).

**The cMyc/MAX/MAD network** was suppressed by downregulating miR-365a-3p target proteins MAX (94.2%), and coincidentally downregulating cMyc (86%) and upregulating MAD1 (109.9%).

**The Wnt/β-catenin signaling** appears to be inactivated by downregulating AXIN2 (88.2%) and TCF1 (87.1%), despite the upregulation of Wnt1 (120.7%) and β-catenin (108.2%).

**The p53/Rb/E2F1 signaling** appeared to be inactivated by downregulating p-Rb1 (88.5%) and coincidentally downregulating miR-365a-3p-reponsible protein E2F1 (93.1%), despite the downregulation of miR-365a-3p target protein E2F4 (78.6%), and coincidental upregulation of p53 (108.9%) and CDK4 (115.5%). P53 is a miR-365a-3p target protein, but it was unexpectedly upregulated compared to the reference data.

**The SHH/PTCH/GLI signaling** was suppressed by downregulating miR-365a-3p target proteins PTCH1 (94.2%), and coincidentally downregulating SHH (87.8%) and GLI1 (83.1%).

**The Notch/Jagged signaling** appears to be inactivated by downregulating Notch1 (81%).

**The protein translation signaling** was inactivated by downregulating miR-365a-3p-responsible proteins eIF5A2 (89.9%) and eIF2AK3 (88.4%), and coincidentally downregulating eIF5A1 (89.5%), despite the upregulation of DHPS (107.7%), DOHH (110.8%), and eIF2α (119.5%).

**The growth factor signaling** appear to be suppressed by downregulating miR-365a-3p target proteins EGF (94.1%), IGF1 (87.1%), TGFα (94.2%), SMAD2/3 (94.1%), SMAD4 (71.7%), HGFα (94.3%), FGF1 (94.9%), FGF2 (93.4%), and FGF7 (86.7%), as well as coincidentally downregulating Met (88.8%), despite the upregulation of GHRH (115.8%), TGFβ1 (118.1%), CTGF (110.7%), and ERβ (113.7%). TGFβ1 is a miR- 365a-3p target protein, but it was unexpectedly upregulated compared to the reference data.

**The cytodifferentiation signaling** appear to be suppressed by downregulating miR-365a-3p target protein SOX9 (94.5%) and tenascin C (93.8%), as well as coincidentally downregulating α-tubulin (94%), GAPDH (83.5%), TGase2 (88.9%), CaM (75.4%), calnexin (93.9%), AP1M1 (83.4%), caveolin1 (94.8%), and SOSTDC1 (84.2%), despite the upregulation of miR-365a-3p-responsible protein VE-cadherin (114.5%) and coincidental upregulation of β-actin (115.4%), vimentin (109.8%), FAK (113.1%), CRIP1 (129.9%), and cystatin A (111%).

**The RAS signaling** was suppressed by downregulating miR-365a-3p target proteins KRAS (89.8%), pAKT1/2/3 (85.9%), RAF-B (94.7%), p-ERK1 (83.8%), PTEN (89.2%), AMPKα (92.8%), and SOS1 (93.3%), as well as coincidentally downregulating AKAP13 (92.5%) and JNK (94.5%), despite the upregulation of Rab1 (111.1%), STAT3 (119.4%), PLCβ2 (115.8%), c-Fos (110.8%), and p-PKCα (110%). Rab1 is a miR-365a-3p target protein, but it was unexpectedly upregulated compared to the reference data.

**The NFkB signaling** was suppressed by downregulating miR-365a-3p target proteins NFkB (93.5%), p38 (85.7%), p-p38 (90.9%), SRC1 (94.2%), and NFAT5 (78.8%) and miR-365a-3p-responsible protein mTOR (88.1%), as well as coincidentally downregulating IKK (94.1%), MDR (89%), and NFATc1 (83%).

**The protection signaling** was suppressed by downregulating miR-365a-3p target proteins FOXP (94.4%), NOS1 (89%), FOXO1 (85.6%), and NRF2 (75.7%), as well as coincidentally downregulating leptin (94.7%), HO1 (92.4%), HSP27 (89.4%), hepcidin (87%), SVCT2 (83%), and SOD1 (71.6%), despite the upregulation of HSP90 (130%), FOXO3 (123.3%), ALDH1A1 (109.5%), HSP70 (108.4%), and GSTO1/2 (106.5%). FOXO3 is a miR-365a-3p target protein, but it was unexpectedly upregulated compared to the reference data.

**The ER stress signaling** was suppressed by downregulating miR-365a-3p target proteins eIF2AK3 (88.4%) and ATF6 (81.8%), as well as coincidentally downregulating HSP27 (89.4%), GADD153 (81.3%), ATF4 (80.8%), and LC3β (91.7%), despite the upregulation of eIF2α (119.5%), p-GADD153 (119.5%), BIP (111.3%), and IRE1α (114%).

**The innate immunity signaling** appear to be inactivated by downregulating miR-365a-3p target proteins TLR2 (93.4%), as well as coincidentally downregulating CAMP (90.6%), β-defensin1 (90.6%), β-defensin3 (94.2%), and elafin (84.4%), despite the upregulation of β-defensin2 (137.6%) and TLR4 (115.8%).

**The cellular immunity signaling** was suppressed by downregulating miR-365a-3p target proteins CD44 (92%) and CD54 (93.9%), as well as coincidentally downregulating CD4 (92.6%), CD8 (83.9%), CD20 (94.3%), CD34 (89.8%), CD40 (92.4%), granzyme B (77.2%), and CTLA4 (86.8%), despite the upregulation of CD99 (129.4%), versican (122.3%), perforin (111.1%), PDL1 (108.7%), and LTA4H (107.3%).

**The chronic inflammation signaling** appears to be inactivated by downregulating miR-365a-3p target proteins IL6 (79.9%) and M-CSF (90.7%) and miR-365a-3p-responsible protein COX2 (84.1%), as well as coincidentally downregulating CD68 (94.8%), MMP10 (83.3%), MMP12 (78%), and cathepsin K (92.9%), despite the upregulation of IL12 (105.4%), lysozyme (110.9%), MMP3 (130.4%), MMP9 (109.6%), TIMP2 (110.9%), COX1 (120.7%), and kininogen (140.1%). MMP3 is a miR-365a-3p target protein, but it was unexpectedly upregulated compared to the reference data.

**The FAS-mediated apoptosis signaling** was suppressed by downregulating miR-365a-3p target proteins BID (94.5%), as well as coincidentally downregulating FAS (86%), FADD (79.8%), and c-caspase3 (81.1%), despite the upregulation of FASL (121%) and caspase3 (111.1%) and coincidental downregulation of FLIP (88.9%).

**The glycolysis signaling** appears to be suppressed by downregulating miR-365a-3p target protein HK2 (85.5%) and LGR4 (82.1%), as well as coincidentally downregulating GAPDH (83.5%).

Conversely, **the miRNA biogenesis signaling** appears to be activated by upregulating Drosha (108.3%), Exportin5 (120.2%), TRBP2 (135%), and GW182 (106.7%) and coincidentally downregulating MTDH (94.5%), despite the downregulation of miR-365a-3p target protein Dicer (79.8%) and AGO2 (88.7%).

**The osteogenesis signaling** appears to be activated by upregulating BMP4 (120.5%), OPG (110.2%), PTH/PTHrPR (105.6%), RANKL (105.4%), aggrecan (112.1%), despite the downregulation of miR-365a-3p target protein RUNX2 (87.6%) and coincidental downregulation of osteocalcin (81.9%) and osteonectin (84.2%).

**The neuromuscular differentiation signaling** appears to be activated by upregulating NK1R (105.1%), S100 (110%), myosin-1a (112.6%), and MYH2 (111.2%), despite the downregulation of αSMA (86.1%).

**The angiogenesis signaling** appears to be activated by upregulating VEGFR2 (117.4%), TEM8 (124.2%), CMG2 (106.4%), and PDGFA (107.1%), despite the downregulation of miR-365a-3p target proteins FLT4 (87.5%), LYVE1 (87.2%), and FGF2 (93.4%), as well as coincidental downregulation of vWF (83.8%).

**The survival signaling** appears to be activated by upregulating SP1 (105.6%), sirtuin1 (110.2%), and sirtuin7 (106.8%), despite the downregulation of miR-365a-3p target protein XIAP (89.6%) and coincidental downregulation of sirtuin6 (87.1%). Sirtuin1 is a miR-365a-3p target protein, but it was unexpectedly upregulated compared to the reference data.

**The acute inflammation signaling** appears to be activated by upregulating TNFα (107.7%), TRAF6 (111.8%), IL8 (119.5%), and MMP1 (115.9%), despite the downregulation of miR-365a-3p target protein CRP (89%) and coincidental downregulation of MIF (78.1%) and CXCR4 (86.7%). TRAF6 is a miR-365a-3p target protein, but it was unexpectedly upregulated compared to the reference data.

**The p53-mediated apoptosis signaling** appears to be activated by upregulating p53 (108.9%), p63 (108.8%), p73 (112.1%), NOXA (105.1%), caspase9 (110.1%), and caspase3 (111.1%), despite the downregulation of miR-365a-3p target proteins MDM2 (96%), BAX (78.9%), and APAF1 (88.1%), and coincidental downregulation of c-caspase3 (81.1%). P53, p63, and p73 are miR-365a-3p target proteins, but they were unexpectedly upregulated compared to the reference data.

**The PARP-mediated apoptosis signaling** was activated by upregulating miR-365a-3p-responsible proteins PARP1 (118.4%) and c-PARP (123.6%), as well as coincidentally upregulating AIF1 (105%).

**The fibrosis signaling** appears to be activated by upregulating collagen 3A1 (107.3%), CTGF (110.7%), laminin α5 (107.7%), and integrin α5 (130.4%) and coincidentally downregulating FGF7 (86.7%), despite the downregulation of miR-365a-3p target proteins FGF1 (94.9%), FGF2 (93.4%), and tenascin C (93.8%), as well as coincidental downregulation of integrin β1 (92.4%), uPA (88.9%), and PAI1 (86.2%). Integrin α5 is a miR- 365a-3p target protein, but it was unexpectedly upregulated compared to the reference data.

**The senescence signaling** appears to be activated by upregulating klotho (114.7%), sirtuin1 (110.2%), and sirtuin7 (106.8%), despite the despite the downregulation of sirtuin6 (87.1%) and caveolin1 (94.8%). Sirtuin1 is a miR-365a-3p target protein, but it was unexpectedly upregulated compared to the reference data.

In addition, miR-365a-3p*-CEDS greatly influenced **the epigenetic modification signaling** by downregulating miR-365a-3p target proteins EZH2 (83.9%), as well as the coincidental downregulation of histone H1 (93.6%), KDM4D (93.7%), HDAC10 (94%), MBD4 (91.9%), DMAP1 (65.5%), DNMT1 (91.3%), PCAF (92.5%), BRG1 (77.7%), Ac-lysine (91.9%), and lamin A/C (94.4%), while it upregulated HMGA2 (118%) and HMGB1 (106.1%). HMGA2 is a miR-365a-3p target protein, but it was unexpectedly upregulated compared to the reference data. Consequently, miR-365a-3p*-CEDS induced a trend towards a reduction in the methylation of histones and DNAs, and attenuation of transcriptional repression.

MiR-365a-3p*-CEDS significantly attenuated **the oncogenesis signaling** by downregulating miR-365a-3p target proteins BMI1 (65%), BACH1 (64.8%), as well as the coincidental downregulation of EWSR1 (91.9%), XBP1 (88.4%), HER2 (85.1%), and CEA (83.2%), while it widely enhanced **the oncogenesis signaling** by upregulating CRIP1 (129.9%), MTA2 (121%), NF1 (115.8%), ATR (117.4%), ZEB1 (119.7%), MALT1 (111.8%), BRCA1 (109.9%), TWIST1 (109.6%), cMyb (109.4%), YAP1 (109%), and KLF4 (106.7%). MTA2, NF1, ZEB1, MALT1, and YAP1 are miR-365a-3p target proteins, but they were unexpectedly upregulated compared to the reference data. Among the 28 oncoproteins, 11 were found to be overexpressed, six were underexpressed, and 11 exhibited minimal change in expression compared to the untreated controls.

Consequently, miR-365a-3p*-CEDS significantly affected the entire protein signaling in RAW 264.7 cells. MiR-365a-3p*-CEDS suppressed the main RAS-NFkB signaling axis, and subsequently inactivated the proliferation-related signaling axis, epigenetic modification-protection-ER stress signaling axis, and FA-mediated apoptosis-glycolysis signaling axis. Conversely, it activated angiogenesis-survival-senescence-acute inflammation signaling axis, and p53- and PARP-mediated apoptosis-fibrosis-oncogenesis signaling axis (Fig. 17B Table 10).

**Table 10.**
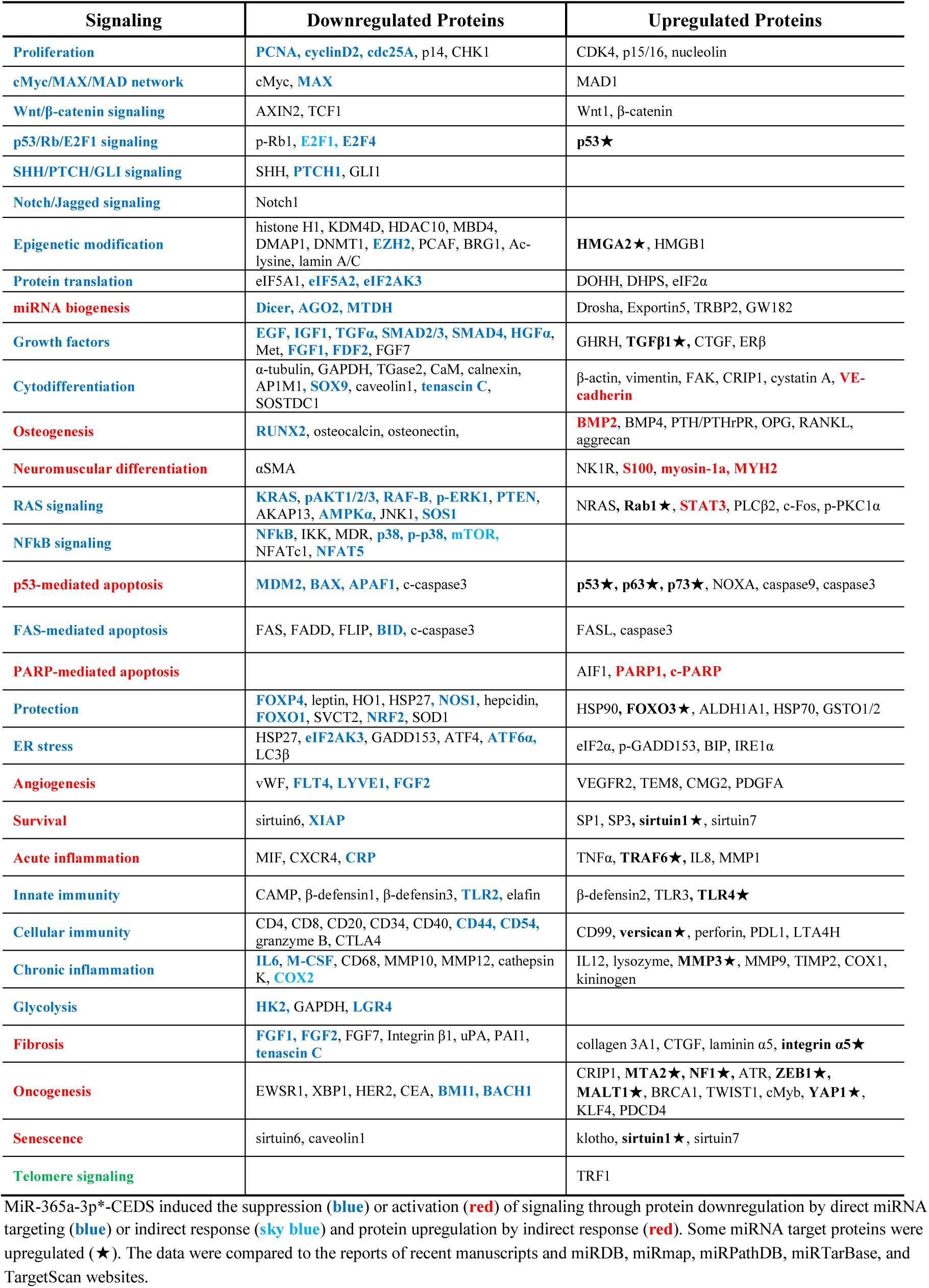
Protein expressions in the RAW 264.7 cells treated with miR-365a-3p*-CEDS.

The results indicate that miR-365a-3p*-CEDS exerted an anti-oncogenic effect on RAW 264.7 cells by suppressing proliferation, ROS protection, ER stress, and chronic inflammation. Conversely, CEDS also induced an oncogenic effect by increasing angiogenesis, survival, senescence, fibrosis, and oncogenesis. It is also proposed that miR-365a-3p*-CEDS stimulated p53- and PARP-mediated apoptosis signaling, while simultaneously attenuating the innate and cellular immunity signaling.

#### MiR-655-3p*-CEDS influenced the protein signaling in RAW 264.7 cells

RAW 264.7 cells treated with mmu-miR-655-3p sequence ATAATACATGGTTAACCTCTTT*-CEDS showed the characteristic protein expressions, resulting in the suppression of, Wnt/β-catenin signaling, p53/Rb/E2F1 signaling, SHH/PTCH/GLI signaling, Notch/Jagged signaling, protein translation, miRNA biogenesis, growth factors, cytodifferentiation, RAS signaling, NFkB signaling, protection, ER stress, survival, innate immunity, cellular immunity, p53-mediated apoptosis, FAS-mediated apoptosis, and glycolysis signaling, as well as the activation of proliferation, cMyc/MAX/MAD network, epigenetic modification, osteogenesis, neuromuscular differentiation, angiogenesis, acute inflammation, chronic inflammation, PARP-mediated apoptosis, fibrosis, oncogenesis, senescence, and telomere signaling compared to the untreated controls (Fig. 18A).

**Fig. 18.**
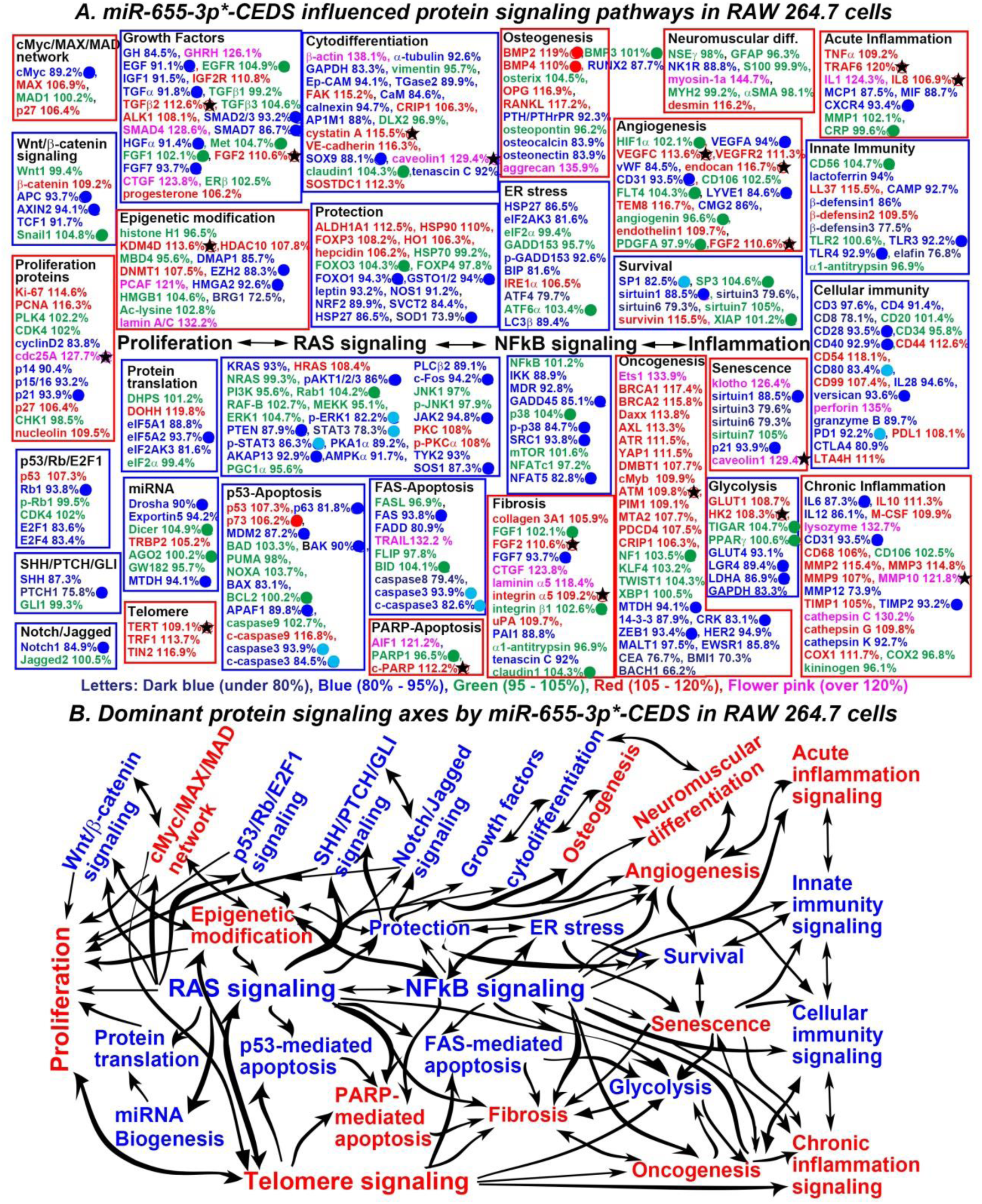
MiR-655-3p*-CEDS influenced the protein signaling (A) and axes (B) in RAW 264.7 cells. The IP- HPLC revealed proteins downregulated 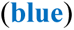, upregulated 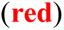, and minimally changed 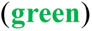 compared to the untreated controls. Dominantly suppressed 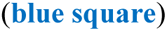 and activated 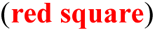 signaling. Downregulated proteins by direct targeting 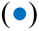 and indirect response 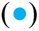, upregulated proteins by indirect response 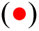, and minimal changed (±5%) proteins 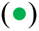. Some target proteins 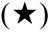 were unexpectedly upregulated contrast to the current manuscripts and data from miRDB and TargetScan websites.

**The Wnt/β-catenin signaling** was suppressed by downregulating miR-655-3p target proteins APC (93.7%), AXIN2 (94.1%), and TCF1 (91.7%), despite the upregulation of β-catenin (109.2%).

**The p53/Rb/E2F1 signaling** was suppressed by downregulating miR-655-3p target protein Rb1 (93.8%) and coincidentally downregulating E2F1 (83.6%), despite the upregulation of p53 (107.3%) and coincidental downregulation of E2F4 (83.4%).

**The SHH/PTCH/GLI signaling** was suppressed by downregulating miR-655-3p target protein PTCH1 (75.8%) and coincidentally downregulating SHH (87.3%).

**The Notch/Jagged signaling** was suppressed by downregulating miR-655-3p target protein Notch1 (84.9%).

**The protein translation signaling** was suppressed by downregulating miR-655-3p target protein eIF5A2 (93.7%) and coincidentally downregulating eIF5A1 (88.8%) and eIF2AK3 (81.6%), despite the upregulation of DOHH (119.8%).

**The miRNA biogenesis signaling** was suppressed by downregulating miR-655-3p target protein Drosha (90%) and coincidentally downregulating Exportin5 (94.2%), despite the upregulation of TRBP2 (105.2%) and downregulating miR-655 target protein MTDH (94.1%).

**The growth factor signaling** were partially suppressed by downregulating miR-655-3p target proteins EGF (91.1%), TGFα (91.8%), SMAD2/3 (93.2%), SMAD7 (86.7%), HGFα (91.4%), and FGF7 (93.7%), as well as coincidentally downregulating GH (84.5%) and IGF1 (91.5%), despite the upregulation of GHRH (126.1%), IGF2R (110.8%), TGFβ2 (112.6%), ALK1 (108.1%), SMAD4 (128.6%), FGF2 (110.6%), CTGF (123.8%), and progesterone (106.2%). TGFβ2 and FGF2 are miR-655-3p target proteins, but they were unexpectedly upregulated compared to the reference data.

**The cytodifferentiation signaling** were partially suppressed by downregulating miR-655-3p target protein SOX9 (88.1%), as well as coincidentally downregulating α-tubulin (92.6%), GAPDH (83.3%), Ep-CAM (94.1%), TGase2 (89.9%), CaM (84.6%), calnexin (94.7%), AP1M1 (88%), and tenascin C (92%), despite the upregulation of β-actin (138.1%), FAK (115.2%), CRIP1 (106.3%), cystatin A (115.5%), VE-cadherin (116.3%), caveolin1 (129.4%), and SOSTDC1 (112.3%).

**The RAS signaling** was suppressed by downregulating miR-655-3p target proteins pAKT1/2/3 (86%), PTEN (87.9%), AKAP13 (92.9%), c-Fos (94.2%), JAK2 (94.8%), and SOS1 (87.3%) and miR-655-3p-responsible proteins p-ERK1 (92.2%), STAT3 (78.3%), and p-STAT3 (86.3%), as well as coincidentally downregulating KRAS (93%), PKA1α (89.2%), AMPKα (91.7%), PLCβ2 (89.1%), and TYK2 (93%), despite the upregulation of HRAS (108.4%), PKC (108%), and p-PKCα (108%).

**The NFkB signaling** was suppressed by downregulating miR-655-3p target proteins GADD45 (85.1%), p- p38 (84.7%), SRC1 (93.8%), and NFAT5 (82.8%), as well as coincidentally downregulating IKK (88.9%) and MDR (92.8%).

**The protection signaling** was suppressed by downregulating miR-655-3p target proteins FOXO1 (94.3%), GSTO1/2 (94%), and SOD1 (73.9%), as well as coincidentally downregulating leptin (93.2%), NOS1 (91.2%), NRF2 (89.9%), SVCT2 (84.4%), and HSP27 (86.5%), despite the upregulation of ALDH1A1 (112.5%), HSP90 (110%), FOXP (108.2%), HO1 (106.3%), and hepcidin (106.2%).

**The ER stress signaling** was suppressed by downregulating HSP27 (86.5%), eIF2α (81.6%), p-GADD153 (92.6%), BIP (81.6%), ATF4 (79.7%), and LC3β (103.4%), despite the upregulation of IRE1α (106.5%).

**The survival signaling** was suppressed by downregulating miR-655-3p target protein sirtuin1 (88.5%) and miR-655-3p-responsible protein SP1 (82.5%), as well as coincidentally downregulating sirtuin3 and sirtuin7, despite the upregulation of survivin (115.5%).

**The innate immunity signaling** appears to be inactivated by downregulating miR-655-3p target proteins TLR3 (92.2%) and TLR4 (92.9%), as well as coincidentally downregulating lactoferrin (94%), CAMP (92.7%), β-defensin1 (86%), and β-defensin3 (77.5%), despite the upregulation of LL37 (115.5%) and β-defensin2 (109.5%).

**The cellular immunity signaling** was suppressed by downregulating miR-655-3p-responsible proteins CD28 (93.5%), CD40 (92.9%), and versican (93.6%) and miR-655-3p-responsible protein CD80 (83.4%), as well as coincidentally downregulating CD3 (97.6%), CD4 (91.4%), CD8 (78.1%), IL28 (94.6%), granzyme B (89.7%), and CTLA4 (80.9%), despite the upregulation of CD44 (112.6%), CD54 (118.1%), CD99 (107.4%), perforin (135%), PDL1 (108.1%), and LTA4H (111%).

**The p53-mediated apoptosis signaling** was suppressed by downregulating miR-655-3p target proteins p63 (81.8%), MDM2 (87.2%), BAK (90%), and APAF1 (89.8%), and miR-655-3p-responsible proteins caspase3 (93.9%) and c-caspase3 (84.5%), as well as coincidentally downregulating BAX (83.1%), despite the upregulation of p53 (107.3%) and p73 (106.2). P73 is a miR-655-3p target protein, but it was unexpectedly upregulated compared to the reference data.

**The FAS-mediated apoptosis signaling** was suppressed by downregulating miR-655-3p target protein FAS (93.8%) and miR-655-3p-responsible proteins caspase3 (93.9%) and c-caspase3 (82.6%), as well as coincidentally downregulating FADD (80.9%) and caspase8 (79.4%), despite the upregulation of TRAIL (132.2%).

**The glycolysis signaling** was suppressed by downregulating miR-655-3p target proteins LGR4 (89.4%) and LDHA (86.9%), as well as coincidentally downregulating GLUT4 (93.1%) and GAPDH (83.3%), despite the slight upregulation of GLUT1 (108.7%) and HK2 (108.3%). HK2 is a miR-655-3p target protein, but it was unexpectedly upregulated compared to the reference data.

Conversely, **the proliferation signaling** appears to be activated by upregulating Ki-67 (114.6%), PCNA (116.3%), cdc25A (127.7%), and nucleolin (109.5%) and downregulating miR-655-3p target protein p21 (93.9%), as well as coincidental downregulation p14 (90.4%) and p15/16 (93.2%), despite the downregulation of cyclin D2 (83.8%) and upregulation of p27 (106.4%). Cdc25A is a miR-655-3p target protein, but it was unexpectedly upregulated compared to the reference data.

**The cMyc/MAX/MAD network** appears to be activated by upregulating MAX (106.9%) and p27 (106.4%), despite the downregulation of miR-655-3p target protein cMyc (89.2%).

**The osteogenesis signaling** appears to be activated by upregulating miR-655-3p-responsible proteins BMP2 (119%) and BMP4 (110%) and coincidentally upregulating OPG (116.9%), RANKL (117.2%), and aggrecan (135.9%), despite the downregulation of RUNX2 (87.7%), PTH/PTHrPR (92.3%), osteocalcin (83.9%), and osteonectin (83.9%).

**The neuronal differentiation signaling** appears to be inactivated by downregulating NK1R (88.8%), while **the muscular differentiation signaling** was activated by upregulating myosin-1a (144.7%) and desmin (116.2%).

**The angiogenesis signaling** appeared to be activated by upregulating VEGFC (113.6%), VEGFR2 (111.3%), endocan (116.7%), TEM8 (116.7%), endothelin1 (109.7%), and FGF2 (110.6%), despite the downregulation of miR-655-3p target proteins VEGFA (94%), CD31 (93.5%), and LYVE1 (84.6%), as well as coincidental downregulation of vWF (84.5%) and CMG2 (86%). VEGFC, endocan, and FGF2 are miR-655-3p target proteins, but they were unexpectedly upregulated compared to the reference data.

**The acute inflammation signaling** appears to be activated by upregulating TNFα (109.2%), TRAF6 (120%), IL1 (124.3%), and IL8 (106.9%), despite the downregulation of miR-655-3p target protein CXCR4 (93.4%) and coincidental downregulation of MCP1 (87.5%) and MIF (88.7%). TRAF6 and IL8 are miR-655-3p target proteins, but they were unexpectedly upregulated compared to the reference data.

**The chronic inflammation signaling** appears to be activated by upregulating M-CSF (109.9%), lysozyme (132.7%), CD68 (106%), MMP2 (115.4%), MMP3 (114.8%), MMP9 (107%), MMP10 (121.8%), TIMP1 (105%), cathepsin C (130.2%), cathepsin G (109.8%), and COX1 (11.7%), despite the downregulation of miR- 655-3p target proteins IL6 (87.3%), CD31 (93.5%), and TIMP2 (93.2%) and coincidental downregulation of IL12 (86.1%), MMP12 (73.9%), and cathepsin K (92.7%). MMP10 is a miR-655-3p target protein, but it was unexpectedly upregulated compared to the reference data.

**The PARP-mediated apoptosis signaling** was activated by upregulating AIF1 (121.2%) and c-PARP (112.2%). C-PARP is a miR-655-3p target protein, but it was unexpectedly upregulated compared to the reference data.

**The fibrosis signaling** appears to be activated by upregulating collagen 3A1 (105.9%), FGF2 (110.6%), CTGF (123.8%), laminin α5 (118.4%), integrin α5 (109.2%), and uPA (109.7%) and coincidentally downregulating miR-655-3p target protein FGF7 (93.7%), despite the downregulation of PAI1 (88.8%) and tenascin C (92%). FGF2 and integrin α5 are miR-655-3p target proteins, but they were unexpectedly upregulated compared to the reference data.

**The senescence signaling** appears to be aberrant by downregulating miR-655-3p target protein sirtuin1 (88.5%) and p21 (93.9%) and coincidentally downregulating sirtuin3 (79.6%) and sirtuin6 (79.3%), even though the upregulation of klotho (126.4%) and caveolin1 (129.4%). Caveolin1 is a miR-655-3p target protein, but it was unexpectedly upregulated compared to the reference data.

**The telomere signaling** was activated by upregulating TERT (109.1%), TRF1 (113.7%), and TIN2 (116.9%). TERT is a miR-655-3p target protein, but it was unexpectedly upregulated compared to the reference data.

In addition, miR-655-3p*-CEDS significantly influenced **the epigenetic modification signaling** by downregulating miR-655-3p target proteins EZH2 (88.3%) and HMGA2 (92.6%) and coincidentally downregulating DMAP1 (85.7%) and BRG1 (72.5%), while it upregulated KDM4D (113.6%), HDAC10 (107.8%), PCAF (121%), and lamin A/C (132.2%). KDM4D is a miR-655-3p target protein, but it was unexpectedly upregulated compared to the reference data. Consequently, miR-655-3p*-CEDS induced a trend towards an increase in the methylation of histones and DNAs, and affected the transcriptional repression.

MiR-655-3p*-CEDS partially suppressed **the oncogenesis signaling** by downregulating miR-655-3p target proteins MTDH (94.1%), CRK (83.1%), and ZEB1 (93.4%), while it widely enhanced **the oncogenesis signaling** by upregulating Ets1 (133.9%), BRCA1 (117.4%), BRCA2 (115.8%), Daxx (113.8%), AXL (113.3%), ATR (111.5%), YAP1 (111.5%), DMBT1 (107.7%), cMyb (109.9%), ATM (109.8%), PIM1 (109.1%), MTA2 (107.7%), PDCD4 (107.5%), and CRIP (106.3%). ATM is a miR-655-3p target protein, but it was unexpectedly upregulated compared to the reference data. Among the 28 oncoproteins, 14 were found to be overexpressed, 10 were underexpressed, and four exhibited minimal change in expression compared to the untreated controls.

Consequently, miR-655-3p*-CEDS exhibited a variable effect on the entire protein signaling in RAW 264.7 cells. MiR-655-3p*-CEDS suppressed the RAS-NFkB signaling axis, subsequently inactivating the protection-ER stress-survival signaling axis and the p53- and FAS-mediated apoptosis signaling axis. Concurrently, miR-655-3p*-CEDS activated the epigenetic modification-cMyc/MAX/MAD network- proliferation signaling axis, epigenetic modification-telomere-PARP-mediated apoptosis-fibrosis signaling axis, angiogenesis-acute inflammation signaling axis, and oncogenesis-senescence-chronic inflammation signaling axis (Fig. 18B, Table 11).

**Table 11.** Protein expressions in the RAW 264.7 cells treated with miR-655-3p*-CEDS.

| Signaling | Downregulated Proteins | Upregulated Proteins |
| --- | --- | --- |
| **Proliferation** | cyclin D2, p14, p15/16, **p21** | Ki-67, PCNA, **cdc25A**★, p27, nucleolin |
| **cMyc/MAX/MAD network** | **cMyc** | MAX, p27 |
| Wnt/β-catenin signaling | APC, AXIN2, TCF1 | β-catenin |
| p53/Rb/E2F1 signaling | Rb1, E2F1, E2F4 | p53 |
| SHH/PTCH/GLI signaling | SHH, PTCH1 |  |
| Notch/Jagged signaling | Notch1 |  |
| **Epigenetic modification** | DMAP1, **EZH2**, PCAF, **HMG2A**, BRG1 | **KDM4D**★, HDAC10, lamin A/C |
| **Protein translation** | eIF5A1, **eIF5A2**, eIF2AK3 | DOHH |
| **miRNA biogenesis** | **Drosha**, Exportin5, **MTDH** | TRBP2 |
| Growth factors | GH, **EGF**, IGF1, **TGFα**, **SMAD2/3**, **SMAD7**, **HGFα**, **FGF7** | GHRH, EGFR, IGF2R, **TGFβ2**★, ALK1, SMAD4, Met, **FGF2**★, CTGF, progesterone |
| Cytodifferentiation | α-tubulin, GAPDH, claudin1, Ep-CAM, TGase2, CaM, calnexin, APIM1, **SOX9**, tenascin C | β-actin, FAK, CRIP1, **cystatin A**★, VE-cadherin, **caveolin1**★ |
| **Osteogenesis** | RUNX2, PTH/PTHrPR, osteocalcin, osteonectin | **BMP2**, **BMP4**, OPG, RANKL, aggrecan, SOSTDC1 |
| **Neuromuscular differentiation** | NK1R | myosin-1a |
| **RAS signaling** | KRAS, **pAKT1/2/3**, **p-ERK1**, **PTEN**, **STAT3**, **p-STAT3**, PKA1α, **AKAP13**, AMPKα, PLCβ2, **c-Fos**, **JAK2**, TYK2, **SOS1** | HRAS, PKC, p-PKCα |
| NFκB signaling | IKK, MDR, **GADD45**, **p-38**, **SRC1**, **NFAT5** |  |
| p53-mediated apoptosis | **p63**, **MDM2**, **BAK**, **BAX**, **APAF1**, **caspase3**, **c-caspase3** | p53, **p73**, c-caspase9 |
| FAS-mediated apoptosis | **FAS**, FADD, caspase8, **caspase3**, **c-caspase3** | TRAIL, BID |
| **PARP-mediated** |  | AIF1, **c-PARP**★ |
| **Protection** | **FOXO1**, **GSTO1/1**, leptin, NOS1, NRF2, SVCT2, HSP27, **SOD1** | ALDH1A1, FOXP3, HO1, hepcidin |
| **ER stress** | HSP27, eIF2AK3, GADD153, p-GADD153, BIP, ATF4, LC3β | IRE1α |
| **Angiogenesis** | **VEGFA**, vWF, **CD31**, **LYVE1**, CMG2 | **VEGFC**★, VEGFR2, **endocan**★, FLT4, TEM8, endothelin1, **FGF2**★ |
| **Survival** | **SP1**, **sirtuin1**, sirtuin3, sirtuin6 | survivin, |
| **Acute inflammation** | MCPI, MIF, **CXCR4** | TNFα, **TRAF6**★, IL1, **IL8**★ |
| **Innate immunity** | Lactoferrin, CAMP, β-defensin1, β-defensin3, **TLR3**, **TLR4**, elafin | CD56, LL37, β-defensin2, |
| **Cellular immunity** | CD3, CD4, CD8, **CD28**, **CD40**, **CD80**, IL28, **versican**, granzyme B, **PD1**, CTLA4 | CD44, CD54, CD99, perforin, PDL1, LTA4H |
| **Chronic inflammation** | **IL6**, IL12, **CD31**, MMP12, **TIMP2**, cathepsin K | IL10, M-CSF, lysozyme, CD31, CD68, MMP2, MMP3, MMP9, **MMP10**★, TIMP1, cathepsin C, cathepsin G, COX1 |
| **Glycolysis** | GLUT1, **LGR4**, **LDHA**, GAPDH | GLUT1, **HK2**★, TIGAR |
| **Fibrosis** | **FGF7**, PAI1, tenascin C | collagen 3A1, **FGF2**★, CTGF, laminin α5, **integrin α5**★, uPA, |
| **Oncogenesis** | **MTDH**, 14-3-3, **CRK**, **ZEB1**, HER2, MALT1, EWSR1, CEA, BMI1, BACH1 | Ets1, BRCA1, BRCA2, Daxx, AXL, ATR, YAP1, DMBT1, cMyb, **ATM**★, PIM1, MTA2, PDCD4, CRIP1 |
| **Senescence** | **sirtuin1**, sirtuin3, sirtuin6, **p21** | klotho, p21, **caveolin**★ |
| **Telomere signaling** |  | **TERT**★, TRF1, TIN2 |

MiR-655-3p\*-CEDS induced the suppression (blue) or activation (red) of signaling through protein downregulation by direct miRNA targeting (blue) or indirect response (sky blue) and protein upregulation by indirect response (red). Some miRNA target proteins were upregulated (★). The data were compared to the reports of recent manuscripts and miRDB, miRmap, miRPathDB, miRTarBase, and TargetScan websites.

The results indicate that miR-655-3p*-CEDS exerted an anti-oncogenic effect on RAW 264.7 cells by suppressing ROS protection, ER stress, survival, and glycolysis. However, it also markedly induced an oncogenic effect by increasing proliferation, telomere instability, fibrosis, angiogenesis, senescence, oncogenesis, and acute and chronic inflammation. It is also observed that miR-655-3p*-CEDS inhibited p53- and PARP-mediated apoptosis signaling, while simultaneously stimulating PARP-mediated apoptosis. Additionally, it impaired innate and cellular immunity signaling.

The above protein expression results after CEDS targeting nine mature miRNAs during RAW 264.7 cell culture could be summarized as follows. The RAW 264.7 cells treated with miR-26a-5p*-CEDS showed downregulation of 62 proteins, upregulation of 11 proteins, and minimal expressional change (±5%) of four proteins among 77 miR-26a-5p target proteins, which were identified by the current reports and the data on the miRDB and TargetScan websites. Consequently, the miRNA targeting ratio, defined as the number of downregulated proteins by targeting divided by the number of miR-26a-5p target proteins, was 80.5%. Additionally, CEDS indirectly decreased and increased the expression of 11 and 10 proteins, respectively, which were not targeted by miR-26a-5p (Table 12).

**Table 12.**
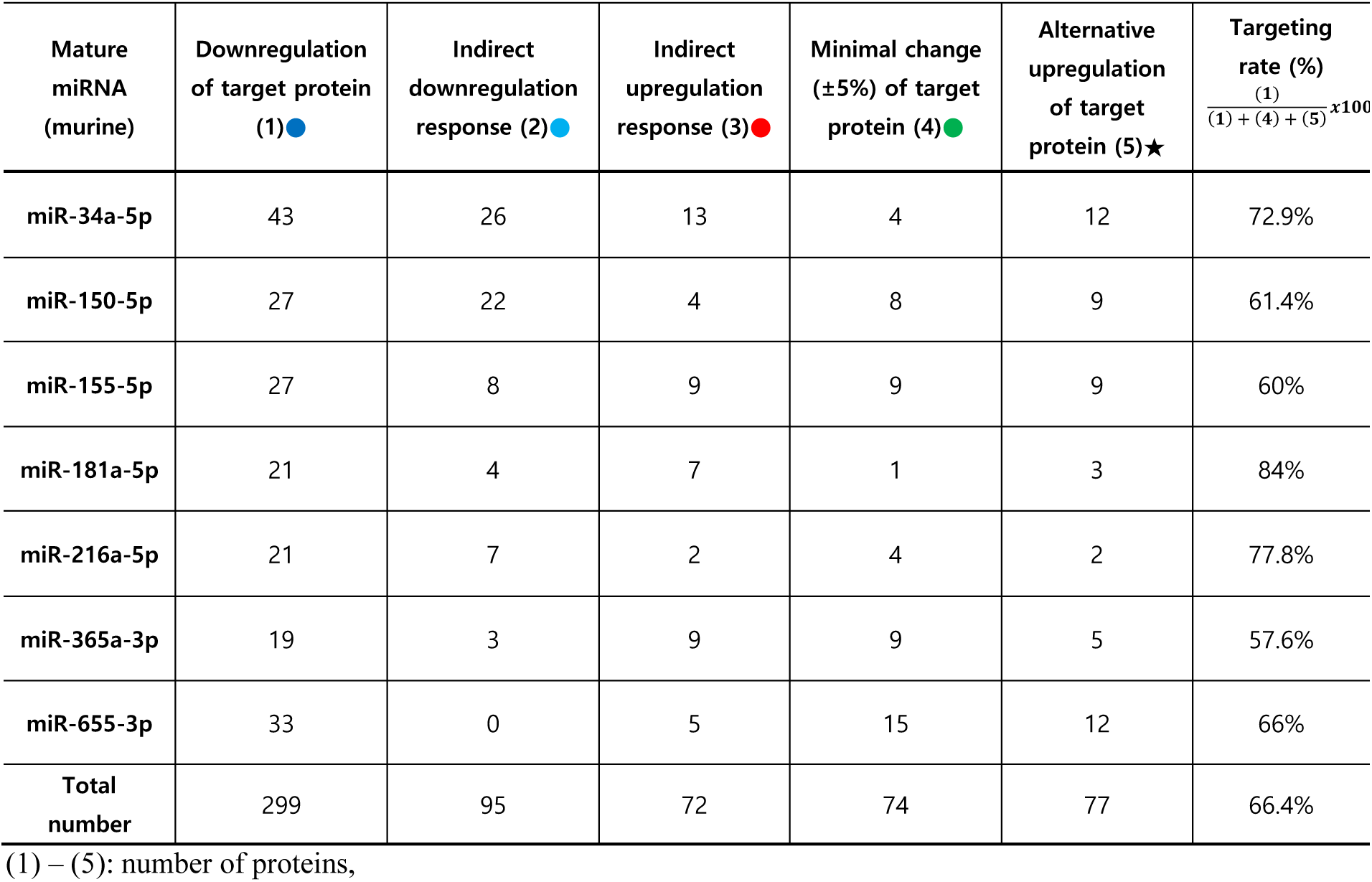
The efficiency of CEDS-induced miRNA targeting in the protein expression (n=350) of RAW 264.7 cells.

| Mature miRNA (murine) | Downregulation of target protein (1)● | Indirect downregulation response (2)● | Indirect upregulation response (3)● | Minimal change (±5%) of target protein (4)● | Alternative upregulation of target protein (5)★ | Targeting rate (%)<br>$\frac{(1)}{(1) + (4) + (5)} \times 100$ |
| --- | --- | --- | --- | --- | --- | --- |
| miR-34a-5p | 43 | 26 | 13 | 4 | 12 | 72.9% |
| miR-150-5p | 27 | 22 | 4 | 8 | 9 | 61.4% |
| miR-155-5p | 27 | 8 | 9 | 9 | 9 | 60% |
| miR-181a-5p | 21 | 4 | 7 | 1 | 3 | 84% |
| miR-216a-5p | 21 | 7 | 2 | 4 | 2 | 77.8% |
| miR-365a-3p | 19 | 3 | 9 | 9 | 5 | 57.6% |
| miR-655-3p | 33 | 0 | 5 | 15 | 12 | 66% |
| **Total number** | 299 | 95 | 72 | 74 | 77 | 66.4% |

(1) – (5): number of proteins,

The RAW 264.7 cells treated with miR-34a-5p*-CEDS showed downregulation of 43 proteins, upregulation of 12 proteins, and minimal expressional change (±5%) of four proteins among 59 miR-34a-5p target proteins, which were identified by the current reports and the data on the miRDB and TargetScan websites. Consequently, the miRNA targeting ratio, defined as the number of downregulated proteins by targeting divided by the number of miR-34a-5p target proteins, was 72.9%. Additionally, CEDS indirectly decreased and increased the expression of 26 and 13 proteins, respectively, which were not targeted by miR- 34a-5p (Table 12).

The RAW 264.7 cells treated with miR-126-5p*-CEDS showed downregulation of 46 proteins, upregulation of 14 proteins, and minimal expressional change (±5%) of 20 proteins among 80 miR-126-5p target proteins, which were identified by the current reports and the data on the miRDB and TargetScan websites. Consequently, the miRNA targeting ratio, defined as the number of downregulated proteins by targeting divided by the number of miR-126-5p target proteins, was 57.5%. Additionally, CEDS indirectly decreased and increased the expression of 15 and 13 proteins, respectively, which were not targeted by miR- 126-5p (Table 12).

The RAW 264.7 cells treated with miR-150-5p*-CEDS showed downregulation of 27 proteins, upregulation of nine proteins, and minimal expressional change (±5%) of eight proteins among 44 miR-150-5p target proteins, which were identified by the current reports and the data on the miRDB and TargetScan websites. Consequently, the miRNA targeting ratio, defined as the number of downregulated proteins by targeting divided by the number of miR-150-5p target proteins, was 61.4%. Additionally, CEDS indirectly decreased and increased the expression of 22 and four proteins, respectively, which were not targeted by miR- 150-5p (Table 12).

The RAW 264.7 cells treated with miR-155-5p*-CEDS showed downregulation of 27 proteins, upregulation of nine proteins, and minimal expressional change (±5%) of nine proteins among 45 miR-155-5p target proteins, which were identified by the current reports and the data on the miRDB and TargetScan websites. Consequently, the miRNA targeting ratio, defined as the number of downregulated proteins by targeting divided by the number of miR-155-5p target proteins, was 60%. Additionally, CEDS indirectly decreased and increased the expression of eight and nine proteins, respectively, which were not targeted by miR-155-5p sequence (Table 12).

The RAW 264.7 cells treated with miR-181a-5p*-CEDS showed downregulation of 21 proteins, upregulation of three proteins, and minimal expressional change (±5%) of one proteins among 25 miR-181a-5p target proteins, which were identified by the current reports and the data on the miRDB and TargetScan websites. Consequently, the miRNA targeting ratio, defined as the number of downregulated proteins by targeting divided by the number of miR-181a-5p target proteins, was 84%. Additionally, CEDS indirectly decreased and increased the expression of four and seven proteins, respectively, which were not targeted by miR-181a-5p (Table 12).

The RAW 264.7 cells treated with miR-216a-5p*-CEDS showed downregulation of 21 proteins, upregulation of two proteins, and minimal expressional change (±5%) of four proteins among 27 miR-216a-5p target proteins, which were identified by the current reports and the data on the miRDB and TargetScan websites. Consequently, the miRNA targeting ratio, defined as the number of downregulated proteins by targeting divided by the number of miR-216a-5p target proteins, was 77.8%. Additionally, CEDS indirectly decreased and increased the expression of seven and two proteins, respectively, which were not targeted by miR-216a-5p (Table 12).

The RAW 264.7 cells treated with miR-365a-3p*-CEDS showed downregulation of 19 proteins, upregulation of five proteins, and minimal expressional change (±5%) of nine proteins among 33 miR-365a-3p target proteins, which were identified by the current reports and the data on the miRDB and TargetScan websites. Consequently, the miRNA targeting ratio, defined as the number of downregulated proteins by targeting divided by the number of miR-365a-3p target proteins, was 57.6%. Additionally, CEDS indirectly decreased and increased the expression of three and nine proteins, respectively, which were not targeted by miR-365a-3p (Table 12).

The RAW 264.7 cells treated with miR-655-3p*-CEDS showed downregulation of 33 proteins, upregulation of 12 proteins, and minimal expressional change (±5%) of 15 proteins among 60 miR-655-3p target proteins, which were identified by the current reports and the data on the miRDB and TargetScan websites. Consequently, the miRNA targeting ratio, defined as the number of downregulated proteins by targeting divided by the number of miR-655-3p target proteins, was 66%. Additionally, CEDS indirectly increased the expression of five proteins, which were not targeted by miR-655-3p (Table 12).

As a result, the RAW 264.7 cells treated with CEDSs using nine miRNA sequences demonstrated downregulation of 299 proteins (66.4%), upregulation of 77 proteins (17.1%), and minimal expressional change (±5%) of 74 proteins (16.4%) among 450 miRNA target proteins, which were identified by the current reports and the data on the miRDB and TargetScan websites. Consequently, the targeting ratio of miRNA sequence*- CEDS, defined as the number of downregulated miRNA target proteins by CEDS divided by the number of miRNA target proteins examined in this study, was 66.4% on average, with a maximum of 84% and a minimum of 57.5%. In addition, CEDS indirectly decreased and increased the expression of 95 and 72 proteins, respectively, which were not targeted by CEDS examined in this study (Table 12).

The results indicate that CEDS using a miRNA sequence may significantly downregulate the target proteins by approximately 66.4% in ratio, as observed in this study. However, there was still marked variation in the nine cases of CEDS experiments due to the limitation of the number of each miRNA target proteins in this study.

### IP-HPLC analysis for the changes of the protein signaling after CEDS targeting different protein binding site sequences in RAW 264.7 cells

In addition to mature miRNA sequences, various protein binding site sequences located throughout genomic DNAs may also be favorable targets for CEDS. Among many protein binding site sequences in murine genomic DNA, two binding site sequences including p53 cluster 1 response element (RE) binding site sequence (AGACATGCCT) and canonical E2Fs binding site sequence (TTTC(C/G)CGC) were selected for the application of CEDS in this study.

#### The p53 binding site sequence (AGACATGCCT)*-CEDS influenced the protein signaling in RAW 264.7 cells

The consensus binding site of p53 response elements (REs) is comprised of two decameric DNA motifs, RRRCWWGYYY (R = A, G; W = A, T; Y = C, T), which are redundant, approximately palindromic, and separated by 0–18 bps^12^. In the present study, the decameric binding site sequence for p53 cluster 1 RE^21^, AGACATGCCT was selected for CEDS experiment on KE cells.

The IP-HPLC results demonstrated that the AGACATGCCT*-CEDS suppressed protein signaling, including SHH/PTCH/GLI signaling, Notch/Jagged signaling, miRNA biogenesis, growth factors, neuromuscular differentiation, RAS signaling, NFkB signaling, protection, cellular immunity, chronic inflammation, FAS-mediated apoptosis, and senescence signaling. Concurrently, CEDS activated protein signaling, such as proliferation, cMyc/MAX/MAD network, Wnt/β-catenin signaling, p53/Rb/E2F1 signaling, epigenetic modification, protein translation, cytodifferentiation, osteogenesis, angiogenesis, survival, acute inflammation, innate immunity, p53-mediated apoptosis, PARP-mediated apoptosis, fibrosis, glycolysis, and telomere signaling compared to the untreated controls (Fig. 21A).

**Fig. 21.**
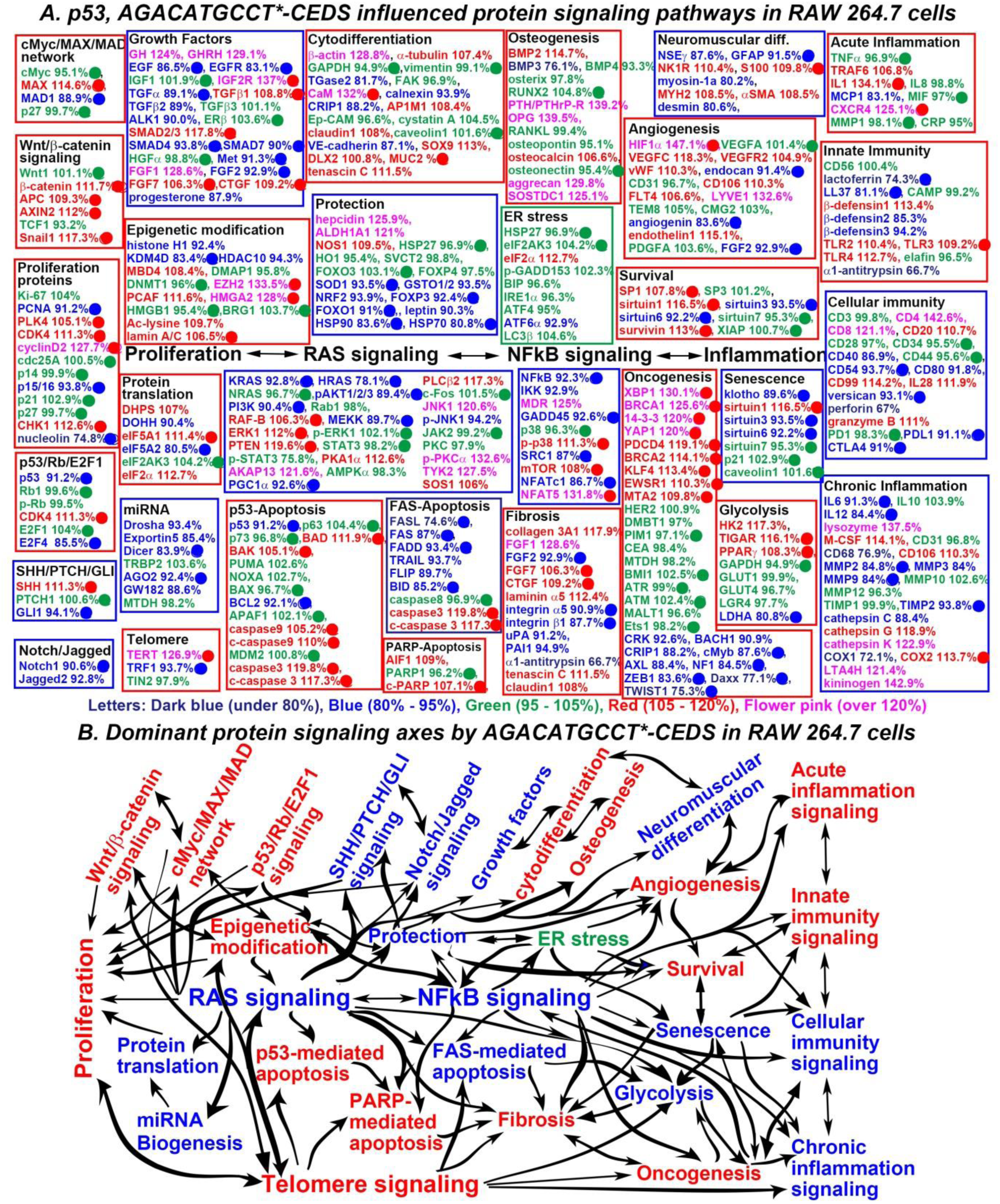
The p53 binding site sequence (AGACATGCCT)*-CEDS influenced the protein signaling (A) and axes (B) in RAW 264.7 cells. The IP-HPLC results revealed proteins that were downregulated 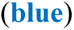, upregulated 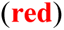, and minimally changed 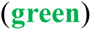 compared to the untreated controls. Dominant trend of suppressed 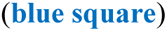 and activated 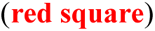 signaling. p53 target proteins (Harmonizome 3.0), downregulated 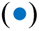 or upregulated 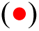 by AGACATGCCT*-CEDS.

**The SHH/PTCH/GLI signaling** appear to be inactivated by downregulating the p53 target protein (GLI1 94.1%), despite the upregulation of the p53 target protein SHH (111.3%).

**The Notch/Jagged signaling** was suppressed by downregulating the p53 target protein Notch1 (90.6%) and coincidentally downregulating Jagged2 (92.8%).

**The miRNA biogenesis signaling** was suppressed by downregulating the p53 target proteins Dicer (83.9%) and AGO2 (92.4%), as well as coincidentally downregulating Drosha (93.4%), Exportin5 (85.4%), and GW182 (88.6%).

**The growth factor signaling** partially was partially inactivated by downregulating the p53 target proteins EGF (86.5%), EGFR (83.1%), TGFα (89.1%), SMAD4 (93.8%), SMAD7 (90%), Met (91.3%), and FGF2 (92.9%) and coincidentally downregulating TGFβ2 (89%), ALK1 (90.0%), and progesterone (87.9%), despite the upregulation of the p53 target proteins IGF2R (137%), TGFβ1 (108.8%), SMAD2/3 (117.8%), FGF7 (106.3%), and CTGF (109.2%) and coincidental upregulation of GH (124%), GHRH (129.1%), and FGF1 (128.6%).

**The neuromuscular differentiation signaling** appears to be inactivated by downregulating the p53 target proteins GFAP (91.5%) and coincidentally downregulating NSEγ (87.6%), myosin-1a (80.2%), and desmin (80.6%), despite the upregulation of the p53 target protein S100 (109.8%) and coincidental upregulation of NK1R (110.4%), MYH2 (108.5%), and αSMA (108.5%).

**The RAS signaling** was suppressed by downregulating the p53 target proteins KRAS (92.8%), HRAS (78.1%), pAKT1/2/3 (89.4%), PI3K (90.4%), MEKK (89.7%), and PGC1α (92.6%) and coincidentally downregulating p-JNK1 (94.2%), despite the upregulation of the p53 target proteins RAF-B (106.3%), ERK1 (112%), and PTEN (119.6%) and coincidental upregulation of PKA1α (112.6%), AKAP13 (121.6%), PLCβ2 (117.3%), JNK1 (120.6%), p-PKCα (132.6%), TYK2 (127.5%), and SOS1 (106%).

**The NFkB signaling** appears to be inactivated by downregulating the p53 target proteins NFkB (92.3%), GADD45 (92.6%), SRC1 (87%), and NFATc1 (86.7%) and coincidentally downregulating IKK (92.9%), despite the upregulation of the p53 target proteins p-p38 (111.3%), mTOR (108%), and NFAT5 (131.8%) and coincidental upregulation of MDR (125%).

**The protection signaling** was inactivated by downregulating the p53 target proteins SOD1 (93.5%), FOXP3 (92.4%), FOXO1 (91%), HSP90 (83.6%), and HSP70 (80.8%), as well as coincidentally downregulating GSTO1/2 (93.5%), NRF2 (93.9%), and leptin (90.3%), despite the upregulation of hepcidin (125.9%), ALDH1A1 (121%), and NOS1 (109.5%).

**The cellular immunity signaling** appear to be inactivated by downregulating the p53 target proteins CD54 (93.7%) and versican (93.1%), and coincidental downregulation of CD40 (86.9%) and perforin (67%), despite the upregulation of CD4 (142.6%), CD8 (121.1%), CD20 (110.7%), CD99 (114.2%), IL28 (111.9%), and granzyme B (111%) and downregulation of the p53 target proteins PDL1 (91.1%) and CTLA4 (91%).

**The chronic inflammation signaling** appears to be suppressed by downregulating the p53 target proteins IL6 (91.3%), IL12 (84.4%), MMP2 (84.8%), MMP9 (84%), and TIMP2 (93.8%), as well as coincidentally downregulating CD68 (76.9%), cathepsin C (88.4%), and COX1 (72.1%), despite the upregulation of the p53 target protein COX2 (113.7%) and coincidental upregulation of lysozyme (137.5%), M-CSF (114.1%), CD106 (110.3%), cathepsin C (88.4%), and COX1 (72.1%).

**The FAS-mediated apoptosis signaling** was suppressed by downregulating the p53 target proteins FASL (74.6%), FAS (87%), FADD (93.4%), and BID (85.2%) and coincidentally downregulating TRAIL (93.7%) and FLIP (89.7%), despite the alternative upregulation of the p53 target proteins caspase3 (119.8%) and c-caspase3 (117.3%).

**The senescence signaling** was suppressed by downregulating the p53 target proteins klotho (89.6%), sirtuin3 (93.5%), and sirtuin6 (92.2%), despite the upregulation of the p53 target protein sirtuin1 (116.5%).

Conversely, **the proliferating signaling** appears to be activated by upregulating the p53 target proteins PLK4 (105.1%), CDK4 (111.3%), cyclinD2 (127.7%), and CHK1 (112.6%) and downregulating p15/16 (93.8%), despite the downregulation of PCNA (91.2%) and nucleolin (74.8%).

**The cMyc/MAX/MAD network** was activated by upregulating the p53 target proteins MAX (95.1%) and downregulating MAD1 (88.9%).

**The Wnt/β-catenin signaling** was activated by upregulating the p53 target proteins β-catenin (111.7%), APC (109.3%), AXIN2 (112%), and Snail1 (117.3%).

**The p53/Rb/E2F1 signaling** appears to be activated by upregulating the p53 target proteins CDK4 (111.3%) and downregulating p53 (91.2%).

**The protein translation signaling** was activated by upregulating the p53 target proteins eIF5A1 (111.4%) and coincidentally upregulating DHPS (107%) and eIF2α (112.7%), despite the downregulation of the p53 target protein eIF5A2 (80.5%) and coincidental downregulation of DOHH (90.4%).

**The cytodifferentiation signaling** were partially activated by upregulating the p53 target protein CaM (132%) and coincidentally upregulating β-actin (128.8%), α-tubulin (107.4%), AP1M1 (108.4%), claudin1 (108%), SOX9 (113%), DLX2 (100.8%), and tenascin C (111.5%), despite the downregulation of TGase2 (81.7%), calnexin (93.9%), and VE-cadherin (87.1%).

**The osteogenesis signaling** was activated by upregulating BMP2 (114.7%), PTH/PTHrP-R (139.2%), OPG (139.5%), osteocalcin (106.6%), and aggrecan (129.8%), despite the downregulation of BMP3 (76.1%) and upregulation of SOSTDC1 (125.1%).

**The angiogenesis signaling** was activated by upregulating the p53 target protein HIF1α (147.1%), as well as coincidentally upregulating VEGFC (118.3%), VEGFR2 (104.9%), vWF (110.3%), CD106 (110.3%), FLT4 (106.6%), LYVE1 (132.6%), and endothelin1 (115.1%), despite the downregulation of the p53 target protein endocan (91.4%), angiogenin (83.6%), and FGF2 (92.9%).

**The survival signaling** was activated by upregulating the p53 target proteins SP1 (107.8%), sirtuin1 (116.5%), and survivin (113%), despite the downregulation of the p53 target proteins sirtuin3 (93.5%) and sirtuin6 (92.2%).

**The acute inflammation signaling** was activated by upregulating the p53 target proteins IL1 (134.1%) and (CXCR4 125.1%), and coincidentally upregulating TRAF6 (106.8%), despite the downregulation of MCP1 83.1%.

**The innate immunity signaling** appears to be activated by upregulating the p53 target protein TLR3 (109.2%) and coincidentally upregulating TLR2 (110.4%), TLR4 (112.7%), and β-defensin1 (113.4%), despite the downregulation of the p53 target proteins lactoferrin (74.3%) and LL37 (81.1%), and coincidental downregulation of β-defensin2 (85.3%), β-defensin3 (94.2%), and α1-antitrypsin (66.7%).

**The p53-mediated apoptosis signaling** appears to be activated by upregulating the p53 target proteins BAD (111.9%), BAK (105.1%), caspase9 (105.2%), c-caspase9 (110%), caspase3 (119.8%), and c-caspase3 (117.3%), despite the downregulation of the p53 target proteins p53 (91.2%) and BCL2 (92.1%). **The PARP-mediated apoptosis signaling** was activated by upregulating the p53 target protein c-PARP (107.1%) and coincidentally upregulating AIF1 (109%).

**The fibrosis signaling** appears to be activated by upregulating the p53 target protein CTGF (109.2%), as well as coincidentally upregulating collagen 3A1 (117.9%), FGF1 (128.6%), laminin α5 (112.4%), tenascin C (111.5%), and claudin1 (108%) and upregulating the p53 target protein FGF7 (106.3%), despite the downregulation of the p53 target proteins FGF2 (92.9%), integrin α5 (90.9%), and integrin β1 (87.7%), and coincidental downregulation of uPA (91.2%), PAI1 (94.9%), and α1-antitrypsin (66.7%).

**The glycolysis signaling** was activated by upregulating the p53 target proteins TIGAR (116.1%), PPARγ (108.3%) and coincidentally upregulating HK2 (117.3%), despite the downregulation of the p53 target protein LDHA (80.8%).

**The telomere signaling** was activated by upregulating the p53 target protein TERT (126.9%), despite of slight downregulation of the p53 target protein TRF1 (93.7%).

In addition, the p53 binding site sequence*-CEDS significantly influenced **the epigenetic modification signaling** by downregulating the p53 target protein KDM4D (83.4%) and coincidentally downregulating histone H1 (92.4%) and HDAC10 (94.3%), while it upregulated the p53 target proteins EZH2 (133.5%), HMGA2 (128%), and lamin A/C (106.5%) and coincidentally upregulated MBD4 (108.4%), PCAF (111.6%), and Ac-lysine (109.7%). Consequently, the AGACATGCCT*-CEDS induced a trend towards an increase in the methylation of histones and DNAs, and transcriptional repression.

The p53 binding site sequence*-CEDS significantly suppressed **the oncogenesis signaling** by downregulating the p53 target proteins cMyb (87.6%), NF1 (84.5%), ZEB1 (83.6%), Daxx (77.1%), and TWIST1 (75.3%) and coincidentally downregulating CRK (92.6%), BACH1 (90.9%), CRIP1 (88.2%), and AXL (88.4%), while it widely enhanced **the oncogenesis signaling** by upregulating the p53 target proteins XBP1 (130.1%), BRCA1 (125.6%), 14-3-3 (120%), YAP1 (120%), PDCD4 (119.1%), BRCA2 (114.1%), KLF4 (113.4%), EWSR1 (110.3%), and MTA2 (109.8%). Among the 28 oncoproteins, nine were found to be overexpressed, nine were underexpressed, and ten exhibited minimal change in expression compared to the untreated controls.

Consequently, the p53 binding site sequence (AGACATGCCT)*-CEDS significantly impacted the protein signaling in RAW 264.7 cells. The AGACATGCCT*-CEDS primary suppressed the RAS and NFkB signaling axis, subsequently inactivated the protection-Notch/Jagged-SHH/PTCH/GLI signaling axis, FAS- and PARP-mediated apoptosis signaling axis, and senescence-glycolysis-cellular immunity-chronic inflammation signaling axis. In contrast, it enhanced the epigenetic modification-proliferation-related signaling axis, angiogenesis-survival-innate immunity-acute inflammation signaling axis, and epigenetic modification-telomere-p53- mediated apoptosis-fibrosis-oncogenesis signaling axis (Fig. 21B, Table 15).

**Table 15.** Protein expressions in the RAW 264.7 cells treated with p53 binding site sequence (AGACATGCCT)*-CEDS.

| Signaling | Downregulated Proteins | Upregulated Proteins |
| --- | --- | --- |
| Proliferation | PCNA, p15/16, nucleolin | PLK4, CDK4, cyclin D2, CHK1 |
| cMyc/MAX/MAD network | MAD1 | MAX |
| Wnt/β-catenin signaling | | β-catenin, APC, AXIN2, Snail |
| p53/Rb/E2F1 signaling | p53, E2F4 | CDK4 |
| SHH/PTCH/GLI signaling | GLI1 | SHH |
| Notch/Jagged signaling | Notch1, Jagged |  |
| Epigenetic modification | histone H1, KDM4D, HDAC10 | MBD4, EZH2, PCAF, HMG A2, Ac-lysine, lamin A/C |
| Protein translation | DOHH, eIF5A2 | DHPS, eIF5A1, eIF2α |
| miRNA biogenesis | Drosha, Exportin5, Dicer, AGO2, GW182 |  |
| Growth factors | EGF, EGFR, TGFα, TGFβ2, ALK1, SMA4, SMAD7, Met, FGF2, progesterone | GH, GHRH, IGF2R, TGFβ1, SMAD2/3, FGF1, FGF7, CTGF |
| Cytodifferentiation | TGase2, calnexin, CRIP1, VE-cadherin | β-actin, α-tubulin, CaM, AP1M1, claudin1, SOX9, DLX2, tenascin C |
| Osteogenesis | BMP3 | BMP2, PTH/PTHrPR, OPG, osteocalcin, aggrecan, SOSTDC1 |
| Neuromuscular differentiation | NSEγ, GFAP, myosin-1a, desmin | NK1R, S100, MYH2, αSMA |
| RAS signaling | KRAS, HRAS, pAKT1/2/3, PI3K, MEKK, PGC1α, p-JNK1 | RAF-B, ERK1, PTEN, PKA1α, AKAP13, PLCβ2, JNK1, p-PKC1α, TYK2, SOS1 |
| NFκB signaling | NFκB, IKK, GADD45, SRC1, NFATc1 | MDR, p-p38, mTOR, NFAT5 |
| p53-mediated apoptosis | p53, BCL2 | BAD, BAK, caspase9, c-caspase9, caspase3, c-caspase3 |
| FAS-mediated apoptosis | FASL, FAS, FADD, TRAIL, FLIP, BID | caspase3, c-caspase3 |
| PARP-mediated apoptosis |  | AIF1, c-PARP |
| Protection signaling | SOD1, GSTO1/2, NRF2, FOX P3, FOXO1, leptin, HSP90, HSP70 | hepcidin, ALDH1A1, NOS1 |
| ER stress signaling | ATF6α | eIF2α |
| Angiogenesis signaling | endocan, angiogenin, FGF2 | HIF1α, VEGFC, VEGFR2, vWF, CD106, FLT4, LYVE1, endothelin1 |
| Survival signaling | sirtuin3, sirtuin6 | SPI, sirtuin1, survivin |
| Acute inflammation | MCP1 | TRAF6, IL1, CXCR4 |
| Innate immunity | lactoferrin, LL37, β-defensin2, β-defensin3, α1-antitrypsin | β-defensin1, TLR2, TLR3, TLR4 |
| Cellular immunity | CD40, CD54, CD80, versican, perforin, PDL1, CTLA4 | CD4, CD8, CD20, CD99, IL28, granzyme B |
| Chronic inflammation | IL6, IL12, CD68, MMP2, MMP3, MMP9, TIMP2, cathepsin C, COX1 | Lysozyme, M-CSF, CD106, cathepsin G, cathepsin K, COX2, LTA4H, kininogen |
| Glycolysis | LDHA | HK2, TIGAR, PPARγ, |
| Fibrosis | FGF2, integrin α5, integrin β1, uPA, PAI1, α1-antitrypsin | collagen 3A1, FGF1, FGF7, CTGF, laminin α5, tenascin C claudin1 |
| Oncogenesis | CRK, BACH1, CRIP1, cMyb, AXL, NF1, ZEB1, Daxx, TWIST1 | XBPI, BRCA1, 14-3-3, YAP1, PDCD4, BRCA2, KLF4, EWSR1, MTA2 |
| Senescence | klotho, sirtuin3, sirtuin6 | sirtuin1 |
| Telomere signaling | TRF1 | TERT |

The p53 binding site sequence\*-CEDS, AGACATGCCT\*-CEDS induced the suppression (blue) and activation (red) of signaling by downregulating (blue) and upregulating (red) the proteins, respectively. The data were compared to the reports of recent manuscripts and Harmonizome 3.0 website.

The results indicate that the AGACATGCCT*-CEDS had both anti-oncogenic and oncogenic effects on RAW 264.7 cells. The anti-oncogenic effect was weak and was found to decrease ROS protection, senescence, and glycolysis. In contrast, the oncogenic effect was strong and was found to increase cellular proliferation, angiogenesis, survival, fibrosis, oncogenesis, and telomere instability. It was additionally observed that the AGACATGCCT*-CEDS can diminish the capacity of FAS- and PARP-mediated apoptosis and cellular immunity in the cells.

#### The canonical E2Fs binding site sequence (TTTC(C/G)CGC)*-CEDS influenced the protein signaling in RAW 264.7 cells

E2Fs are a family of transcription factors found in higher eukaryotes. They encode eight genes, with E2F1, E2F2, and E2F3a functioning as transcription activators and E2F3b, E2F4, E2F5, E2F6, E2F7, and E2F8 functioning as transcription repressors. On the other hand, the typical E2Fs with DP proteins, including E2F1, E2F2, E2F3a, E2F3b, E2F4, E2F5, and E2F6, are known to bind to a canonical sequence TTTC(C/G)CGC, in contrast to the atypical E2Fs with no DP protein, including E2F7 and E2F8^22^. The present study utilized the canonical E2Fs binding site sequence TTTC(C/G)CGC, for CEDS on RAW 264.7 cells. It is therefore anticipated that the TTTC(C/G)CGC*-CEDS will affect the transcriptional roles of both transcription activators (E2F1, E2F2, and E2F3a) and transcription repressors (E2F3b, E2F4, E2F5, and E2F6).

E2Fs are a family of transcription factors found in higher eukaryotes. They encode eight genes, with E2F1, E2F2, and E2F3a acting as transcriptional activators and E2F3b, E2F4, E2F5, E2F6, E2F7, and E2F8 acting as transcriptional repressors. On the other hand, the typical E2Fs with DP proteins, including E2F1, E2F2, E2F3a, E2F3b, E2F4, E2F5, and E2F6, are known to bind to a canonical sequence TTTC(C/G)CGC, in contrast to the atypical E2Fs with no DP protein, including E2F7 and E2F8. In the present study, the canonical E2F binding site sequence TTTC(C/G)CGC, was used for CEDS on RAW 264.7 cells. Therefore, it is expected that the TTTC(C/G)CGC*-CEDS will affect the transcriptional roles of both transcriptional activators (E2F1, E2F2, and E2F3a) and transcriptional repressors (E2F3b, E2F4, E2F5, and E2F6).

The IP-HPLC results demonstrated that the TTTC(C/G)CGC*-CEDS suppressed protein signaling, including the Wnt1/β-catenin signaling, SHH/PTCH/GLI signaling, protein translation, miRNA biogenesis, osteogenesis, ER stress, survival, innate immunity, FAS-mediated apoptosis, PARP-mediated apoptosis, fibrosis, oncogenesis, and glycolysis signaling. Concurrently, CEDS activated protein signaling, such as the proliferation, cMyc/MAX/MAD network, p53/Rb/E2F1 signaling, Notch/Jagged signaling, epigenetic modification, growth factors, cytodifferentiation, neuromuscular differentiation, RAS signaling, NFkB signaling, protection, angiogenesis, acute inflammation, cellular immunity, chronic inflammation, p53-mediated apoptosis, senescence, and telomere signaling compared to the untreated controls (Fig. 22A).

**Fig. 22.**
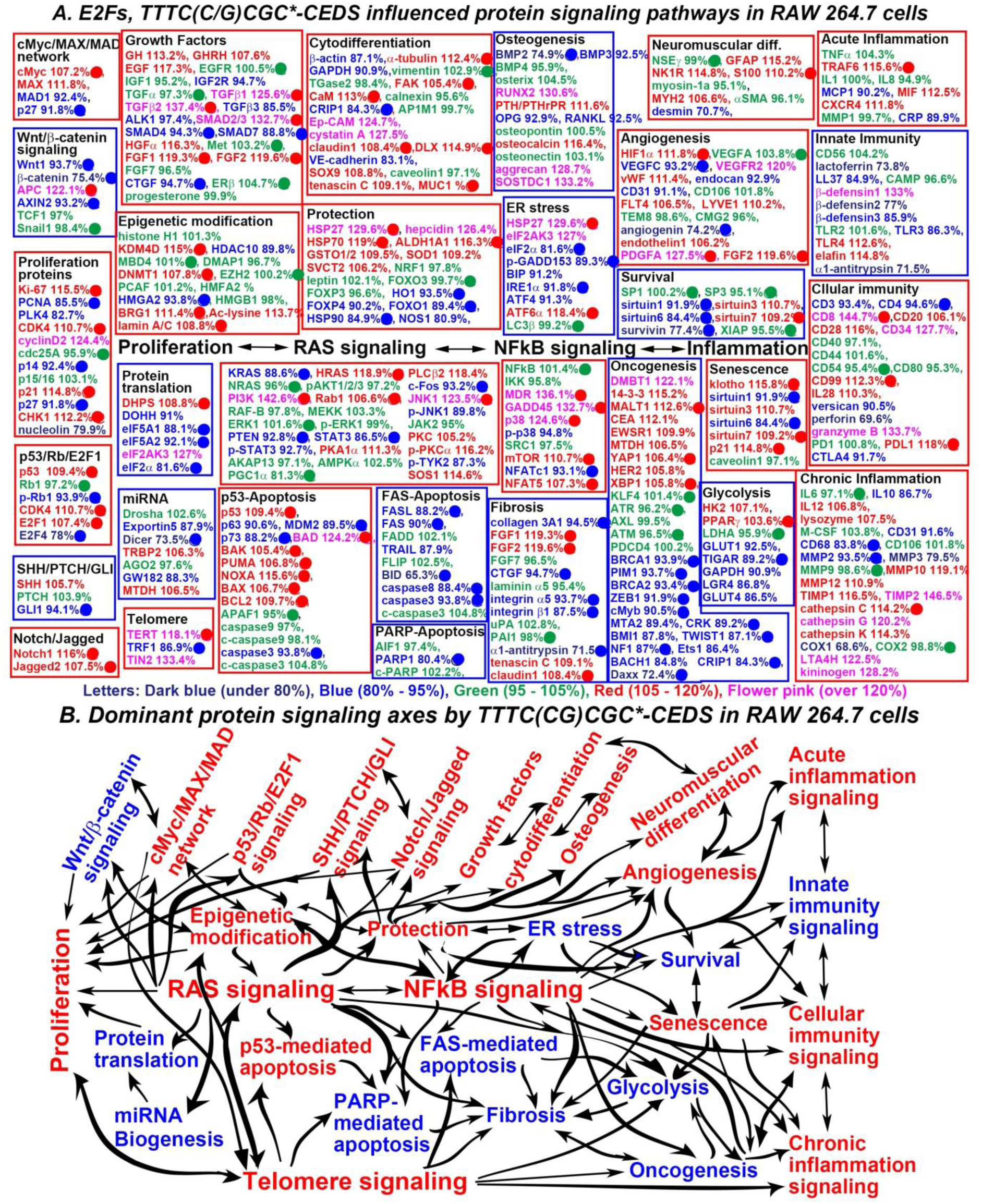
The E2Fs binding site sequence (TTTC(C/G)CGC)*-CEDS influenced the protein signaling (A) and axes (B) in RAW 264.7 cells. The IP-HPLC results revealed proteins downregulated 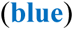, upregulated 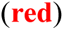, and minimally changed 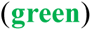 compared to the untreated controls. Dominant trend of suppressed 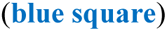 and activated 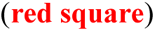 signaling to be defined. E2F1 target proteins (Harmonizome 3.0), downregulated 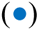 or upregulated 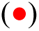 by (TTTC(C/G)CGC)*-CEDS.

However, the canonical E2Fs binding site sequence*-CEDS increased E2F1 expression (107.4%) while decreased E2F4 expression (78%) in RAW 264.7 cells. Therefore, this study focused on observing the expression changes of E2F1 targeting proteins in the cells after CEDS.

**The Wnt/β-catenin signaling** was suppressed by downregulating the E2F1 target proteins Wnt1 (93.7%), β-catenin (75.4%), AXIN2 (93.2%), despite the upregulation of the E2F1 target protein APC (122.1%).

**The protein translation signaling** was suppressed by downregulating the E2F1 target proteins eIF5A1 (88.1%), eIF5A2 (92.1%), and eIF2α (81.6%), despite the upregulation of the E2F1 target protein DHPS (108.8%) and coincidental upregulation of eIF2AK3 (127%).

**The miRNA biogenesis signaling** appears to be inactivated by downregulating the E2F1 target protein Dicer (73.5%) and coincidentally downregulating Exportin5 (87.9%) and GW182 (88.3%), despite the upregulation of TRBP2 (106.3%) and MTDH (106.5%).

**The osteogenesis signaling** appears to be inactivated by downregulating the E2F1 target protein BMP2 (74.9%) and coincidentally downregulating BMP3 (92.5%), OPG (92.9%), and RANKL (92.5%), as well as coincidentally upregulating SOSTDC1 (133.2%), despite the upregulation of RUNX2 (130.6%), PTH/PTHrPR (111.6%), osteocalcin (116.4%), and aggrecan (128.7%).

**The ER stress pathway** was suppressed by downregulating the E2F1 target proteins eIF2a (81.6%), p- GADD153 (89.3%), and IRE1α (91.8%) and coincidentally downregulating BIP (91.2%) and ATF4 (91.3%), despite the upregulation of the E2F1 target proteins HSP27 (129.6%) and ATF6α (118.4%), and the coincidental upregulation of eIF2AK3 (127%).

**The survival signaling** appears to be inactivated by downregulating the E2F1 target proteins sirtuin1 (91.9%), sirtuin6 (84.4%), and survivin (77.4%), despite the upregulation of the E2F1 target protein sirtuin7 (109.2%) and coincidental upregulation of sirtuin3 (110.7%).

**The innate immunity signaling** appears to be inactivated by downregulating lactoferrin (73.8%), LL37 (84.9%), β-defensin2 (77%), β-defensin3 (85.9%), TLR3 (86.3%), and α1-antitrypsin (71.5%), despite the coincidental upregulation of β-defensin1 (133%), TLR4 (112.6%), and elafin (114.8%).

**The FAS-mediated apoptosis signaling** was suppressed by downregulating the E2F1 target proteins FASL (88.2%), FAS (90%), BID (65.3%), caspase8 (88.4%), and caspase3 (93.8%) and coincidentally downregulating TRAIL (87.9%).

**The PARP-mediated apoptosis signaling** was suppressed by downregulating the E2F1 target protein PARP1 (80.4%).

**The fibrosis signaling** was suppressed by downregulating collagen 3A1 (90.9%), FGF1 (82.9%), laminin α5 (93.9%), integrin α5 (89.2%), integrin β1 (86.9%), α1-antitrypsin (91.6%), and tenascin C (91.3%).

**The glycolysis signaling** was activated by upregulating the E2F1 target proteins collagen 3A1 (94.5%), CTGF (94.7%), integrin α5 (93.7%), integrin β1 (87.5%), and α1-antitrypsin (71.5%), despite the downregulation of the E2F1 target proteins FGF1 (119.3%), FGF2 (119.6%), claudin1 (108.4%) and coincidental downregulation of tenascin C (109.1%).

Conversely, **the proliferating signaling** was activated by upregulating the E2F1 target proteins Ki-67 (115.5%), CDK4 (110.7%), p21 (114.8%), and CHK1 (112.2%), despite the downregulation of the E2F1 target proteins PCNA (85.5%), p14 (92.4%), and nucleolin (79.9%) and coincidental downregulation of PLK4 (82.7%).

**The cMyc/MAX/MAD network** was suppressed by downregulating the E2F1 target protein cMyc (107.2%) and coincidentally downregulating MAX (111.8%), as well as the downregulation of MAD1 (92.4%) and p27 (91.8%).

**The p53/Rb/E2F1 signaling** was activated by upregulating the E2F1 target proteins p53 (109.4%), CDK4 (110.7%), and E2F1 (107.4%) and downregulating the E2F1 target proteins p-Rb1 (93.9%) and E2F4 (78%).

**The SHH/PTCH/GLI signaling** appears to be activated by upregulating SHH (105.7%), despite downregulation of the E2F1 target protein GLI1 (94.1%).

**The Notch/Jagged signaling** was activated by upregulating the E2F1 target proteins Notch1 (116%) and Jagged2 (107.5%).

**The growth factor signaling** were partially activated by upregulating the E2F1 target proteins TGFβ1 (125.6%), TGFβ2 (137.4%), SMAD2/3 (132.7%), FGF1 (119.3%), and FGF2 (119.6%) and coincidentally upregulating GH (113.2%), GHRH (107.6%), EGF (117.3%), despite the downregulation of the E2F1 target proteins SMAD4 (94.3%), SMAD7 (88.8%), and CTGF (94.7%) and coincidental downregulation of IGF2R (94.7%), TGFβ3 (85.5%), and ALK1 (97.4%).

**The cytodifferentiation signaling** were partially activated by upregulating the E2F1 target proteins α-tubulin (112.4%), FAK (105.4%), CaM (113%), claudin1 (108.4%), DLX (114.9%) and coincidentally upregulating Ep-CAM (124.7%), cystatin A (127.5%), SOX9 (108.8%), tenascin C (109.1%), despite the downregulation of the E2F1 target protein CRIP1 (84.3%) and coincidental downregulation of β-actin (87.1%), GAPDH (90.9%), and VE-cadherin (83.1%).

**The RAS signaling** was weakly activated by upregulating the E2F1 target proteins HRAS (118.9%), PI3K (142.6%), Rab1 (106.6%), and JNK1 (123.5%) and coincidentally upregulating PKA1α (111.3%), PLCβ2 (118.4%), PKC (105.2%), p-PKCα (116.2%), and SOS1 (114.6%), despite the downregulation of the E2F1 target proteins KRAS (88.6%), PTEN (92.8%), STAT3 (86.5%), and c-Fos (93.2%), and coincidental downregulation of p-STAT3 (92.7%), p-JNK1 (89.8%), and p-TYK2 (87.3%).

**The NFkB signaling** was activated by upregulating the E2F1 target proteins MDR (136.1%), GADD45 (132.7%), p38 (124.6%), mTOR (110.7%), and NFAT5 (107.3%), despite the downregulation of the E2F1 target protein NFATc1 (93.1%) and coincidental downregulation p-p38 (94.8%).

**The protection signaling** appears to be activated by upregulating the E2F1 target proteins HSP27 (129.6%), HSP70 (119%), and ALDH1A1 (116.3%) and coincidentally upregulating hepcidin (126.4%), GSTO1/2 (109.5%), SOD1 (109.2%), and SVCT2 (106.2%), despite the downregulation of the E2F1 target protein HO1 (93.5%), FOXO1 (89.4%), and HSP90 (84.9%) and coincidental downregulation of FOXP4 (90.2%) and NOS1 (80.9%).

**The angiogenesis signaling** appears to be activated by upregulating the E2F1 target proteins HIF1α (111.8%), PDGFA (127.5%), and FGF2 (119.6%) and coincidentally upregulating VEGFR2 (120%), vWF (111.4%), FLT4 (106.5%), LYVE1 (110.2%), and endothelin1 (106.2%), despite the downregulation of the E2F1 target proteins VEGFC (93.2%) and angiogenin (74.2%), and coincidental downregulation of endocan (92.9%) and CD31 (91.1%).

**The acute inflammation signaling** appears to be activated by upregulating the E2F1 target protein TRAF6 (115.6%) and coincidentally upregulating MIF (112.5%) and CXCR4 (111.8%), despite the downregulation of MCP1 (90.2%) and CRP (89.9%).

**The cellular immunity signaling** appears to be activated by upregulating the E2F1 target proteins CD8 (144.7%), and CD99 (112.3%) and coincidentally upregulating CD28 (116%), CD34 (127.7%), IL28 (110.3%), and granzyme B (133.7%) and downregulating CTLA4 (91.7%), despite the downregulation of the E2F1 target protein CD4 (94.6%) and upregulation of PDL1 (118%), as well as the coincidental downregulation of CD3 (93.4%), versican (90.5%), and perforin (69.6%).

**The chronic inflammation signaling** was activated by upregulating the E2F1 target protein cathepsin C (114.2%) and coincidentally upregulating IL12 (106.8%), lysozyme (107.5%), MMP10 (119.1%), MMP12 (110.9%), TIMP1 (116.5%), TIMP2 (146.5%), cathepsin G (120.2%), cathepsin K (114.3%), LTA4H (122.5%), and kininogen (128.2%), despite the downregulation of the E2F1 target proteins CD68 (83.8%) and MMP2 (93.5%), and coincidental downregulation of IL10 (86.7%), CD31 (91.6%), MMP3 (79.5%), and COX1 (68.6%).

**The p53-mediated apoptosis signaling** appears to be activated by upregulating the E2F1 target proteins p53 (109.4%), BAD (124.2%), BAK (105.4%), PUMA (106.8%), NOXA (115.6%), and BAX (106.7%) and coincidentally downregulating MDM2 (89.5%), despite the downregulation of the E2F1 target proteins p73 (88.2%) and caspase3 (93.8%), and upregulation of BCL2 (109.7%), as well as coincidental downregulation of p63 (90.6%).

**The senescence signaling** was activated by upregulating the E2F1 target proteins klotho (115.8%), sirtuin7 (109.2%), and p21 (114.8%) and coincidentally upregulating sirtuin3 (110.7%), despite the downregulation of the E2F1 target proteins sirtuin1 (91.9%) and sirtuin6 (84.4%).

**The telomere signaling** was activated by upregulating the E2F1 target protein TERT (118.1%) and coincidentally upregulating TIN2 (133.4%), despite the downregulation of the E2F1 target protein TRF1 (86.9%).

In addition, the E2Fs binding site sequence*-CEDS significantly influenced **the epigenetic modification signaling** by downregulating the E2F1 target protein HMGA2 (93.8%) and coincidentally downregulating HDAC10 (89.8%), while it upregulated the E2F1 target proteins KDM4D (115%), DNMT1 (107.8%), BRG1 (111.4%), and lamin A/C (108.8%) and coincidentally upregulated Ac-lysine (113.7%). Consequently, the TTTC(C/G)CGC CEDS induced a trend towards an increase in the methylation of histones and DNAs, and transcriptional repression.

The E2Fs binding site sequence*-CEDS significantly suppressed **the oncogenesis signaling** by downregulating the E2F1 target proteins BRCA1 (93.9%), PIM1 (93.7%), BRCA2 (93.4%), ZEB1 (91.9%), cMyb (90.5%), CRK (89.2%), TWIST1 (87.1%), NF1 (87%), CRIP1 (84.3%), and Daxx (72.4%) and coincidentally downregulating MTA2 (89.4%), BMI1 (87.8%), and BACH1 (84.8%). while it also enhanced **the oncogenesis signaling** by upregulating the E2F1 target proteins MALT1 (112.6%), YAP1 (106.4%), and XBP1 (105.8%) and coincidentally upregulating DMBT1 (122.1%), 14-3-3 (115.2%), CEA (112.1%), EWSR1 (109.9%), MTDH (106.5%), and HER2 (105.8%). Among the 28 oncoproteins, 14 were found to be underexpressed, nine were overexpressed, and five exhibited minimal change in expression compared to the untreated controls.

Consequently, the canonical E2Fs binding site sequence (TTTC(C/G)CGC)*-CEDS strongly impact the protein signaling in RAW 264.7 cells. The TTTC(C/G)CGC*-CEDS primary enhanced the RAS-NFkB signaling axis, subsequently activated proliferation-related signaling axis, epigenetic modification-protection, cytodifferentiation-angiogenesis-acute inflammation signaling axis, senescence-cellular immunity-chronic inflammation signaling axis, and epigenetic modification-p53-mediated apoptosis-telomere signaling axis, while it inactivated the ER stress-survival-innate immunity signaling axis and the FAS- and PARP-mediated apoptosis-fibrosis-glycolysis-oncogenesis signaling axis (Fig. 22B, Table 16).

**Table 16.** Protein expressions in the RAW 264.7 cells treated with canonical E2Fs binding site sequence (TTTC(C/G)CGC)*-CEDS.

| Signaling | Downregulated Proteins | Upregulated Proteins |
| --- | --- | --- |
| **Proliferation** | PCNA, PLK4, p14, p27, nucleolin | Ki-67, CDK4, cyclin D2, p21, CHK1 |
| **cMyc/MAX/MAD network** | MAD1, p27 | cMyc, MAX |
| **Wnt/β-catenin signaling** | Wnt1, β-catenin, AXIN2 | APC |
| **p53/Rb/E2F1 signaling** | p-Rb1, E2F4 | p53, CDK4, E2F1 |
| **SHH/PTCH/GLI signaling** | GLI1, | SHH |
| **Notch/Jagged signaling** |  | Notch1, Jagged2 |
| **Epigenetic modification** | HDAC10, HMG A2 | KDM4D, DNMT1, BRG1, Ac-lysine, lamin A/C |
| **Protein translation** | DOHH, eIF5A1, eIF5A2, eIF2α | DHPS, eIF2AK3 |
| **miRNA biogenesis** | Exportin5, Dicer, GW182 | TRBP2, MTDH |
| **Growth factors** | IGF2R, TGFβ3, ALK1, SMAD4, SMAD7, CTGF | GH, GHRH, EGF, TGFβ1, TGFβ2, SMAD2/3, HGFα, FGF1, FGF2, |
| **Cytodifferentiation** | β-actin, GAPDH, CRIP1, VE-cadherin | α-tubulin, FAK, CaM, Ep-CAM, cystatin A, claudin1, DLX2, SOX9, tenascin C |
| **Osteogenesis** | BMP2, BMP3, OPG, RANKL | RUNX2, PTH/PTHrPR, osteocalcin, aggrecan, SOSTDC1 |
| **Neuromuscular differentiation** | Desmin | GFAP, NK1R, S100, MYH2 |
| **RAS signaling** | KRAS, PTEN, STAT3, p-STAT3, c-Fos, p-JNK1, p-TYK2 | HRAS, PI3K, Rab1, PKA1α, PLCβ2, JNK1, PKC< p-PKC1α, SOS1 |
| **NFκB signaling** | p-p38, NFATc1 | MDR, GADD45, p38, mTOR, NFAT5 |
| **p53-mediated apoptosis** | p63, p73, MDM2, caspase3 | p53, BAD, BAK, PUMA, NOXA, BAX, BCL2 |
| **FAS-mediated apoptosis** | FASL, FAS, TRAIL, BID, caspase8, caspase3 |  |
| **PARP-mediated apoptosis** | PARP1 |  |
| **Protection** | HO1, FXP4, FOXO1, HSP90, NOS1 | HSP27, hepcidin, HSP70, ALDH1A1, GSTO1/2, SOD1, SVCT2 |
| **ER stress** | eIF2α, p-GADD153, BIP, IRE1α, ATF4 | HSP27, eIF2AK3, ATF6α |
| **Angiogenesis** | VEGFC, endocan, CD31, angiogenin | HIF1α, VEGFR2, vWF, FLT4, LYVE1, angiogenin, PDGFA, FGF2 |
| **Survival** | sirtuin1, sirtuin6, survivin | sirtuin3, sirtuin7 |
| **Acute inflammation** | MCPI, CRP | TRAF6, MIF, CXCR4 |
| **Innate immunity** | Lactoferrin, LL37, β-defensin2, β-defensin3, TLR3, α1-antitrypsin | β-defensin1, TLR4, elafin |
| **Cellular immunity** | CD3, CD4, versican, perforin, CTLA4 | CD8, CD20, CD28, CD34, CD99, IL28, granzyme B, PDL1 |
| **Chronic inflammation** | IL10, CD31, CD68, MMP2, MMP3, COX1 | IL12, lysozyme, MMP10, MMP12, TIMP1, TIMP2, cathepsin C, cathepsin G, cathepsin K, LTA4H, kininogen |
| **Glycolysis** | GLUT1, TIGAR, GAPDH, LGR4, GLUT4 | HK2, PPARγ, |
| **Fibrosis** | collagen 3A1, CTGF, integrin α5, integrin β1, α1-antitrypsin | FGF1, FGF2, tenascin C, claudin1 |
| **Oncogenesis** | BRCA1, PIM1, BRCA2, ZEB1, cMyb, MTA2, CRK, BMI1, TWIST1, NF1, BACH1, CRIP1, Daxx | DMBT1, 14-3-3, MALTI1, CEA, EWSR1, MTDH, YAP1, HER2, XBP1 |
| **Senescence** | sirtuin1, sirtuin6 | klotho, sirtuin3, sirtuin7, p21 |
| **Telomere signaling** | TRF1 | TERT, TIN2 |

The canonical E2Fs binding site sequence\*-CEDS, TTTC(C/G)CGC\*-CEDS induced the suppression (blue) and activation (red) of signaling by downregulating (blue) and upregulating (red) the proteins, respectively. The data were compared to the reports of recent manuscripts and Harmonizome 3.0 website.

The results indicate that TTTC(C/G)CGC*-CEDS had a strong potential of oncogenic effect on RAW 264.7 by activating proliferation, ROS protection, angiogenesis, senescence, chronic inflammation, and telomere instability, while it had a weak potential of anti-oncogenic effect by attenuating ER stress, survival, fibrosis, glycolysis, and oncogenesis. It was also found that the TTTC(C/G)CGC*-CEDS can induce acute and chronic inflammation but diminish the FAS- and PARP-mediated apoptosis.

#### The poly-A sequence (12A)*-CEDS influenced the protein signaling in RAW 264.7 cells

The poly-A sequences (4A-12A) are ubiquitously distributed throughout the mouse genome. Therefore, the poly-A sequence may play a role in gene transcription in different regions of the genome, but its function will be variable and nonspecific depending on the context of the genomic environment. In the present study, the poly-A sequence (12A) was selected for CEDS experiment as a nonspecific control to compare with other CEDSs using different miRNA and DNA motif sequences that have been shown to be essential for gene expression.

RAW 264.7 cells treated with the 12A*-CEDS exhibited changes in protein expression throughout essential signaling, both positively and negatively. However, each protein signaling exhibited unique trends of activation or inactivation. As a result, the 12A*-CEDS increased protein signaling related to proliferation, cMyc/MAX/MAD network, p53/Rb/E2F1 signaling, SHH/PTCH/GLI signaling, Notch/Jagged signaling, miRNA biogenesis, growth factors, cytodifferentiation, RAS signaling, ER stress, angiogenesis, survival, acute inflammation, PARP-mediated apoptosis, and fibrosis. On the other hand, it activated protein signaling related to Wnt/β-catenin signaling, epigenetic modification, protein translation, osteogenesis, neuromuscular differentiation, NFkB signaling, protection, innate immunity, cellular immunity, chronic inflammation, p53- mediated apoptosis, FAS-mediated apoptosis, oncogenesis, senescence, glycolysis, and telomere signaling (Fig. 23A).

**Fig. 23.**
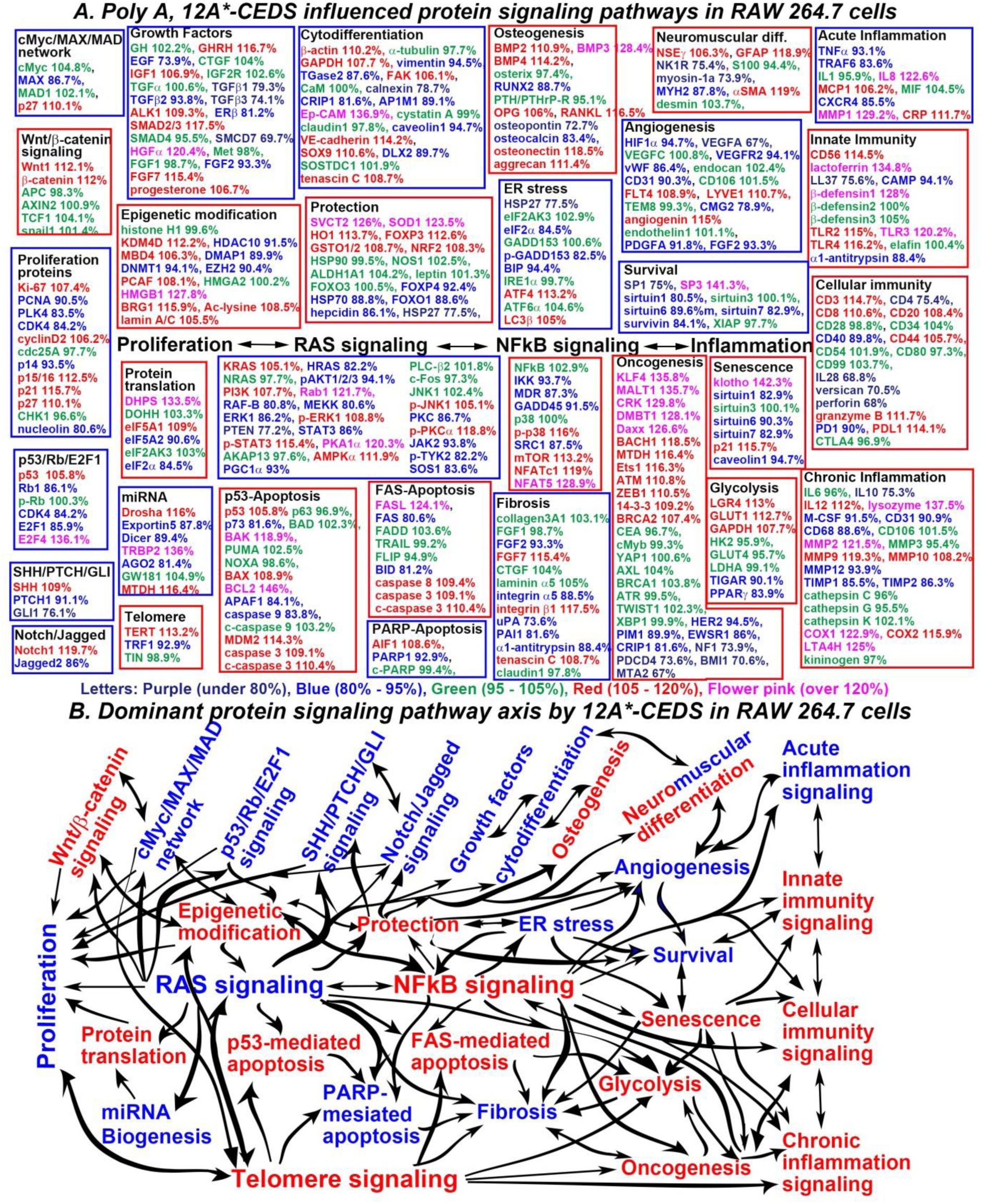
The nonspecific sequence (12A)*-CEDS influenced the protein signaling (A) and axes (B) in RAW 264.7 cells. The IP-HPLC results revealed proteins that were downregulated 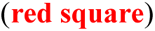, upregulated 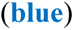, and minimally changed 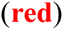 compared to the untreated controls. This allowed the dominant trend of suppressed 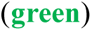 and activated 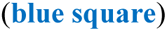 signaling to be defined.

In RAW264.7 cells treated with the 12A*-CEDS, **the proliferation signaling** appears to be inactivated by downregulating PCNA (up to 90.5%), PLK4 (83.5%), CDK4 (84.2%), and nucleolin (80.6%), and upregulating p15/16 (112.5%), p21 (115.7%), and p27 (110.1%), despite the upregulation of Ki-67 (107.4%) and cyclinD2 (106.2%).

**The cMyc/MAX/MAD network** appears to be inactivated by downregulating MAX (86.7%), and upregulating p27 (110.1%).

**The p53/Rb/E2F1 signaling** was inactivated by downregulating Rb1 (86.1%), CDK4 (84.2%), and E2F1 (85.9%), as well as coincidentally upregulating p53 (105.8%) and E2F4 (136.1%).

**The SHH/PTCH/GLI signaling** was inactivated by downregulating GLI1 (76.1%) despite the slight upregulation of SHH (109%).

**The Notch/Jagged signaling** appears to be inactivated by downregulating Jagged2 (86%) despite the upregulation of Notch1 (119.7%).

**The growth factor signaling** were partially inactivated by downregulating EGF (73.9%), TGFβ1 (79.3%), TGFβ2 (93.8%), TGFβ3 (74.1%), ERβ (81.2%), SMCD7 (69.7%), and FGF2 (93.3%), despite the upregulation of GHRH (116.7%), IGF1 (106.9%),, ALK1 (109.3%), SMAD2/3 (117.5%), HGFα (120.4%), FGF7 (115.4%), and progesterone (106.7%).

**The cytodifferentiation signaling** were partially inactivated by downregulating vimentin (94.5%), TGase2 (87.6%), calnexin (78.7%), CRIP1 (81.6%), AP1M1 (89.1%), caveolin1 (94.7%), and DLX2 (89.7%), despite the upregulation of β-actin (110.2%), GAPDH (107.7 %), FAK (106.1%), Ep-CAM (136.9%), VE-cadherin (114.2%), SOX9 (110.6%), and tenascin C (108.7%).

**The RAS signaling** appears to be inactivated by downregulating HRAS (82.2%), pAKT1/2/3 (94.1%), RAF-B (80.8%), MEKK (80.6%), ERK1 (86.2%), PTEN (77.2%), STAT3 (86%), PGC1α (93%), PKC (86. 7%), JAK2 (93.8%), p-TYK2 (82.2%), and SOS1 (83.6%), despite the upregulation of KRAS (105.1%), PI3K (107.7%), Rab1 (121.7%), p-ERK1 (108.8%), p-STAT3 (115.4%), PKA1α (120.3%), AMPKα (111.9%), p-JNK1 (105.1%), and p-PKCα (118.8%).

**The ER stress signaling** was inactivated by downregulating HSP27 (77.5%), eIF2α (84.5%), and p- GADD153 (82.5%), and BIP (94.4%), despite the upregulation of ATF4 (113.2%) and LC3β (105%).

**The angiogenesis signaling** was suppressed by downregulating HIF1α (94.7%), VEGFA (67%), vWF (86.4%), CD31 (90.3%), CMG2 (78.9%), PDGFA (91.8%), and FGF2 (03.3%), despite the upregulation of FLT4 108.9%), LYVE1 (110.7%), and angiogenin (115%).

**The survival signaling** was suppressed by downregulating SP1 (75%), sirtuin1 (80.5%), sirtuin6 (89.6%), sirtuin7 (82.9%), and survivin (84.1%), despite the upregulation of SP3 (141.3%).

**The acute inflammation signaling** appears to be inactivated by downregulating TNFα (93.1%), TRAF6 (83.6%), and CXCR4 (85.5%), despite the upregulation of IL8 (122.6%), MCP1 (106.2%), MMP1 (129.2%), and CRP (111.7%).

**The PARP-mediated apoptosis signaling** was inactivated by downregulating PARP1 (92.9%), despite the upregulation of AIF1 (108.6%).

**The fibrosis signaling** was inactivated by downregulating FGF2 (93.3%), integrin α5 (88.5%), uPA (73.6%), PAI1 (81.6%), and α1-antitrypsin (88.4%), despite the upregulation of FGF7 (115.4%), integrin β1 (117.5%), and tenascin C (108.7%).

Conversely, **the Wnt/β-catenin signaling** appears to be activated by upregulating Wnt1 (112.1%) and β- catenin (112%).

**The protein translation signaling** appears to be activated by upregulating DHPS (133.5%) and eIF5A1 (109%), despite the downregulation of eIF5A2 (90.6%) and eIF2α (84.5%).

**The osteogenesis signaling** was activated by upregulating BMP2 (110.9%), BMP3 (128.4%), BMP4 (114.2%), OPG (106%), RANKL (116.5%), osteonectin (118.5%), and aggrecan (111.4%), despite the downregulation of RUNX2 (88.7%), osteopontin (72.7%), and osteocalcin (83.4%).

**The neuronal differentiation signaling** was activated by upregulating NSEγ (106.3%), GFAP (118.9%), despite the downregulation of NK1R (75.4%), while cellular differentiation signaling appeared to be inactivated by downregulating myosin-1a (73.9%) and MYH2 (87.8%), despite the upregulation of αSMA (119%).

**The NFkB signaling** appears to be activated by upregulating p-p38 (116%), mTOR (113.2%), NFATc1 (119%), and NFAT5 (128.9%) and coincidentally upregulating IKK (93.7%), despite the downregulation of MDR (87.3%), GADD45 (91.5%), and SRC1 (87.5%).

**The protection signaling** appears to be inactivated by upregulating SVCT2 (126%), SOD1 (123.5%), HO1 (113.7%), FOXP3 (112.8%), GSTO1/2 (108.7%), and NRF2 (108.3%), despite the downregulation of FOXP4 (92.4%), HSP70 (88.8%), FOXO1 (88.6%), hepcidin (86.1%), and HSP27 (77.5%).

**The innate immunity signaling** was activated by upregulating CD56 (114.5%), lactoferrin (134.8%), β- defensin1 (128%), TLR2 (115%), TLR3 (120.2%), and TLR4 (116.2%), despite the downregulation of LL37 (75.6%), CAMP (94.1%), and α1-antitrypsin (88.4%).

**The celluar immunity signaling** appears to be activated by upregulating CD3 (up to 114.7%), CD8 (110.6%), CD20 (108.4%), CD44 (105.7%), granzyme B (111.7%), and PDL1 (114.1%), despite the downregulation of CD40 (89.8%), CD40 (89.8%), IL28 (68.8%), versican (70.5%), perforin (68%), and PD1 (90%).

**The chronic inflammation signaling** appear to be activated by upregulating IL12 (112%), lysozyme (137.5%), MMP2 (121.5%), MMP9 (119.3%), MMP10 (108.2%), COX1 (122.9%), COX2 (115.9%), and LTA4H (125%), despite the downregulation of IL10 (75.3%), M-CSF (91.5%), CD31 (90.9%), CD68 (88.6%), MMP12 (93.9%), TIMP1 (85.5%), and TIMP2 (86.3%).

**The p53-mediated apoptosis signaling** was activated by upregulating p53 (105.8%), BAK (118.9%), BAX (108.9%), BCL2 (146%), MDM2 (114.3%), caspase3 (109.1%), and c-caspase3 (110.4%), despite the downregulation of p73 (81.6%), APAF1 (84.1%), and caspase9 (83.8%).

**The FAS-mediated apoptosis signaling** appears to be activated by upregulating FASL (124.1%), caspase8 (109.4%), caspase3 (109.1%), and c-caspase3 (110.4%), despite the downregulation of FAS (80.6%) and BID (81.2%).

**The senescence signaling** appears to be activated by upregulating klotho (142.3%) and p21 (115.7%), despite the downregulation of sirtuin1 (82.9%), sirtuin6 (90.3%), sirtuin7 (82.9%), and caveolin1 (94.7%).

**The glycolysis signaling** appears to be activated by upregulating LGR4 (113%), GLUT1 (112.7%), and GAPDH (107.7%), despite the downregulation of TIGAR (90.1%) and PPARγ (83.9%).

**The telomere signaling** was activated by upregulating TERT (up to 113.2%), and downregulating TRF1 (92.9%).

In addition, the 12A*-CEDS markedly influenced **the epigenetic modification signaling** by downregulating HDAC10 (91.5%), DMAP1 (89.9%), and DNMT1 (94.1%), as well as coincidentally upregulating KDM4D (112.2%), MBD4 (106.3%), PCAF (108.1%), HMGB1 (127.8%), BRG1 (115.9%), Ac-lysine (108.5%), and lamin A/C (105.5%). Consequently, the 12A*-CEDS induced a trend towards an increase in the methylation of histones and DNAs, and transcriptional repression.

The 12A*-CEDS significantly suppressed **the oncogenesis signaling** by downregulating HER2 (94.5%), PIM1 (89.9%), EWSR1 (86%), CRIP1 (81.6%), NF1 (73.9%), PDCD4 (73.6%), BMI1 (70.6%), and MTA2 (67%), while it widely activating **the oncogenesis signaling** by upregulating KLF4 (135.8%), MALT1 (135.7%), CRK (129.8%), DMBT1 (128.1%), Daxx (126.6%), BACH1 (118.5%), MTDH (116.4%), Ets1 (116.3%), ATM (110.8%), ZEB1 (110.5%), 14-3-3 (109.2%), and BRCA2 (107.4%). Among the 28 oncoproteins, 12 were found to be overexpressed, eight were overexpressed, and eight exhibited minimal change in expression compared to the untreated controls.

Consequently, the nonspecific sequence (12A)*-CEDS significantly impacted the protein signaling in RAW 264.7 cells. The 12*-CEDS suppressed the RAS signaling, subsequently inactivating proliferation-related signaling axis, ER stress-angiogenesis-survival-acute inflammation signaling axis, and PARP-mediated apoptosis-fibrosis signaling axis. In contrast, it enhanced the NFkB signaling, subsequently activating the p53- and FAS-mediated apoptosis signaling axis, and senescence-glycolysis-oncogenesis-innate immunity-chronic inflammation signaling axis (Fig. 23B, Table 17).

**Table 17.** Protein expressions in the RAW 264.7 cells treated with nonspecific sequence (12A)*-CEDS.

| Signaling | Downregulated Proteins | Upregulated Proteins |
| --- | --- | --- |
| Proliferation | PCNA, PLK4, CDK4, p14, nucleolin | Ki-67, cyclin D2, p15/16, p21, p27 |
| cMyc/MAX/MAD network | MAX | p27 |
| Wnt/β-catenin signaling | | Wnt1, β-catenin |
| p53/Rb/E2F1 signaling | Rb1, CDK4, E2F1 | p53, E2F4 |
| SHH/PTCH/GLI signaling | PTCH1, GLI1 | SHH |
| Notch/Jagged signaling | Jagged2 | Notch1 |
| Epigenetic modification | HDAC10, DMAP1, DNMT1, EZH2 | KDM4D, MBD4, PCAF, HMGB1, BRG1, Ac-lysine, lamin A/C |
| Protein translation | eIF5A2, eIF2α | DHPS, eIF5A1 |
| miRNA biogenesis | Exportin5, Dicer, AGO2, | Drosha, TRBP, MTDH |
| Growth factors | EGF, TGFβ1, TGFβ2, TGFβ3, SMCD7, FGF2, ERβ | GHRH, NF1, ALK1, SMCD2/3, HGFα, FGF7, progesterone |
| Cytodifferentiation | Vimentin, TGase2, calnexin, CRIP1, AP1M1, caveolin1, DLX2 | β-actin, GAPDH, FAK, Ep-CAM, VE-cadherin, SOX9, tenascin C |
| Osteogenesis | RUNX2, osteopontin, osteocalcin, | BMP2, BMP3, BMP4, OPG, RANKL, osteonectin, aggrecan |
| Neuromuscular differentiation | NK1R, myosin-1a, MYH2 | NSEγ, GFAP, αSMA |
| RAS signaling | HRAS, pAKT1/2/3, RAF-B, MEKK, ERK1, PTEN, STAT3, PGC1α, PKC, JAK2, p-TYK2, SOS1 | KRAS, PI3K, Rab1, p-STAT3, PKA1α, AMPKα, p-JNK1, p-PKC1α |
| NFκB signaling | IKK, MDR, GADD45, SRC1 | p-p38, mTOR, NFATc1, NFAT5 |
| p53-mediated apoptosis | p73, APAF1, caspase9 | p53, BAK, BAX, BCL2, MDM2, caspase3, c-caspase3 |
| FAS-mediated apoptosis | FAS, BID | FASL, caspase8, caspase3, c-caspase3 |
| PARP-mediated apoptosis | PARP1 | AIF1 |
| Protection | FOXP4, HSP70, FOXO1, hepcidin, HSP27 | SVCT2, SOD1, HO1, FOXP3, GSTO1/2, NRF2, |
| ER stress | HSP27, eIF2α, p-GADD153, BIP | ATF4, LC3β |
| Angiogenesis | HIF1α, VEGFA, VEGFR2, vWF, CD31, CMG2, PDGFA, FGF2 | FLT4, LYVE1, angiogenin |
| Survival | SP1, sirtuin1, sirtuin1, sirtuin6, sirtuin7, survivin | SP3 |
| Acute inflammation | TNFα, TRAF6, CXCR4 | IL8, MCP1, MMP1, CRP |
| Innate immunity | LL37, CAMP, α1-antitrypsin | CD56, lactoferrin, TLR2, TLR3, TLR4 |
| Cellular immunity | CD4, CD40, IL28, versican, perforin, PD1 | CD3, CD8, CD20, CD44, granzyme B, PDL1 |
| Chronic inflammation | IL6, IL10, M-CSF, CD31, CD68, MMP12, TIMP1, TIMP2 | IL12, lysozyme, MMP2, MMP9, MMP10, COX1, COX2, LTA4H |
| Glycolysis | TIGAR, PPARγ | LGR4, GLUT1, GAPDH |
| Fibrosis | FGF2, integrin α5, uPA, PAH1, α1-antitrypsin | FGF7, integrin β1, tenascin C |
| Oncogenesis | HER2, PIM1, EWSR1, CRIPI, NF1, PDCD4, BMI1, MTA2 | KLF4, MALT1, CRK, DMBT1, Daxx, BACH1, MTDH, Ets1, ATM, ZEB1, 14-3-3, BRCA2 |
| Senescence | sirtuin1, sirtuin6, sirtuin7, caveolin1 | klotho, p21 |
| Telomere signaling | TRF1 | TERT |

The nonspecific sequence\*-CEDS, 12A\*-CEDS induced the suppression (blue) and activation (red) of signaling by downregulating (blue) and upregulating (red) the proteins, respectively.

The results indicate that the 12*-CEDS also affects the protein signaling in a similar manner to the above experiments, due to the existence of numerous poly-A sequences (4A-12A) in the mouse genome. The 12A*- CEDS resulted in imbalanced protein signaling in RAW 264.7 cells, activating epigenetic methylation, oncogenesis, and telomere instability, leading to chronic inflammation and apoptosis, in the absence of cellular growth and differentiation, wound healing, and ROS protection. Therefore, it is postulated that the 12A*-CEDS may influence the cells to undergo aging and retrogressive changes with a high potential of oncogenesis.

The IP-HPLC analysis utilizing 350 antisera demonstrated that CEDS, which employ different DNA motif sequences, can influence the protein expression of global protein signaling in RAW 264.7 cells. The p53 (cluster 1 response element) binding site sequence (AGACATGCCT)*-CEDS upregulated 61 proteins (32.3%), downregulated 70 proteins (37%), and exhibited minimum effect on 58 proteins (30.1%) among 189 p53 targeting proteins. And the canonical E2Fs binding site sequence (TTTC(C/G)CGC)*-CEDS upregulated 69 proteins (41.3%), downregulated 67 proteins (40.1%), and exhibited minimum effect on 31 proteins (18.6%) among 167 E2F1 targeting proteins observed in this study (Table 19).

**Table 19.** The targeting efficiency of CEDS using a different DNA motif sequence in RAW 264.7 cellss.

| **Protein binding site sequence\*-CEDS** | **Up-regulation (over 105%)** | **Down-regulation (under 95%)** | **Minimal change ($\pm 5\%$)** | **Total target proteins** |
| --- | --- | --- | --- | --- |
| **AGACATGCCT\*-CEDS (p53 binding site sequence)** | 61 | 70 | 58 | 189 |
|  | 32.3 | 37% | 30.1% | 100% |
| **TTTC(C/G)CGC\*-CEDS (canonical E2Fs binding site sequence)** | 69 | 67 | 31 | 167 |
|  | 41.3% | 40.1% | 18.6% | 100% |
| **Total** | 232 | 268 | 125 | 625 |
|  | 37.1% | 42.9% | 20% | 100% |
| **12A\*-CEDS (nonspecific sequence)** | 130 | 122 | 98 | 350 |
|  | 37.1% | 34.9% | 28% | 100% |

Consequently, the p53 (cluster 1 response element) binding site sequence (AGACATGCCT), and the canonical E2Fs binding site sequence (TTTC(C/G)CGC) demonstrated a relatively similar ratio of targeting efficiency between the upregulation versus downregulation of their targeting proteins, with 41.8% versus 40%, 32.3% versus 37%, and 41.3% versus 40.1%, respectively.

### Cytological and immunocytochemical observation of CEDS-treated cells

#### Cytological observation of RAW 264.7 cells treated with miRNA sequence*-CEDS

RAW 264.7 cells were cultured on cell culture slides (Hybridwell™, SPL Life Science, Korea) using the same method as the above experiment. Approximately 50% confluent RAW 264.7 cells grown on culture slide surfaces were treated with murine mature miRNA sequence*-CEDS at 23-25 Gauss for 20 min and once more in 12 hours. After a culture period of 24 hours, the cells were fixed with 10% buffered formalin, stained with hematoxylin and eosin, and also underwent immunocytochemical staining for c-caspase 3.

Immunocytochemical (ICC) staining was performed using the indirect triple sandwich method on the Vectastatin system (Vector Laboratories, USA), and visualized using a 3-amino-9-ethylcarbazole solution (Santa Cruz Biotechnology, USA) with no counter staining. The results were observed by optical microscope, and their characteristic images were captured (DP-73, Olympus Co., Japan) and illustrated.

The control group which was not treated with CEDS showed many cells diffusely scattered on culture dish surface, and the cells were occasionally positive for c-caspase 3. Whereas CEDS using mmu-miR-21-5p sequence TAGCTTATCAGACTGATGTTGA*-CEDS, showed proliferative cells which were frequently aggregated into multicellular spheroids. The cells were weakly positive for c-caspase 3. CEDS using mmu-miR- 26a-5p sequence, TTCAAGTAATCCAGGATAGGCT*-CEDS showed marked decrease of cell number, and strong immunoreaction of c-caspase 3 in the remaining cells. CEDS using mmu-miR-29a-3p sequence, TAGCACCATCTGAAATCGGTTA*-CEDS showed relatively abundant cells which were partly aggregated. Some cells were positive for c-caspase 3 (Fig. 24 A-P).

**Fig. 24.**
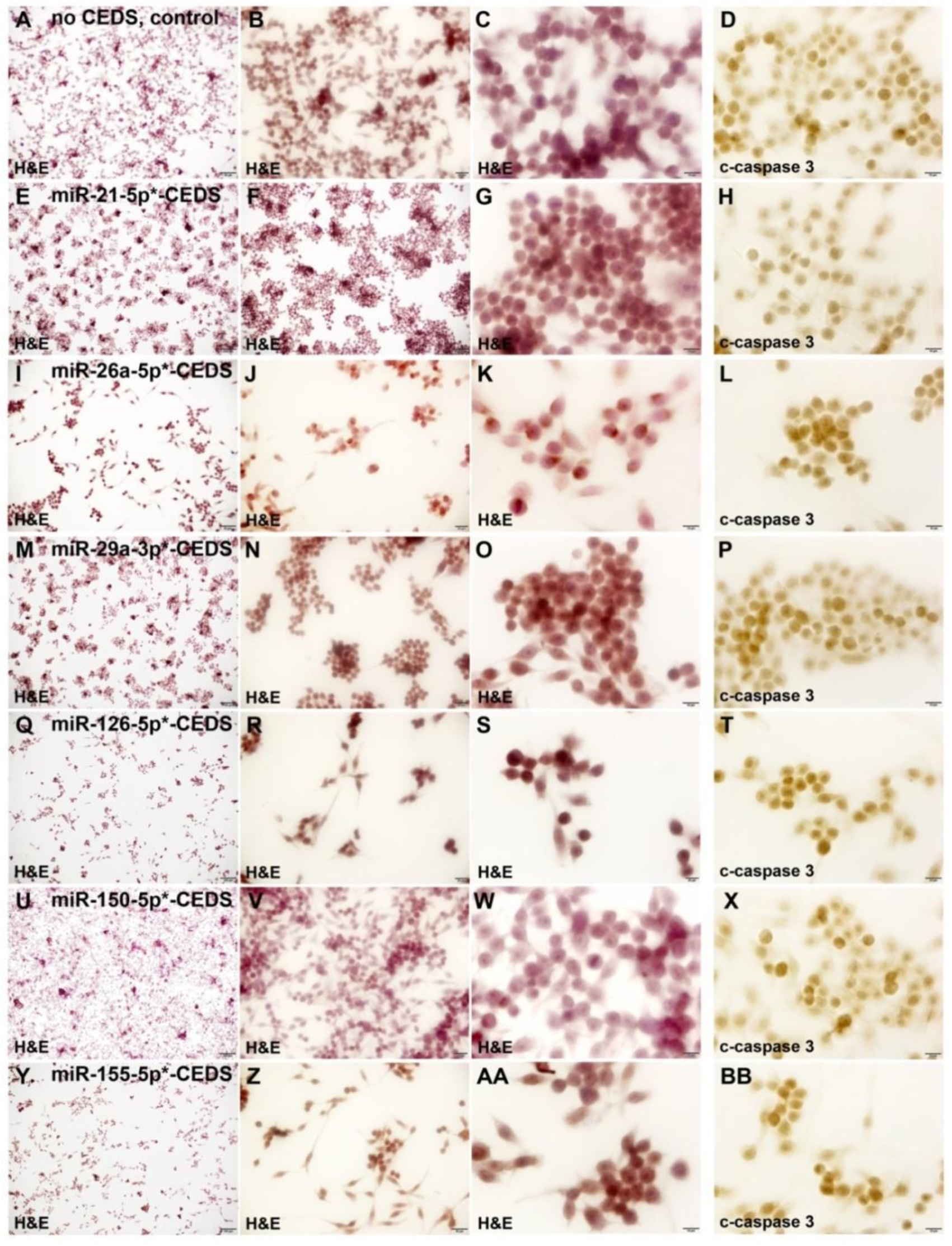
Cytological and immunocytochemical staining on RAW 264.7 cells treated with CEDS using murine mature miRNA sequence. A-D: Control, no CEDS, diffuse spread of cells (A-C), and occasional for c-caspase 3 (D). E-H: MiR-21-5p*-CEDS, marked aggregation of cells (E-G), and weak for c-caspase 3(H). I-L: MiR-26a- 5p*-CEDS, marked decrease of cell number (I-K), and strong for c-caspase 3(L). M-P: MiR-29a-3p*-CEDS, slight aggregation of cells (M-O), and occasional for c-caspase 3 (P). Q-T: MiR-126-5p*-CEDS, marked decrease of cell number (Q-S), and strong for c-caspase 3 (T). U-X: MiR-150-5p*-CEDS, slight decrease of cell number (U-W), and slight for c-caspase 3 (X). Y-BB: MiR-155-5p*-CEDS, significant decrease of cell number (Y-AA), moderate for c-caspase 3 (BB).

CEDS using mmu-miR-126-5p sequence, TTCAAGTAATCCAGGATAGGCT*-CEDS showed marked decrease of cells, and strong immunoreaction of c-caspase 3 in the remaining cells. CEDS using mmu-miR-150- 5p sequence, TCTCCCAACCCTTGTACCAGTG*-CEDS showed slight decrease of cell number, and positive immunoreaction of c-caspase 3 in some cells. CEDS using mmu-miR-155-5p sequence, TTCAAGTAATCCAGGATAGGCT*-CEDS showed significant decrease of cell number, and showed strong immunoreaction of c-caspase 3 (Fig. 24 Q-BB)

The cytological results demonstrated that miR-26a-5p or miR-126-5p*-CEDS induced a strong anti-proliferative and apoptotic effect on RAW 264.7 cells compared to the untreated controls. The anti-proliferative and apoptotic effect was found to be diminished in CEDS using miR-155-5p, miR-150-5p, and miR-29a-3p sequence in order, while CEDS using miR-21-5p sequence was observed to induce the proliferation and aggregation of cells, thereby exerting an oncogenic effect.

The DNA and RNA are the central reservoir in cells, which can modulate cellular structures and functions through DNA/RNA transcription and protein translation. It is thought that they have special biological signals to function as distinctive modulators. The nucleotides themselves are amphoteric, but become acidic when the cryptic internalization of base pairs occurs after hybridization. The acidity of hybridized nucleotide complexes, including dsDNA and dsRNA, may be relevant to the strength of the hybridization. Therefore, it can be postulated that the slight acidity of dsDNA and dsRNA may exert the intramolecular electric conductivity, which is controllable by the level of hydrogen bonding strength between base pairs.

In another sense, the dsDNA and dsRNA may exhibit semiconducting properties that are dependent on changes in the hydrogen bonding strength between base pairs. It is therefore reasonable to postulate that the unique semi-conductive properties of dsDNA and dsRNA will enable them to carry out the full range of biological signaling processes associated with DNA and RNA transcription and protein translation, and that it could be readily affected by CEDS in a sequence-specific manner.

CEDS utilized in this study was designed to regulate the hydrogen bonding strength between base pairs of dsDNA and dsRNA. CEDS is available due to the regular cyclic arrangement of base pairs in dsDNA and dsRNA. It can target a specific sequence of dsDNA and dsRNA by CEDS due to the same angular momentum between macroscale and ultramicroscale structures of dsDNA and dsRNA. The preceding study demonstrated that the targeting efficiency of CEDS to randomly arranged dsRNA or dsRNA is approximately 25%.

The objective of this study is to identify the cellular changes that occur following CEDS targeting nine mature miRNAs and four protein binding site sequences in RAW 264.7 cells. The gross and cytological observations were initially conducted, and subsequently, the protein expression changes that influence the entire protein signaling were examined through IP-HPLC using monospecific 350 antisera.

It was hypothesized that CEDS could enhance the hydrogen bonding strength between DNA/RNA base pairs and induce the native base pair polarities of DNA/RNA, which are required for DNA/RNA transcription. The present study investigated the changes in the expression of mature and primary miRNAs through the real-time qPCR method. The qPCR results demonstrated that both mature and primary miRNAs were increased in number after CEDS. It appears that the mature miRNAs exhibited a greater increase than the primary miRNAs in response to CEDS treatment. These findings suggest that CEDS may facilitate not only the production of mature miRNAs from primary miRNAs but also the formation of primary miRNAs from miRNA gene transcripts.

The nine miRNA sequence*-CEDSs investigated in this study demonstrated a significant impact on the expression of target proteins, with an average reduction of 66.4%. MiR-150-5p*-CEDSs exhibited a potent anti-oncogenic effect, accompanied by a strong anti-inflammatory effect, but a reduced potential for apoptosis, innate and cellular immunity in RAW 264.7 cells. In contrast, miR-34a-5p, miR-181a-5p, miR-216a-5p, miR- 365a-3p, and miR-655-3p*-CEDSs exhibited a dual effect, inducing both anti-oncogenic and oncogenic effects. MiR-155-5p*-CEDS demonstrated a tendency to induce a marked oncogenic effect, with an elevated potential of proliferation, survival, senescence, and fibrosis, rather than an anti-oncogenic effect on RAW 264.7 cells.

Furthermore, it has been demonstrated that CEDS using a single miRNA sequence can influence the expression of other miRNAs that are linked to it through a feedback loop involving gene transcription and protein translation in qPCR and IP-HPLC analysis. For instance, miR-26a-5p*-CEDS was observed to increase the expression of miR-126-5p and miR-655-3p while concomitantly decreasing the expression of miR-365a-3p. Similarly, miR-34a-5p*-CEDS was found to elevate the expression of miR-126-5p and miR-655-3p, while concurrently reducing the expression of miR-21-5p. MiR-126-5p*-CEDS increased the expression of miR-26a- 5p and miR-655-3p. MiR-150-5p*-CEDS increased the expression of miR-126-5p, miR-365a-3p, and miR-655- 3p. MiR-155-5p*-CEDS increased the expression of miR-181a-5p. MiR-181a-5p*-CEDS increased the expression of miR-126-5p, miR-216a-5p, and miR-655-3p. MiR-216a-5p*-CEDS increased the expression of miR-26a-5p and miR-126-5p. MiR-365a-3p*-CEDS was observed to increase the expression of miR-216a-5p while concomitantly decreasing the expression of miR-26a-5p and miR-126-5p. And miR-655-3p*-CEDS was found to increase the expression of miR-365a-3p while concomitantly decreasing the expression of miR-21-5p.

In this study, the 12A sequence was employed as a control sequence in CEDS experiments to be compared with other specific motif sequences, including those of nine oncogenesis-related miRNAs, the p53 binding site, and the E2Fs binding site. However, the 12A sequence may exhibit homology to numerous poly-A dsDNA sequences (4A-12A) within the genome DNAs of RAW 264.7 cells. Consequently, the 12A*-CEDS is also capable of influencing gene transcription and protein expression by targeting nA (n = 4∼12) sequences within genomic DNA and miRNAs, respectively, which are almost impossible to be identified. It is therefore proposed that the 12A*-CEDS be considered a nonspecific positive control in comparison to the other CEDSs performed in this study.

CEDS has been designed to stimulate the DNA base pair polarities of target dsDNA, leading to a change in the DNA conformation. This enables CEDS to exert a regulatory effect on the target dsDNA safely. Therefore, it is suggested that CEDS can influence the target dsDNA in a sequence-specific manner with minimum biohazard, and that its results should be assessed by the probability of targeting, rather than by the mechanical targeting. Although CEDS is a non-invasive method that requires no additional medication and is relatively easy to use with a variety of procedures for multiple CEDS treatments to target specific miRNAs and DNA motifs in cells with minimal electromagnetic exposure, CEDS showed variable targeting efficiency depending on the context and cell types and also showed the secondary effect on other gene via sequence homology or feedback loop of gene transcription and protein translation. Therefore, it is highly recommended that the effect of CEDS should be explored through *in vitro* culture experiments before application in animals and patients, using the hydrogen bonding magnetic resonance (HBMR)-based gene regulation (HBMR-GR) system consisting of CEDS device, IP-HPLC analysis, and big data for gene targeting.

## Acknowledgments

We would like to express our gratitude to the late Professor Je Geun Chi and the late Dr. Soo Il Chung, who contributed to this research in part.

## References

1 Lee, S. K., Lee, D. G. & Kim, Y. S. Development of Hydrogen Bonding Magnetic Reaction-based Gene Regulation through Cyclic Electromagnetic DNA Simulation in Double-Stranded DNA. *arXiv*, arXiv:1210.7091 (2024).

2 Komatsu, S., Kitai, H. & Suzuki, H. I. Network Regulation of microRNA Biogenesis and Target Interaction. Cells 12, doi:10.3390/cells12020306 (2023).

3 Kim, S. M., Eo, M. Y., Cho, Y. J., Kim, Y. S. & Lee, S. K. Differential protein expression in the secretory fluids of maxillary sinusitis and maxillary retention cyst. European archives of oto-rhino-laryngology : official journal of the European Federation of Oto-Rhino-Laryngological Societies 274, 215–222, doi:10.1007/s00405-016-4167-2 (2017).

4 Kim, S. M., Eo, M. Y., Cho, Y. J., Kim, Y. S. & Lee, S. K. Immunoprecipitation high performance liquid chromatographic analysis of healing process in chronic suppurative osteomyelitis of the jaw. Journal of cranio-maxillo-facial surgery : official publication of the European Association for Cranio-Maxillo-Facial Surgery 46, 119–127, doi:10.1016/j.jcms.2017.10.017 (2018).

5 Lee, I. S. et al. Effects of 4-Hexylresorcinol on Craniofacial Growth in Rats. International journal of molecular sciences 22, doi:10.3390/ijms22168935 (2021).

6 Seo, M. H., Kim, D. W., Kim, Y. S. & Lee, S. K. Pentoxifylline-induced protein expression change in RAW 264.7 cells as determined by immunoprecipitation-based high performance liquid chromatography. PloS one 17, e0261797, doi:10.1371/journal.pone.0261797 (2022).

7 Yoon, C. S., Kim, M. K., Kim, Y. S. & Lee, S. K. In vivo protein expression changes in mouse livers treated with dialyzed coffee extract as determined by IP-HPLC. Maxillofacial plastic and reconstructive surgery 40, 44, doi:10.1186/s40902-018-0183-z (2018).

8 Yoon, J. H., Kim, D. W., Lee, S. K. & Kim, S. G. Effects of 4-hexylresorcinol administration on the submandibular glands in a growing rat model. Head & face medicine 18, 16, doi:10.1186/s13005-022-00320-7 (2022).

9 Dharap, A., Pokrzywa, C., Murali, S., Pandi, G. & Vemuganti, R. MicroRNA miR-324-3p induces promoter-mediated expression of RelA gene. PloS one 8, e79467, doi:10.1371/journal.pone.0079467 (2013).

10 Xiao, M. et al. MicroRNAs activate gene transcription epigenetically as an enhancer trigger. RNA biology 14, 1326–1334, doi:10.1080/15476286.2015.1112487 (2017).

11 Friedman, R. C., Farh, K. K., Burge, C. B. & Bartel, D. P. Most mammalian mRNAs are conserved targets of microRNAs. Genome research 19, 92–105, doi:10.1101/gr.082701.108 (2009).

12 Bao, F., LoVerso, P. R., Fisk, J. N., Zhurkin, V. B. & Cui, F. p53 binding sites in normal and cancer cells are characterized by distinct chromatin context. Cell cycle 16, 2073–2085, doi:10.1080/15384101.2017.1361064 (2017).

13 Carson, L. M. & Flynn, R. L. Highlighting vulnerabilities in the alternative lengthening of telomeres pathway. Current opinion in pharmacology 70, 102380, doi:10.1016/j.coph.2023.102380 (2023).

14 Gong, H., Lu, F., Zeng, X. & Bai, Q. E2F transcription factor 1 (E2F1) enhances the proliferation, invasion and EMT of trophoblast cells by binding to Zinc Finger E-Box Binding Homeobox 1 (ZEB1). Bioengineered 13, 2360–2370, doi:10.1080/21655979.2021.2023793 (2022).

15 Pan, Y., van der Watt, P. J. & Kay, S. A. E-box binding transcription factors in cancer. Frontiers in oncology 13, 1223208, doi:10.3389/fonc.2023.1223208 (2023).

16 Taciak, B. et al. Evaluation of phenotypic and functional stability of RAW 264.7 cell line through serial passages. PloS one 13, e0198943, doi:10.1371/journal.pone.0198943 (2018).

17 Hashimoto, Y., Akiyama, Y. & Yuasa, Y. Multiple-to-multiple relationships between microRNAs and target genes in gastric cancer. PloS one 8, e62589, doi:10.1371/journal.pone.0062589 (2013).

18 Park, M. H. & Wolff, E. C. Hypusine, a polyamine-derived amino acid critical for eukaryotic translation. The Journal of biological chemistry 293, 18710–18718, doi:10.1074/jbc.TM118.003341 (2018).

19 Chung, S. I. Comparative studies on tissue transglutaminase and factor XIII. Annals of the New York Academy of Sciences 202, 240–255, doi:10.1111/j.1749-6632.1972.tb16338.x (1972).

20 Chung, S. I., Shrager, R. I. & Folk, J. E. Mechanism of action of guinea pig liver transglutaminase. VII. Chemical and stereochemical aspects of substrate binding and catalysis. The Journal of biological chemistry 245, 6424–6435 (1970).

21 Lim, J. H., Latysheva, N. S., Iggo, R. D. & Barker, D. Cluster Analysis of p53 Binding Site Sequences Reveals Subsets with Different Functions. Cancer informatics 15, 199–209, doi:10.4137/CIN.S39968 (2016).

22 Morgunova, E. et al. Structural insights into the DNA-binding specificity of E2F family transcription factors. Nature communications 6, 10050, doi:10.1038/ncomms10050 (2015).

